# Whole genome sequencing of 2,023 colorectal cancers reveals mutational landscapes, new driver genes and immune interactions

**DOI:** 10.1101/2022.11.16.515599

**Authors:** Alex J. Cornish, Andreas J. Gruber, Ben Kinnersley, Daniel Chubb, Anna Frangou, Giulio Caravagna, Boris Noyvert, Eszter Lakatos, Henry M. Wood, Claudia Arnedo-Pac, Richard Culliford, Jacob Househam, William Cross, Amit Sud, Philip Law, Maire Ni Leathlobhair, Aliah Hawari, Steve Thorn, Kitty Sherwood, Güler Gül, Juan Fernandez-Tajes, Luis Zapata, Ludmil B. Alexandrov, Nirupa Murugaesu, Alona Sosinsky, Jonathan Mitchell, Genomics England Research Consortium, Nuria Lopez-Bigas, Philip Quirke, David N Church, Ian P.M. Tomlinson, Andrea Sottoriva, Trevor A. Graham, David C. Wedge, Richard S. Houlston

## Abstract

To characterise the somatic alterations in colorectal cancer (CRC), we conducted whole-genome sequencing analysis of 2,023 tumours. We provide the most detailed high-resolution map to date of somatic mutations in CRC, and demonstrate associations with clinicopathological features, in particular location in the large bowel. We refined the mutational processes and signatures acting in colorectal tumorigenesis. In analyses across the sample set or restricted to molecular subtypes, we identified 185 CRC driver genes, of which 117 were previously unreported. New drivers acted in various molecular pathways, including Wnt (*CTNND1, AXIN1, TCF3*), TGF-β/BMP (*TGFBR1*) and MAP kinase (*RASGRF1, RASA1, RAF1*, and several MAP2K and MAP3K loc*i*). Non-coding drivers included intronic neo-splice site alterations in *APC* and *SMAD4*. Whilst there was evidence of an excess of mutations in functionally active regions of the non-coding genome, no specific drivers were called with high confidence. Novel recurrent copy number changes included deletions of *PIK3R1* and *PWRN1*, as well as amplification of *CCND3* and *NEDD9*. Putative driver structural variants included *BRD4* and *SOX9* regulatory elements, and *ACVR2A* and *ANKRD11* hotspot deletions. The frequencies of many driver mutations, including somatic Wnt and Ras pathway variants, showed a gradient along the colorectum. The Pks-pathogenic *E. coli* signature and *TP53* mutations were primarily associated with rectal cancer. A set of unreported immune escape driver genes was found, primarily in hypermutated CRCs, most of which showed evidence of genetic evasion of the anti-cancer immune response. About 25% of cancers had a potentially actionable mutation for a known therapy. Thirty-three of the new driver genes were predicted to be essential, 17 possessed a druggable structure, and nine had a bioactive compound available. Our findings provide further insight into the genetics and biology of CRC, especially tumour subtypes defined by genomic instability or clinicopathological features.

## INTRODUCTION

Colorectal cancer (CRC) ranks as the second most lethal and third commonest malignancy worldwide ^1^, with cases predicted to increase from 1.9 million in 2020 to 3.1 million in 2040. Anatomical location of cancer within the colorectum is reflected in differences in aetiopathology, treatment and prognosis. CRCs are typically divided using mutational criteria into three groups. Two hypermutant groups are denoted as *MSI* (microsatellite instability-positive) caused by defective mismatch repair, and *POL* caused by *POLE* mutations that impair polymerase proofreading. The remaining largest group of CRCs are microsatellite stable (MSS) and have varying degrees of chromosomal instability (CIN) ^2^. Several driver genes and their pathways are recognised as important in the development of CRC, including Wnt, Ras-Raf-Mek-Erk, PI3-kinase, TGF-β and p53 ^2^. While large-scale sequencing projects have identified recurrent base pair-level mutations and chromosomal translocations, analyses to date have been based primarily on either whole-exome or panel sequencing of primary tumours ^2–5^. These studies have therefore given only a partial description of the CRC genome.

Herein we provide the largest and most comprehensive analysis of the genetic landscape of CRC to date, based on whole genome sequencing (WGS) of 2,017 CRC patients recruited to the UK 100,000 Genomes Project^6^. The results of our study provide new insights into disease-causing small mutations, structural aberrations and copy number alterations, within both the coding and non-coding genome. Several new driver genes represent potential therapeutic targets. Our findings also reveal differences in the genetics of CRC by molecular subtype, stage of tumorigenesis, anatomical location and prior therapy.

## RESULTS

We performed WGS **(Supplementary Figs 1-4; Supplementary Tables 1-2; Methods),** on 2,023 paired cancer (~100x average depth) and normal (blood, 33x) samples from 2,017 CRC patients (median age at sampling 69 years, range 23-94; 59.4% male). Sequenced CRCs were either primary carcinomas (n=1,898), metastases from CRCs (n=122) or recurrences (n=3).

Three hundred and fifteen patients had received chemotherapy or radiotherapy for CRC before sampling. Nineteen patients, aged 26-82 years at diagnosis had a previously unreported Mendelian CRC predisposition syndrome caused by pathogenic germline mutations in Lynch Syndrome genes (7 *MSH2*, 3 *MLH1*, and 8 *MSH6*) or *POLE*.

### Mutational processes

Three hundred and sixty-four CRCs were MSI, 18 were POL, and the remaining 1,641 MSS. To examine mutational processes underlying tumorigenesis in detail, we extracted single base substitution (SBS), doublet base substitution (DBS), small insertion and deletion (ID) and copy number alteration (CNA) mutational signatures by non-negative matrix factorisation, and structural variation (SV) mutational signatures by hierarchical Dirichlet processes ^7–9^. Extracted SBS/DBS/ID signatures were broadly concordant with those previously reported in CRC ^7–9^, with no evidence of novel signatures, but the availability of a large set of whole genome sequences resulted in updates to the frequency and activity levels of several signatures in CRCs **(Fig. 1, Supplementary Table 3).** For example, the recently uncovered SBS89 (29 cancers; unknown aetiology) and SBS94 (35 cancers; unknown aetiology) signatures were present in our samples, whereas we did not detect previously reported signatures SBS30, SBS40 and ID7. In addition to the ubiquitous clock-like signatures SBS1 and SBS5 ^7–9^, MSI tumours were characterised by SBS44 (mismatch repair deficiency) and POL cancers by SBS10a/b (POLE proofreading defect) and SBS28 (unknown aetiology). In contrast MSS cancers were enriched for SBS2/13 (AID/APOBEC activity), SBS18 (reactive oxygen species damage), SBS8 and SBS93 (both unknown aetiology) ^7–9^. SBS88, reflecting PKS-pathogenic *E. coli* exposure was a feature of 115 (5.7%) cancers (all MSS)^10^. **SBS93** (unknown aetiology, typified by T>C and T>G transitions; **Supplementary Fig. 5)** correlated with ID14 activity (unknown aetiology) (Pearson’s R = 0.81; P<2.2×10^-16^), implying a common underlying cause or mechanism.

**Fig. 1:**
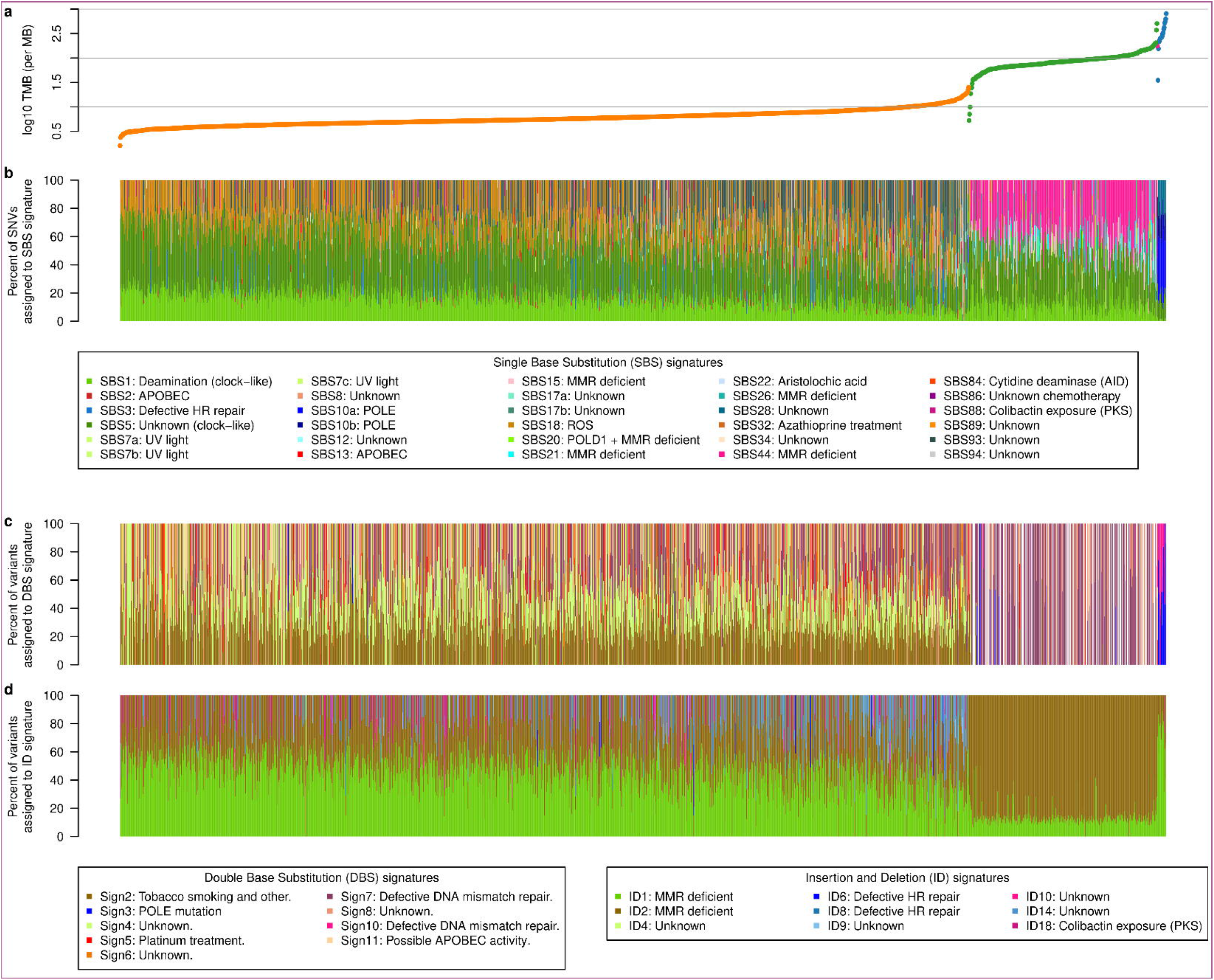
Mutational processes. **(a)** Tumour mutation burden (TMB) per megabase (MB) of microsatellite stable (MSS; n=1641; orange), microsatellite instable (MSI; n=364; green), POLE mutated MSS (POL; n=17; blue) and POLE mutated MSI (POL & MSI; n=1; magenta) tumours. Samples are first grouped according to their tumour subtype (MSS, MSI, POLE, POLE & MSI) and then ordered within each group from the sample with the lowest (left) to the sample with the highest TMB. (b) Percent of single nucleotide variants (SNV) assigned to single base substitution (SBS) mutational signatures for every sample. (c) Relative occurrence (percent) of double base substitution (DBS) mutational signatures per sample. **(d)** Percent of insertion and deletion (ID) events assigned to ID mutational signatures per sample. Samples in all panels are ordered according to the sorting described in (a) and putative artefact signatures (Supplementary Table 3) are not shown.

The 9 SV signatures extracted resembled those previously reported in CRC **(Fig. 2a, Supplementary Fig. 6)^9^.** Higher activities of SV4 (fragile site; P=6×10^-8^), SV8 (unbalanced inversions; P=6×10^-9^) and SV9 (unbalanced translocations; P=3×10^-9^)^9^ were associated with *TP53* mutation **(Fig. 2b).** SV5 (small tandem duplication) and SV6 (early replicating medium tandem duplication) activities were positively correlated with *FBXW7* mutation (P=5×10^-5^ and P=6×10^-6^ respectively), concordant with other cancers ^11^. Additionally, mutation *of ATM* correlated with SV1 (small deletions; P=1×10^-9^), consistent with its double-strand DNA break repair function^2^.

**Fig. 2:**
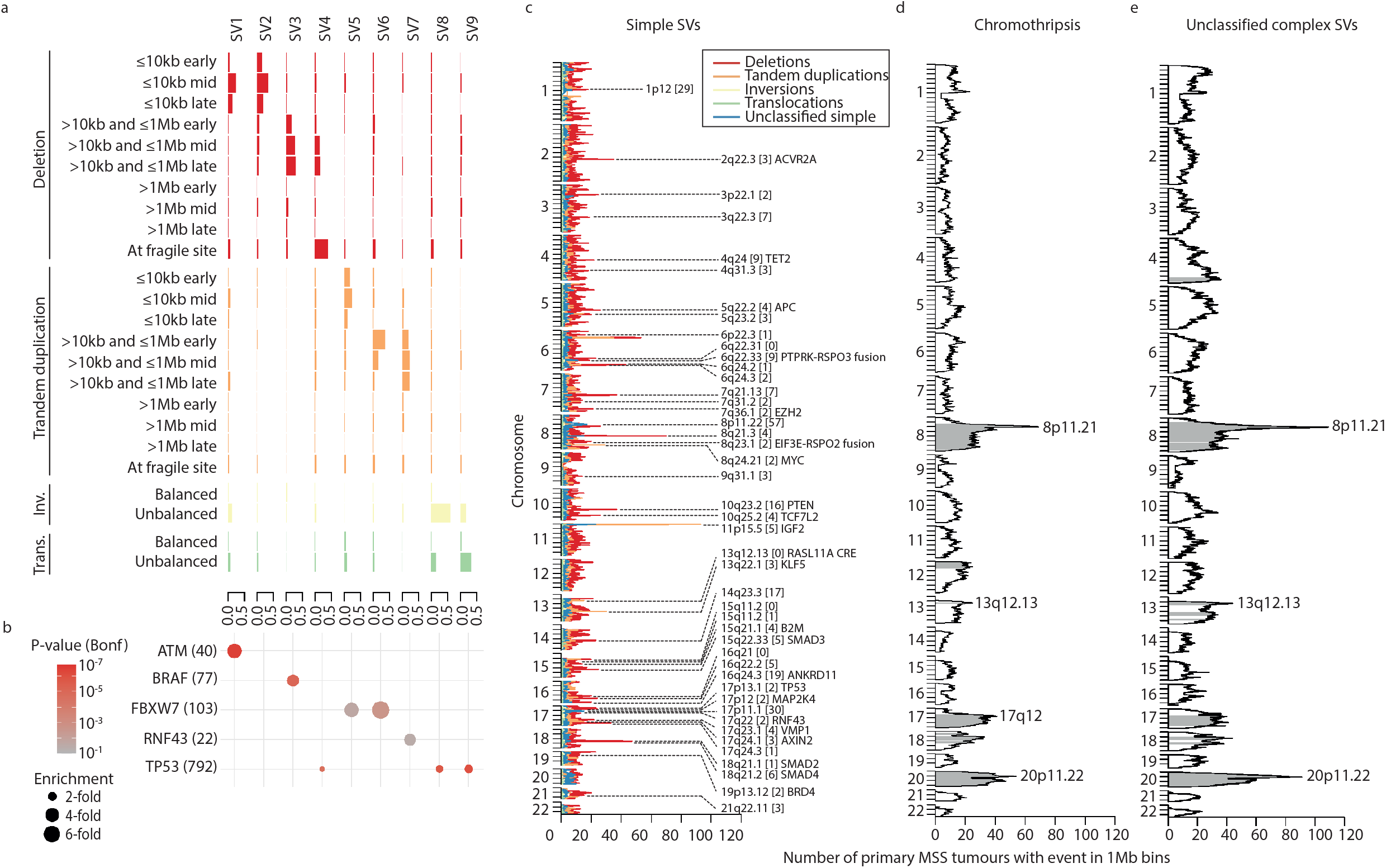
Patterns of structural variation in primary CRC. **(a)** Nine SV signatures identified in MSS, MSI and POLE-mutated CRC. Bars represent the contribution of each SV category to each signature. (b) Association of putative pathogenic germline and somatic variants in CRC driver and DNA repair genes with SV signature activity. Genes positively associated with the activity of at least one SV signature at a Bonferroni-corrected P-value <0.05 are shown. Numbers of tumours with a potentially pathogenic germline or somatic variant indicated in parentheses. Due to mutational burden heterogeneity between CRC subtypes, only primary MSS CRCs were considered. Enrichment computed as the ratio of mean number of SVs attributed to the signature in mutated and non-mutated tumours. **(c)** Significant hotspots of simple SVs identified in primary microsatellite stable (MSS) CRCs (n=1354). Non-fragile SV hotspots identified at a 5% false discovery rate (FDR) are annotated with their cytoband, the number of contained genes (in brackets) and any candidate gene. Coloured lines represent numbers of samples with an SV break point of each class in 1Mb genome regions. SVs at fragile sites have not been plotted. Y axis ticks represent 20Mb intervals. **(d)** Frequency of chromothripsis events and **(e)** frequency of unclassified complex SVs across chromosomes in primary MSS CRCs. Regions enriched for chromothripsis and unclassified complex SVs at a 5% FDR and greater than 5Mb in size are coloured blue. SVs at fragile sites have not been plotted. CRE: cis-regulatory element; inv.: inversions; trans.: translocations.

We identified six copy number signatures **(Fig. 3, Supplementary Figure 7):** CN1 (near-diploid state); CN2 (genome doubling); CN6 (chromothripsis/amplification with WGD); CN9 (CIN without WGD); CN15 (chromosomal-scale LOH); and CN18 (unknown aetiology). All except CN1, which represents a diploid state, were enriched in MSS tumours **(Supplementary Table 4).** CN18, which is associated with homologous recombination repair deficiency (HRD), has not previously been reported in CRC. The role of HRD in CRC has been debated. Using HRDetect, we predicted HRD in 12 (0.7%) cancers ^12^: three with pathogenic germline *BRCA1* mutations with loss of the wild-type allele, and three with biallelic mutation or homozygous *BRCA2* deletion **(Supplementary Fig. 8).** No cancers without HRD carried pathogenic germline *BRCA1* or *BRCA2* mutations with somatic loss. CN15 was associated with HRD deficient tumours (Fisher’s exact test P=4×10^-6^), in agreement with its previously established aetiology.

**Fig. 3:**
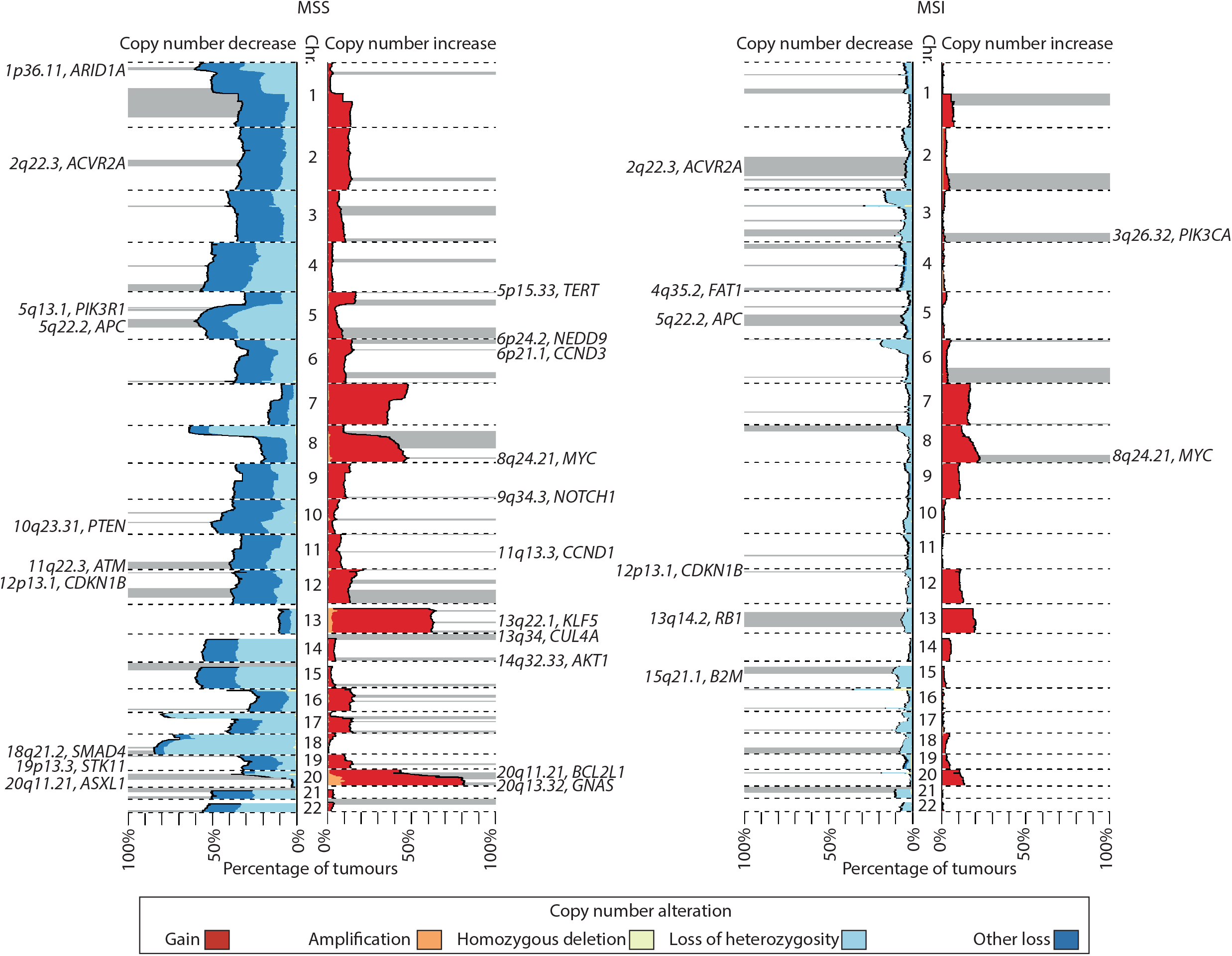
Patterns of CNA in primary CRC. Genome-wide frequency of CNA in primary microsatellite stable (MSS; n=1354) and primary microsatellite unstable (MSI; n=292) CRC. Focal amplifications and deletions reported by GISTIC are shown as grey bars and annotated with a cytoband and likely candidate gene where identified. Black dashed lines represent chromosome boundaries. Chr: chromosome.

### Coding driver mutations

Consensus driver identification using IntOGen identified a total of 185 driver genes in one or more of the three mutational subgroups. Specifically, 87, 95, 44 and 37 putative drivers were respectively identified in primary MSS, MSI and POL cancers, and metastatic MSS CRCs **(Fig. 4a; Supplementary Table** 5)^13^. These included 68 genes previously identified as CRC drivers, 51 identified as drivers in cancer types other than CRC, and 66 not previously implicated in cancer^2–4,14–18^. Primary and metastatic MSS tumours had four pathogenic driver mutations on average, whereas there were significantly elevated numbers of driver mutations in primary MSI and POL tumours, with averages of 23 and 30 respectively (Kruskal-Wallis P-value< 2×10^-16^; **Supplementary Fig. 9; Supplementary Table 6).**

**Fig. 4.**
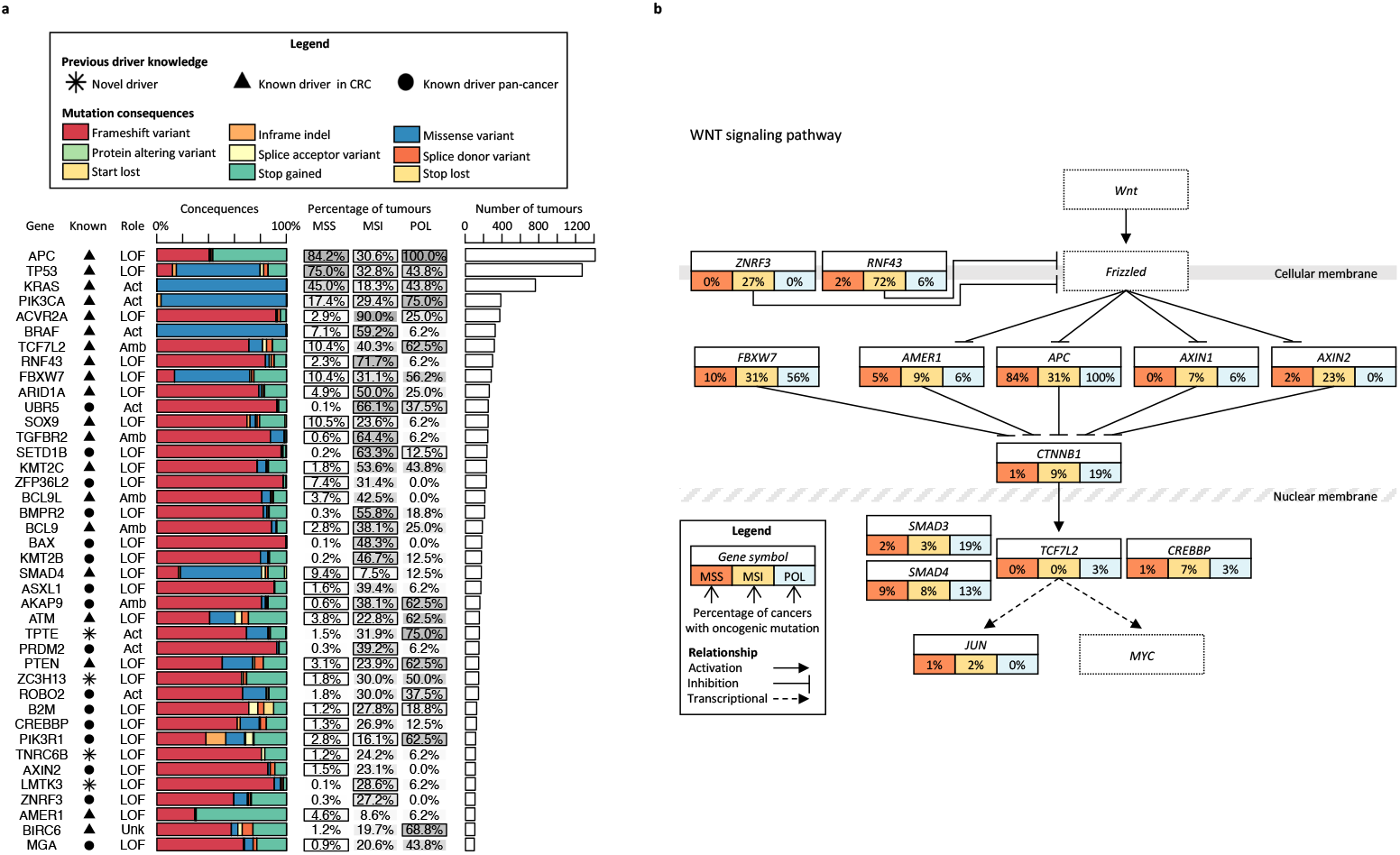
Driver genes in CRC. **(a)** 40 genes with highest oncogenic mutation frequency. A gene was considered to be a known CRC driver if it had previously been identified in CRC cohort(s)^13^, a known pan-cancer driver if it had previously been identified as a driver in only non-CRC cancer cohorts ^13,14^ and a novel driver otherwise. Mutation mechanism of action (loss of function [LOF], activating [Act], unknown [Unk] or ambiguous [Amb]) was assigned considering previously curated roles ^13^ and roles predicted by this study’s IntOGen for the primary MSS cohort, with conflicts marked as ambiguous. Pathogenic mutation consequences were predicted using VEP. Percentage of tumours in the primary microsatellite stable (MSS; n=1,521), primary microsatellite unstable (MSI; n=360) and primary POLE-mutated (POL; n=16) cohorts with an oncogenic mutation are shown with a greyscale fill corresponding to the percentage. Genes identified as CRC drivers are annotated with a black border around the corresponding cohort percentage cell. Number of samples represents the total number of tumours with an oncogenic mutation across primary MSS, MSI and POL cohorts (n=1,897). (b) Frequency of driver mutations in Wnt signalling pathway. Pathway information obtained from KEGG ^60^ and TCGA^2^. Key pathway genes not identified as CRC drivers by IntOGen are included with dashed borders. **(c)** Frequency of driver mutations in the Ras-Raf-Mek-Erk/MAP-kinase signalling pathway (as per (b).

The major known CRC driver genes were mutated at broadly expected frequencies **(Supplementary Table 7).** Among these, oncogenic *APC, TP53* and *KRAS* mutations were enriched in MSS cancers, while *PIK3CA* and *PTEN* mutations were more common in MSI and POL cancers (all P-values <5×10^-6^; **Supplementary Table 7).** In drivers detected in both the MSS and MSI sub-groups, the latter showed strong enrichment *for BRAF^V600E^* mutations and highly recurrent frameshift mutations at short repeats in *RNF43, TGFBR2, BCL9* and *ARID1A* (P<2.47×10^-32^; **Supplementary Table 8).** Both MSI and POL tumours were enriched for drivers predicted to cause immune evasion (see below). Many genes were identified as drivers in a single molecular subgroup **(Supplementary Tables 7 & 8),** including *NRAS* and *PCBP1* in MSS cancers, *BAX* and *BMPR2* in MSI, and *RB1* in POL.

The 117 genes not previously reported as CRC drivers included those with (i) low-frequency oncogenic hotspot mutations (*e.g.AKT, FGFR2, IDH1, RAF1*), (ii) recurrent variants, including missense mutations in *SMARCB1*, and the A116T *STAG1* substitution, and (iii) loss-of-function (LOF) mutations in tumour suppressors (e.g. *CDKN2A, CTNND1, KMT2B*). Exemplars are shown in **Supplementary Figs 10-12,** and **Supplementary Table 5.** In addition, multiple members of the same family were sometimes mutated (e.g. five genes encoding MAP2 or MAP3 kinases). Novel MSS-specific coding drivers were mutated in a small proportion of cancers (0.9-3.9%). They included the Ras activator *RASGRF1*, recurrently mutated in its RhoGEF domain **(Supplementary Fig. 10),** and genes in signalling pathways with LOF mutations **(Supplementary Table** 5). Among MSI cancers, the recurrent L1178fs (29% cases) and Q1202fs (25% cases) frameshift mutations in the GTPase *RGS12* are likely to enhance MAPK signalling ^19^, while mutations in multiple driver genes mediate immune evasion (see **Immune editing and escape** below).. The A30V substitution in *H3Y2* in 5% MSI cancers suggested that the gene acts as an oncohistone. Other recurrent hotspot mutations in MSI CRCs were in genes with roles in cell signalling, RNA splicing, protein ubiquitylation and apoptosis **(Supplementary Table** 5).

Using MutationTimeR ^20^ to infer the clonal/subclonal status of driver mutations 97% of primary MSS driver mutations were clonal compared with 89% in primary MSI tumours (P<2.2×10^-16^; **Supplementary Table 9).** In general, genes with higher proportions of subclonal mutations had lower overall mutation prevalence in the CRC cohort. Among primary MSS drivers with >5% mutation frequencies, only three genes – *SMAD4* (13%, 18/135), *ZFP36L2* (5.7%, 6/105) and *PIK3CA* (5.7%; 14/253) – had > 5% subclonal mutations; it is of note that two of these are well-established CRC drivers. In contrast, for primary MSI drivers, there were 92 such genes fulfilling these criteria, including *CDK12* (33.3% subclonal), *MAP3K1* (28% subclonal), and *CDKN2A* (25% subclonal), in addition to *ZFP36L2* (20% subclonal; **Supplementary Table 9).**

### Non-coding driver mutations

In primary MSS cancers, we used OncodriveFML^21^ to identify candidate driver mutations in non-coding elements (enhancers, promoters, 30bp regions around splice sites, 3’ and 5’ untranslated regions, non-coding RNAs, ENSEMBL-curated protein binding sites and open chromatin regions; **Methods).** There was evidence of an overall enrichment of mutations in functional locations, notably promoters and transcription factor binding sites **(Supplementary Fig. 13),** and 85 specific elements were also enriched for mutations (Q<0.01; **Supplementary Table 11).** These included *APC* neo-splice site mutations (Q=5×10^-5^) driven by chr5:112,815,487 A>G (c.835-8A>G) in 6.4% cancers, causing disruption of exon 9 splicing and producing a truncated protein^5^. We similarly identified enrichment of mutations in *SMAD4* splice regions (Q=5×10^-5^) driven by A>G changes at chr18:51,058,332 (c.788-A>G; 0.3% cancers), with predicted effects on *SMAD4* splicing (SpliceAI delta scores 0.7 for acceptor loss, 0.87 for acceptor gain). For both genes, the pattern of second hits suggested that the mutations acted as loss-of-function changes equivalent to canonical protein-truncating mutations **(Supplementary Table 12).**

*TERT* promoter mutations were not classed as drivers by OncodriveFML (frequency 0.9%, 14/1521 primary MSS tumours), and no association with telomere length was observed **(Supplementary Tables 10 & 11).** Significant enrichment of mutations in distal promoters of *CSMD3* (3.1%, Q=6×10^-4^) and *ST6GALNAC1* (0.9%, Q=2×10^-4^) were found. In the case of *CSMD3, we* identified two hotspots overlapping a CTCF binding site **(Supplementary Results).** Although CTCF binding site mutations have been suggested as cancer drivers^22^, our recent data suggest that these sequences accumulate passenger mutations^23^, and we are therefore very cautious about declaring the *CSMD3* changes as drivers. In line with this, we found that 57 CTCF binding sites genome-wide were enriched for mutations with predicted high functional impact, but with uncertain true effects (Q< 0.01; **Supplementary Fig. 13, Supplementary Table 11).**

### Structural variants

Simple SVs such as deletions, tandem duplications and inter-chromosomal translocations, and complex SVs such as chromothripsis, were identified adopting a consensus approach^9^. Using a simulation-based procedure, 87 and 23 simple SV hotspots were identified in separate analyses of primary MSS and MSI cancers respectively (false discovery rate<5%; **Fig. 2c; Supplementary Fig. 14; Supplementary Table 13**)^24^. Previously reported SV hotspots in primary MSS cancers included deletions of *APC, PTEN* and *SMAD4*, amplifications of *IGF2*, and a *RASL11A* regulatory element, and *EIF3E-RSPO2* and *PTPRK-RSPO3* fusion events **(Supplementary Table 13**)^2,4,18,25^. Focal *TP53* deletions observed in osteosarcoma and prostate adenocarcinoma were also found in 2.4% of primary MSS cancers ^26^. A region on 17q23.1 containing *VMP1*, previously reported in breast and pancreatic cancers, was recurrently deleted in 1.2% of primary MSS cancers. Recurrent intronic deletions at 19p13.12 included a regulatory element interacting with the *BRD4* promoter^27^.

Novel SV hotspots in primary MSS tumours included those at 2q22.3 and 16q24.3, respectively containing driver tumour suppressor genes *ACVR2A* and *ANKRD11* (focally deleted in 1.5% and 1.6%; **Supplementary Table 13**)^2^. *EZH2* (1.3%) is the most credible target of 7q31.2 deletions, given that lower *EZH2* expression has been associated with worse prognosis in CRC^28^. *TET2* (0.8%) represents a potential target of the 4q24 rearrangements, given the tumour-supressing role it plays in haematological malignancies^29^. Along with *SMAD3* and *SMAD4* deletions, we also identified previously unreported focal deletions of *SMAD2* on 18q21.1 (1.8%). We also found 17q24.3 re-arrangements that included a regulatory element interacting with the promoter of the CRC driver *SOX9* and may therefore be under selection through *SOX9* dysregulation **(Supplementary Fig. 15).**

Kinase fusions represent a unique opportunity for targeted therapies. Fusions involving the kinase domain of previously reported partner genes were identified in 0.4% and 4.1% of MSS and MSI cancers respectively (*8 NTRK1*, 6 *BRAF, 2 ALK*, 1 *NTRK3* and 1 *RET*), supporting their role in MSI CRC in particular^30^.

In primary MSS cancers, enrichments of chromothripsis and miscellaneous complex SVs were observed on chromosomes 8, 17, 18 and 20 **(Fig. 2d-e).** The non-fragile site having the greatest complex SV enrichment was chr8:28-62Mb, which contained a chromothripsis or unclassified complex SV in 9% and 18% of MSS and MSI cancers respectively. 44% of samples with arm-level 8p deletions had a complex SV at chr8:28-62Mb, compared with 12% of samples without arm-level 8p deletions, suggesting that complex SVs are often responsible for 8p arm-level deletions.

### Copy number alterations

Whole genome duplication (WGD)^31^ was identified in 45.0%, 5.8% and 10.0% of primary MSS, MSI and POL cancers respectively **(Supplementary Fig. 16).** WGD was most frequent in *TP53*-mutated primary MSS cancers (logistic regression P=2×10^-12^)^32^. No association was present between WGD and age, tumour site, stage or grade, in either primary MSS or MSI cancers. Hierarchical clustering based on copy number states genome-wide confirmed the split between WGD and non-WGD cancers, and suggested that the former might be comprised of sub-groups with higher and lower copy number change frequencies **(Supplementary Fig. 16).**

We identified all previously reported, recurrent arm-level CNAs (including whole chromosome changes), defined as events comprising >50% of total arm size^2,20^ **(Supplementary Table 14).** Arm-level increased copy number typically involved single or double copy gains, with the exception of 20q which gained 4 or more copies in 18% of primary MSS cancers **(Supplementary Fig. 17).** Whilst MSI and POL cancers were near-diploid, as expected^2^, a small number of changes were present at levels significantly above background **(Supplementary Table 14).**

We identified individual focal CNAs encompassing 3Mb or less, and also mapped minimal common regions shared between larger CNAs^33^. Focal lesions were rare in MSI and POL tumours and no recurrent changes were found. Previously reported focal CNAs in primary MSS cancers included single/double-copy gains of *CCND1, ERBB2* and *MYC, KLF5*, and deletions of *ARID1A, APC* and *SMAD4* **(Fig. 4; Supplementary Tables 15 & 16**)^2,20^. Whilst 5p15.33 (*TERT*) amplification was observed in 24.8% of MSS cancers, a link to telomerase activity could not be established and we found no association with telomere length^34^ **(Supplementary Table 15).** Novel focal CNAs identified in primary MSS cancers are shown in **Supplementary Table 15** and included the following: 5q13.1 deletions (29.3%) containing the driver gene *PIK3R1*; 15q11.2 deletions (42.1%) containing the lncRNA *PWRN1*, an established tumour suppressor in gastric cancer^14^; and amplifications at 6p21.1 (27.9%) and 6p25.3 (25.3%), which may respectively target *CCND3* and *NEDD9*, genes that we also identified as CRC drivers based on analysis of small-scale mutations **(Supplementary Table 5**).

### Extra-chromosomal DNA

Candidate extra-chromosomal DNA (ecDNA) molecules were detected and classified using Amplicon Architect^35^ to distinguish between truly circular ecDNA molecules and those characterised by breakage-fusion-bridge cycles (BFB), representing more complex events as well as being consistent with linear amplification. There were differences in CRC subtype by ecDNA content, with 28% (380/1,354) of primary MSS tumours containing at least one circular ecDNA, compared with 36% (38/105) metastatic MSS, 1.4% (4/292) primary MSI, and 0% (0/10) primary POL tumours (*P*< 0.001, primary MSS v MSI; **Supplementary Fig. 18; Supplementary Table 17).** Primary MSS tumours containing circularised ecDNA were significantly more likely to exhibit chromothripsis, consistent with previous reports^36^ (*P_Fisher_*=1.09×10^-12^; OR=2.43; **Supplementary Table 17).** We further explored the extent to which circularised ecDNA may mechanistically underpin the amplification of known amplification by considering the ecDNA classification of “big gains” (that is, total copy number ≥ 5 in diploid tumours and ≥ 10 in tetraploid tumours). In primary MSS tumours, only 5% of oncogene “big gains” (34/665) encompassed circular ecDNA molecules, consistent with a relatively modest role for circularised ecDNA-mediated oncogene amplification in CRC compared to other cancer types **(Supplementary Fig. 18; Supplementary Table 17).**

### Mitochondrial mutations

We searched for somatic mitochondrial mutations in CRCs **(Supplementary Fig. 19).** The distribution of C>T substitutions on heavy and light mitochondrial DNA strands supported the theory that processes driving mtDNA mutation in CRC are replication rather than transcription coupled ^30^. In the set of primary MSS cancers, there was evidence for positive selection of several somatic changes, notably missense and truncating mutations in *MT-CYB* (encoding cytochrome B) and missense mutations in the NADH dehydrogenase genes *MT-ND1, MT-ND2, MT-ND3, MT-ND4* and *MT-ND4L* (Q<0.05; **Supplementary Table 18).**

The importance of mitochondria to colorectal tumorigenesis was emphasised by the identification of nuclear genes with mitochondrial functions as CRC drivers. DNA polymerase *POLG* was identified as a driver gene in the MSI cancer subtype and this may lead to mitochondrial hypermutation. However, mtDNA mutational burden did not differ significantly between *POLG*-mutated and -wildtype cancers (Wilcoxon test, P=0.417, n=65). Another novel nucleus-encoded driver was *CLUH*, which plays a role in the transport of nucleus-encoded proteins to the mitochondrion.

### Pathways of tumorigenesis

Integrated analysis of all types of coding and non-coding mutations allowed us to obtain a better understanding of which molecular pathways must be disrupted for colorectal carcinogenesis, and which are non-essential, yet contribute to tumorigenesis^37^. We grouped primary cancers by MSI, MSS and POL status and identified recurrent alterations in Wnt, Ras-Raf-Mek-Erk/MAP-kinase, PI3-kinase, TGF-β/BMP and p53 signalling pathways. While the Wnt-signalling pathway was disrupted in almost all cancers, there were differences in driver genes and mutations between MSS, MSI and POL cancers **(Fig. 4b).** Disruption of the other major pathways typically occurred in 20-50% of cancers, with similar heterogeneity between MSS, MSI and POL groups **(Fig. 4c** for exemplar of MAP-kinase pathway).

After taking account of total tumour SNV and small indel mutation burdens (TMBs), in primary MSS cancers we identified just 4 gene pairs for which alterations (mutation, amplification, or deletion) co-occurred more than expected, and 15 gene pairs in which alterations co-occurred less than expected **(Supplementary Table 19).** Previously reported relationships included co-occurrence of *SMAD2-SMAD3* and *BRAF-RNF43* mutations, and tendency towards mutual exclusivity of *KRAS-NRAS* and *KRAS-TP53* mutations (Q<0.05; **Supplementary Table 19**)^2^. Previously unreported relationships included the co-occurrence of *PCBP1-PIK3CA* mutations (Q=0.039). *PCBP1* knockdown activates *AKT*^38^, whilst *PIK3CA* positively regulates *AKT*, suggesting a possible synergistic effect. *PCBP1* mutation was negatively associated with *TP53* mutation, which itself also regulates *AKT*^39^. *KRAS* and *PIK3CA* mutations were negatively associated with *SRSF6* and *TOP1* mutation or amplification (Q<0.05). A weaker negative association was also identified between *SRSF6* and *TCF7L2* (Q=0.014).

### Immune editing and escape

Several of the genes identified as drivers, especially in the MSI and POL CRCs, have a putative role in escape from the anti-cancer immune response, and the diversity of changes implies that there are multiple mechanisms by which this is achieved **(Supplementary Table** 5). Examples include mutations at several HLA loci (including copy number changes), *TAP1* and *TAP2* antigen processing transporters, the *CCR7* and *IL7R* chemokine receptors, interferon regulators *IRF1* and *NLRC5*, the lymphocyte interaction ligand *CD58*, and *LCP1* which is involved in T cell activation.

Predicted neoantigen burden in each CRC was highly correlated with total mutational burden, as expected (Pearson R=0.89, P<10^-16^; **Fig. 5a**) ^40^. Among recurrent non-synonymous mutations, *KRAS* G12V was the most antigenic, predicted to bind patient-specific HLA molecules in 80% (146/181) of mutation-carrying cancers **(Supplementary Fig. 20a; Supplementary Methods).** *KRAS* G12D and G13D were also frequently predicted to be antigenic, whereas the less common *KRAS* mutations G12C, A146T and G12A were less likely to give rise to neoantigens. *BRAF* V600E, the overall most individually recurrent mutation, was predicted to be antigenic in 36% of cancers (98/272), as the resulting epitope was predicted to bind uncommon HLA alleles that were also prone to be lost following common HLA alterations (20% of cancers with epitope binding).

**Fig. 5:**
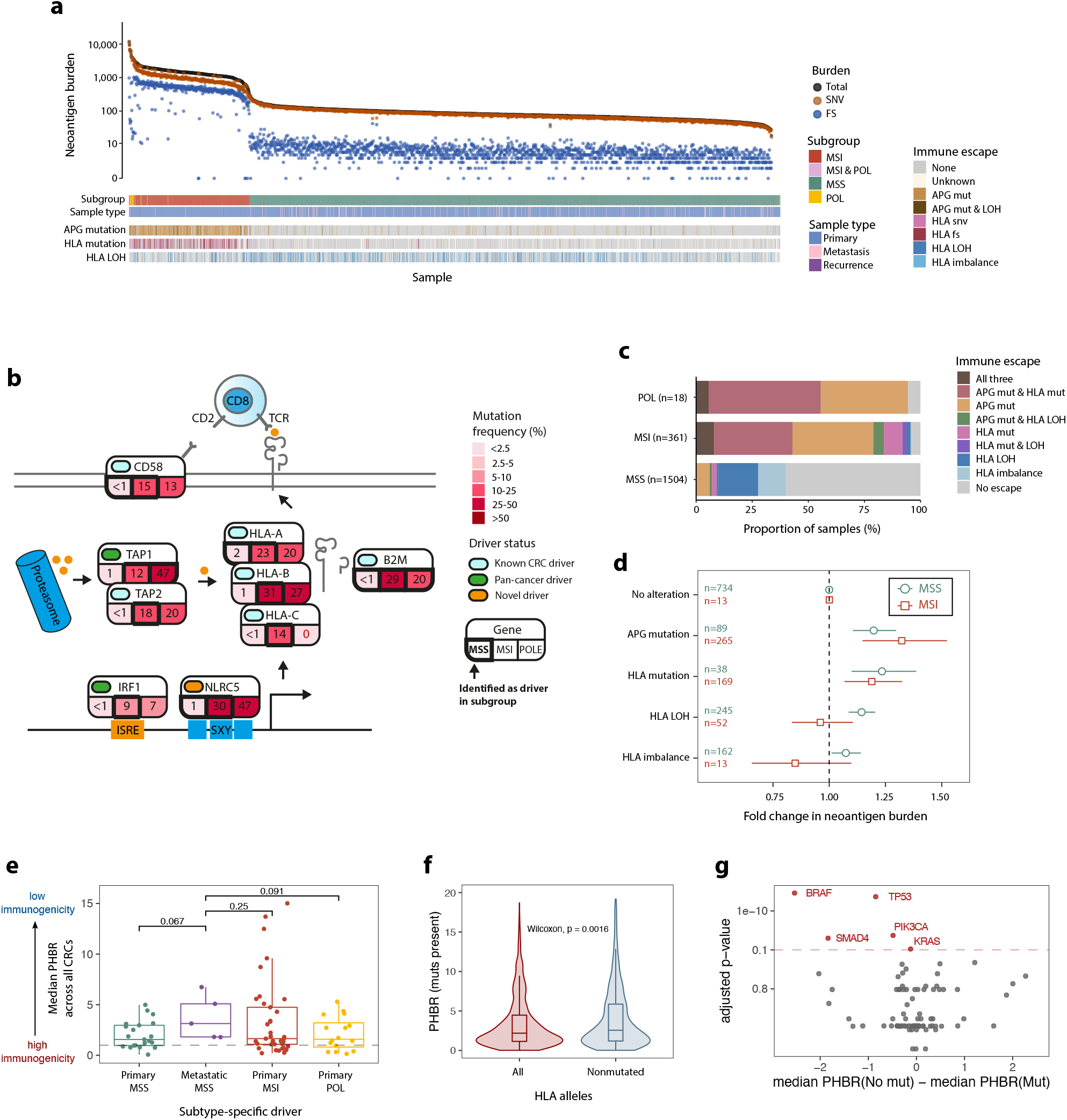
Immune landscape of CRC. **(a)** Neoantigen burden and immune escape mutations of cancers. Bottom panel shows APG and HLA alterations present in each cancer. **(b)** Proportion of cancers with an APG mutation, HLA mutation, HLA LOH, or a combination of these, suggesting immune escape. 140 cancers were excluded from the analysis due to incompatible HLA types. **(c)** Effect of immune-escape-associated alterations on neoantigen burden (with no alteration as baseline) in MSS and MSI cancers derived via multivariate regression analysis. Whiskers show 95% confidence intervals. **(d)** Immunogenicity (median PHBR in all CRCs) of cohort-specific driver mutations. Dashed line denotes substantially immunogenic mutations (PHBR=1). **(e)** The influence of HLA alterations on PHBR in patients that carry an alteration. Values computed for each driver in each HLA-mutated patient using full set of patient-specific HLA alleles (in red) are compared to values computed from a reduced, non-mutated set (in blue). **(f)** Median PHBR difference of nonmutated and mutation-carrier patients for driver genes. Each dot denotes a driver, genes with significant difference (Q<0.1) are highlighted in red.

The most common 20 peptide-changing frameshift mutations, most frequent in MSI cancers in which they were typically present in >40% of cases, showed high immunogenic potential, producing a neoantigen in >95% of cases **(Supplementary Fig. 20b).** *ACVR2A* K437fs was an outlier, as it was antigenic in only 57% of cancers in which it occurred due to a stop codon very soon after the indel. The most frequent protein-changing frameshift mutation across MSS cancers, *APC E*1309fs, was rarely antigenic, with a binding epitope predicted for only 14/47 (30%) cases. Overall, for the 20 most frequent non-synonymous and 20 most frequent frameshift changes, there was a negative association between observed mutation frequency and antigenic frequency (proportion of mutated cancers where the mutation was antigenic) for non-synonymous mutations (P=0.042), and no association for frameshifts (P=0.32).

To evaluate the immunogenic potential of common (driver) mutations in all cancers, whether or not actually present in that cancer, we used the Patient Harmonic Best Rank (PHBR) scores of Marty *et al.* ^41^. PHBR quantifies the potential of a mutation to produce an HLA-binding novel epitope depending on the patient’s HLA haplotypes **(see Supplementary Methods**) – high PHBR reflects low binding and immunogenicity, while low PHBR (<1) denotes strong immunogenic potential. We confirmed in CRC specifically the previous pancancer observations that most common driver mutations are of low immunogenic potential **(Supplementary Fig. 21a).** We also found that drivers that were only identified as such in metastatic MSS cancers had lower immunogenic potential, with no mutations falling below a median PHBR of 1 (median for primary MSS=1.56, median for metastatic MSS=3.13, P=0.067, **Fig. 5b);** and shared driver mutations were slightly more immunogenic in MSI+ cancers **(Supplementary Fig. 21b).** Driver mutations were also found to be restricted by patient-specific immunogenicity, as they were significantly enriched in patients where they were associated **with** a low immunogenic **potential_(Supplementary Fig. 22)** In addition, loss of HLA alleles via mutation or LOH led to a significant decrease in immunogenicity of the driver mutations present **(Fig. 5c).** Differential immunogenicity analysis (comparing the immunogenic potential between mutated and wild-type cancers for each driver mutation) revealed four driver genes (*BRAF, TP53, SMAD4* and *PIK3CA*) that were significantly enriched for mutations in patients where they had reduced immunogenic potential (Q<0.1, **Fig. 5d).** *KRAS* also showed a significant enrichment but the associated difference in immunogenicity was negligible, in line with the observation that the most common *KRAS* mutation is also more antigenic, suggesting that in this case, direct positive selection outweighs immunogenicity. Altogether, these observations confirm that immune editing plays a role in defining the driver landscape of CRCs.

Patterns and prevalence of immune escape differed by CRC subtype **(Fig. 5e)** in agreement with other studies ^4,42–44^. We separately evaluated allelic imbalance, loss-of-heterozygosity and protein-altering mutations in the HLA-A/B/C (MHC type-I) genes and somatic mutations in a core set of other antigen-presenting genes (APGs: *PSME3, PSME1, ERAP2, TAP2, ERAP1, HSPBP1, PDIA3, CALR, B2M, PSME2, PSMA7, IRF1, CANX, TAP1, CIITA*). Multivariate regression analysis revealed that all mechanisms of immune escape were associated with higher neoantigen burden in MSS cancers (P< 0.001, **Fig. 5f).** HLA (type-I) mutation had the strongest effect, with cancers carrying an HLA mutation having a predicted 21% increase in burden compared to HLA wild-type cancers (P=0.001). Conversely in MSI cancers, only mutations of HLA and other APGs were associated with higher neoantigen burden (P=0.002 and P=10^-5^), with APG mutation corresponding to a 35% neoantigen burden increase. Immune escape resulting from any of the above mechanisms remained a significant determinant of neoantigen burden in multivariate regression (P=0.012), even after accounting for clinical characteristics and TMB **(Supplementary Fig. 23).** Tumour site in MSS cancers was also a significant determinant (P< 0.001), cancers from the distal colon and rectum showing a lower neoantigen burden than right-sided cancers. In MSI cancers, on the other hand, prior treatment (N=34) was associated with an increased neoantigen burden independent of overall TMB (P=0.006), potentially linked to the genetic immune escape detected in 33/34 of treated MSI cancers **(Supplementary Table 7).**

### Mutations induced by prior therapy

Colorectum-targeted radiotherapy (RT), which is generally applied in the neoadjuvant setting, was associated with higher ID8 signature activity, as previously reported ^45^. This finding is consistent with conversion of those tumours by selection or drift to an oligoclonal state after treatment (P=5×10^-8^, **Fig. 6a; Supplementary Table 20).** ID8 activity was not observed in 31 metastases with prior colorectum-targeted RT **(Fig. 6e)** consistent with secondary spread occurring prior to RT. Neoadjuvant oxaliplatin treatment was associated with higher DBS5 activity (P=8×10^-5^; **Fig. 6b**) ^46^, although other signatures previously associated with platinum-based chemotherapy, including SBS31, were not observed ^47^. DBS5 activity was observed in both primary tumours and metastases **(Fig. 6f).** For both RT and oxaliplatin, treatment duration (P=0.004, **Fig. 6c;** P=0.006; **Fig. 6d)** and time since treatment (P=2×10^-6^; **Fig. 6e;** P=1×10^-3^; **Fig. 6f)** correlated with their respective signature activity. Of note, signature SBS17b, associated with 5FU therapy^48^, was not detected in pre-treated cancers.

**Fig. 6.**
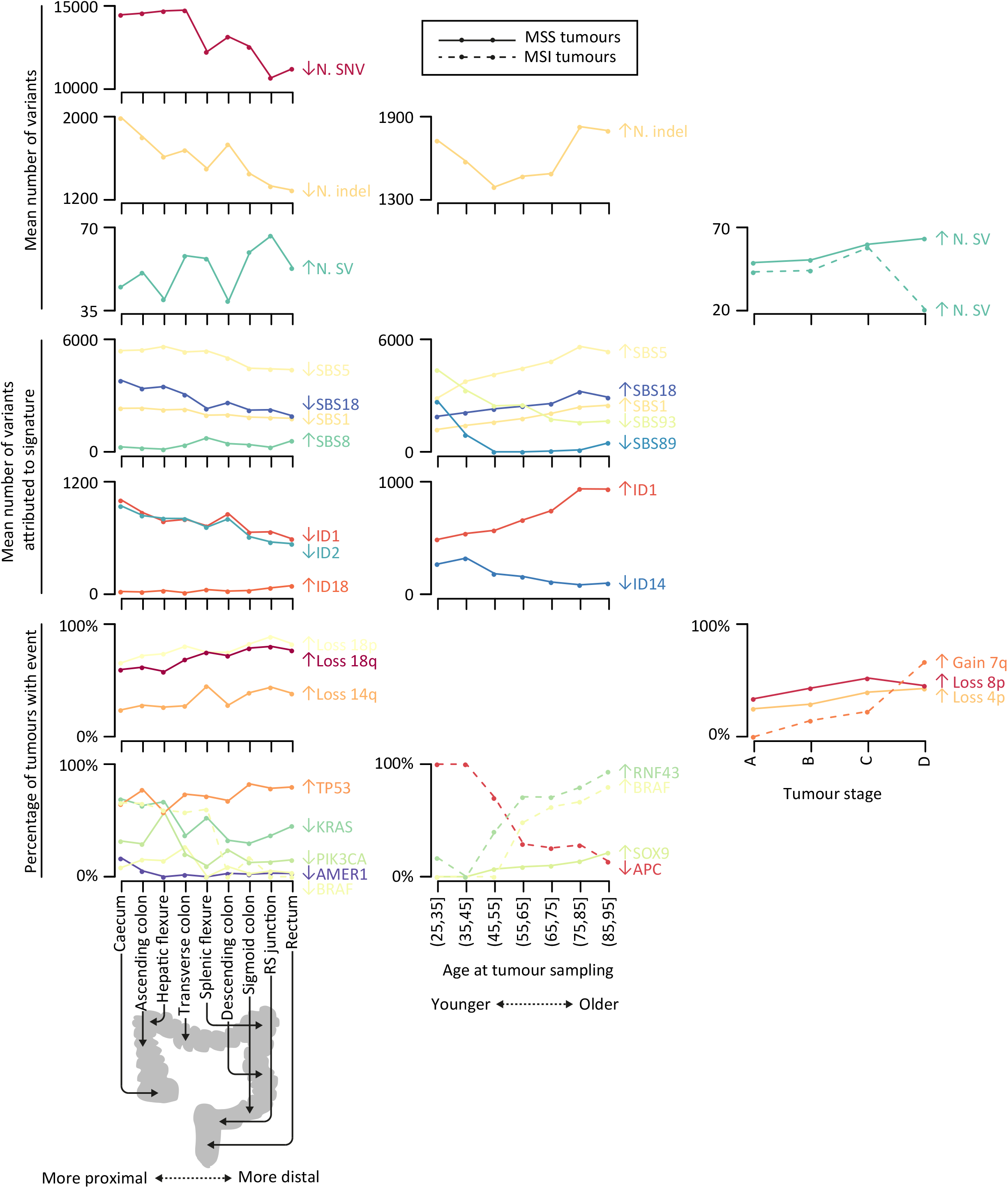
Genomic impact of systemic treatment and radiotherapy. **(a, b)** Proportion of tumours with evidence of ID8 and DBS5 by prior treatment status. n represents the number of tumours in each group. **(c, d)** Numbers of mutations attributed to ID8 and DBS5 for tumours with different prior radiotherapy and oxaliplatin treatment duration lengths. Length of treatment duration measured as the number of days between first and last treatment. Tumours divided into quartiles of equal size. **(e, f)** Proportion of tumours with evidence of ID8 and DBS5 by time since first radiotherapy and oxaliplatin treatment. Grey: primary tumours, black: metastases. (g) ID8 activity in tumours from participants with prior radiotherapy for various cancers.

RT for prostate cancer is associated with higher risk of second malignancies, including CRC ^49^. Twenty-seven participants with RT-naive CRC had previously been treated for prostate cancer by RT and in these ID8 activity was raised (P=3×10^-13^; **Fig. 6g),** consistent with induction of mutations in CRC progenitors by treatment.

### Clinicopathological correlates of primary cancers

We initially considered molecular differences associated with tumour location in the colorectum. We focussed on the primary MSS cancer set and treated location as a continuous variable from caecum (most proximal) to rectum (most distal) **(Fig. 7, Supplementary Table 21).** Whilst we found no associations with tumour grade or patient sex, CRCs originating from more distal sites had a greater number of SVs (P=4×10^-4^), but fewer SNVs (P=6×10^-16^) and indels (P=6×10^-22^; **Fig. 7; Supplementary Table 22).** Higher SBS8 and ID18 activity, and lower SBS1, SBS5, SBS18, ID1 and ID2 activity, was also observed in cancers from more distal sites (P<0.05; **Supplementary Fig. 24)** and it is of note that published data ^50^ indicate lower SBS1, SBS5, SBS18, ID1 and ID2 activity in normal epithelial cells from the distal than the proximal colon (P<0.05; **Supplementary Fig. 25).** Higher ID18 activity in distal CRCs suggested that PKS-pathogenic *E. coli* (see below) exert a greater mutagenic burden at these sites ^10^, consistent with reports that mutational signature IDA, closely resembling ID18, exhibits higher activity in normal epithelial cells from the distal colon **(Supplementary Fig. 25**) ^46^. Distal primary MSS CRCs were typified by higher frequencies of *TP53*, and lower frequencies of *AMER1, BRAF, KRAS* and *PIK3CA* driver mutations (P<0.05; **Fig. 7; Supplementary Table 23**)^5^. Arm-level deletions of 14q, 18p and 18q also occurred more frequently in these cancers, as did focal deletions of 1p36.11, 18q21.2, and 18q22.3, and 20q13.33 gain (P<0.05; **Supplementary Table 24).**

**Fig. 7.**
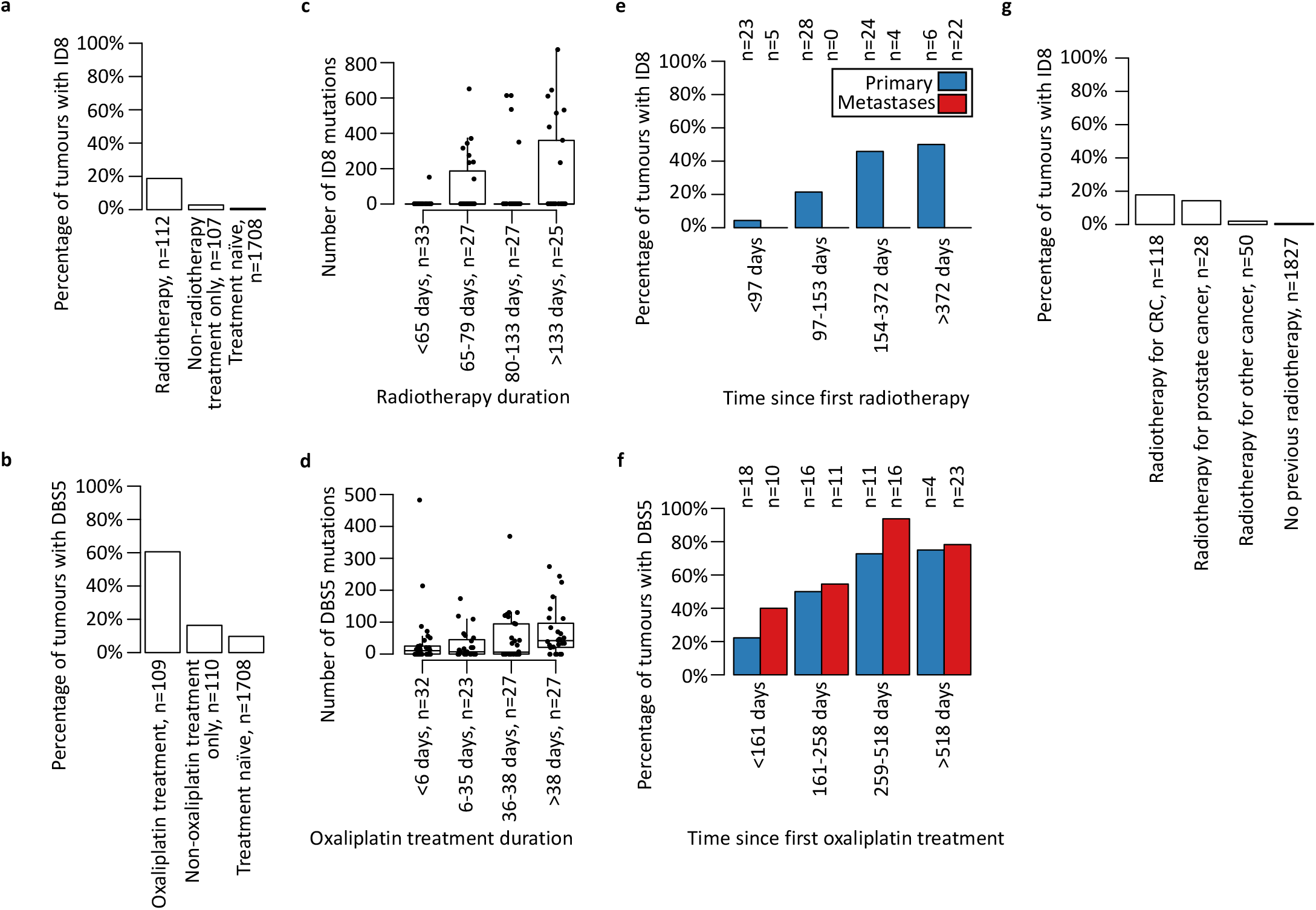
Correlation of primary tumour site and age at sampling with genomic features. Shown are total numbers of variants, numbers of variants attributed to mutational signatures, arm-level copy number alterations and driver gene mutations associated with primary tumour site or age at tumour sampling in multiple regression analyses. Only significant associations identified in primary microsatellite stable (MSS) or microsatellite unstable (MSI) tumours are included (Bonferroni-corrected P<0.05). Up- and down-facing arrows indicate whether primary tumour site (from caecum to rectum) or age at tumour sampling were positively or negatively correlated with the genomic feature. Bottom figures show percentage of tumours with a predicted-oncogenic SNV or indel driver gene mutation. SNV: single nucleotide variant; SV: structural variant; RS: rectosigmoid.

Patient age was positively associated with SBS1, SBS5, SBS18 and ID1 activities, consistent with SBS18 (reactive oxygen species damage) being a clock-like signature (P<0.05; **Fig. 7; Supplementary Fig. 26; Supplementary Table 22)^7,51^.** Conversely, SBS89, SBS93 and ID14 activities were negatively correlated with age at sampling (P<0.05). Whilst SBS89 has unknown aetiology, it has been reported to occur in normal colon tissue during the first decade of life ^50^. Age at sampling also correlated with higher *SOX9* oncogenic mutation frequency in primary MSS cancers, and higher *BRAF* and *RNF43*, and lower *APC* pathogenic mutation frequency in primary MSI cancers (P<0.05; **Supplementary Table 23).**

Tumour stage was associated with higher frequencies of arm-level 4p and 8p deletions in primary MSS cancers, and 7q gains in primary MSI cancers (P<0.05; **Supplementary Table 24**) ^52^. Greater SV numbers were also observed in later-stage primary MSS cancers (P=9×10^-5^; **Supplementary Table 22**), supporting a relationship between tumour stage and chromosome instability.

Under a Cox proportional hazards model, total SV number was associated with shorter patient survival in the primary MSS cohort (P=2×10^-4^). However, mutation of major driver genes, total mutational burden, mutational signature activity, and immune escape status were not associated with patient survival after correction for multiple testing **(Supplementary Table 25).** No molecular associations with survival were found in the MSI cohort.

### Metastases

We compared genomic differences between the 1,354 MSS primaries and 105 unpaired MSS CRC metastases. Although there was no significant difference in mutation burden (Wilcoxon signed rank test; P=0.0731, **Supplementary Fig. 27a),** metastatic cancers tended to have higher ploidy (Wilcoxon signed rank test; P=3.67×10^-5^; **Supplementary Fig. 27b).** This is, consistent with a previous study, which found frequent whole genome doubling in metastatic CRC ^48^). When comparing copy number aberrations between metastases and primaries using 1Mb bins, no significant differences were found **(Supplementary Fig. 27c).** However, when copy number was considered relative to mean ploidy, several regions differed significantly between primaries and metastases (FDR<0.05; **Supplementary Fig. 27d).** Of note, 8p23-8p12 was a region where metastases had significantly lower copy number, recapitulating previous research which posited 8p as a ‘metastatic susceptibility locus’^53^. Whilst differences in immune pressure between primary and metastatic CRCs might be expected due to changes in tumour microenvironment, we detected no significant difference in immune escape frequency (P=0.142; **Supplementary Fig. 28a)** or neoantigen burden (P=0.44; **Supplementary Fig. 28b).**

### The CRC microbiome

We levered cancer sequence reads mapping to microbial genomes to identify microbial taxa. After removal of a high and variable level of likely hospital laboratory contaminants **(Supplementary Fig. 29),** we identified multiple bacterial species, predominantly from the genera *Bacteroides, Fusobacterium, Shigella, Streptococcus* and *Prevotella* **(Supplementary Fig. 30).** We also detected a small number of human viruses (mostly Herpes species) and multiple species of phage. We searched for associations between microbial content and HLA alleles, but none was found (P>0.05). Comparing bacterial load and diversity **(Supplementary Fig. 31),** 98.5% of primary tumours had over 0.001 bacterial cells per human cell, but in contrast to a previous report^54^, most metastases had almost no microbial content. The few metastases with a bacterial content comparable to primary tumours displayed similar taxa. Proximal MSS cancers had greater bacterial load than distal MSS cancers (P=8×10^-6^) but similar diversity. In the proximal colon, MSS cancers had fewer bacteria (P=9×10^-12^), but greater diversity (P=4×10^-9^), than MSI cancers. Anatomic site of the tumour and MSI status had the largest associations with bacterial diversity, but the associations of most other clinicopathological factors were also significant **(Supplementary Fig. 32).** In multivariate analyses of common taxa, several associations with CRC type and location were found. For example, *Fusobacterium* was more abundant in MSI than MSS cancers (Q=5×10^-5^), and *Akkermansia, Roseburia* and *Prevotella* were more abundant in proximal than distal colon (Q=5×10^-5^, Q=2×10^-6^ and Q=6×10^-4^ respectively; **Supplementary Tables 26-28, Supplementary Fig. 33).** *E.coli* abundance; P=2.9×10^-7^ and ID18 activity (PKS-pathogenic *E. coli* exposure; P=4×10^-10^) were both greater in the distal colon and rectum than the proximal colon. There was, however, no association between *E.coli* abundance and SBS88/ID18 activity once anatomical site was accounted for (P>0.05, **Supplementary Figure 34).**

### Clinical actionability and precision medicine

Using OncoKB and COSMIC Mutation Actionability in Precision Oncology databases, genetic alterations for which an approved therapy is indicated were identified in 25% (497/2,023) samples **(Supplementary Table 29).** The most frequent actionable genetic features were high tumour mutational burden (21%), microsatellite instability (18%), and *BRAF* V600E (14%). Genetic features with compelling evidence for predicting response to a therapy, as annotated by OncoKB, were present in 64% (1,304/2,023) of cancers **(Supplementary Table 30).** In our analysis, we identified 33 coding driver oncogenes that are not currently catalogued by COSMIC and OncoKB, raising the prospect that these genes may be viable targets for novel therapeutic intervention **(Supplementary Table 31).** Based on DepMap data, 22 of the 33 genes are predicted to be essential with positive selectivity (the degree to which essentiality varies across cell lines)^55^, 9 have a bioactive compound available, and 17 possess a druggable structure or are druggable by ligand-based assessment.

## DISCUSSION

Our study represents not only the largest WGS analysis of CRC to date, but the largest WGS analysis of any single cancer. This has enabled us to confirm previously reported CRC coding driver genes, to show for the first time that 51 driver genes in other cancers are also CRC drivers, and to identify 66 CRC driver genes not previously implicated as a driver in any cancer. Our WGS analysis has also allowed the identification of CNA and SV mutations that target these driver genes. Furthermore, the data set provided sufficient power to perform separate analyses of the MSI and POL subgroups, with clear benefits in identifying driver genes specific to these types, either resulting from their inherent hypermutational bias (*e.g. FAM83B*) or their distinct biology (e.g. immune escape mutations). We acknowledge that the set of putative drivers in MSI and POL tumours may contain some passengers that contain especially mutable sequences, although many of these drivers are also present in MSS cancers, even if they were not originally detected in that group; additional functional analyses will help to sift out these errors. We are able to conclude that all CRCs are likely to require constitutive Wnt activation through mutation, with inactivation of BMP/TGFβ signalling and/or Ras pathway activation in about half of cases. We found no additional, recurrent defects in base level DNA repair to sit alongside the MSI and POL types of CRC, but we did find a small number of MSS CRCs with specific forms of chromosomal instability, including HRD cancers with *BRCA1/2* mutations that might benefit from the personalized use of drugs such as PARP inhibitors^56^. The use of WGS has enabled the identification of recurrently mutated non-coding elements, including *APC* and *SMAD4* splice regions, and *ST6GALNAC1* distal promoter. However, for mutations outside gene bodies in particular, additional functional analyses will be required to established driver status.

A major advantage of the 100,000 Genomes Project is its integration of NHS clinical data^57^. This facilitates ongoing relation of genomic features to histopathological, treatment and outcome information, something not possible in large-scale cancer sequencing projects such as PCAWG ^58^. For example, we could link mutational signatures with anti-cancer therapies and observe radiation-associated signatures in CRCs from patients who had previously received radiotherapy for prostate cancer^45^. Furthermore, correlation of patient age with mutational profiles revealed both clock-like signatures (SBS1, SBS5, SBS18, ID1) and signatures enriched in younger patients’ cancers (SBS89, SBS93, ID14), supporting aetiopathological differences between early- and late-onset CRCs^59^. A clear relationship between CRC anatomical site and mutational profile was apparent, reflected in driver mutation frequencies and underlying mutational processes. For example, even in MSS cancers, there is greater SBS8 and ID18, and lower SBS1, SBS5, SBS18, ID1 and ID2 activity, the further the cancer from the start of the large bowel (caecum). Similar gradients in the CRC microbiome hint that some of these differences might be caused by mutagenic bacterial toxins, akin to those from PKS-pathogenic *E. coli* which are most prevalent in CRCs from the distal colon and rectum. Overall, our study provides new insights into CRC biology. It demonstrates the benefits of WGS over exome or panel sequencing for cancer clinical diagnostics. The large resource of quality-controlled and annotated CRC genomes should provide the basis for multiple research studies spanning from CRC predisposition through to multi-omic studies of pathogenic mutations outside the coding genome as well as drug discovery projects.

## AUTHOR CONTRIBUTIONS

D. Chubb., A.J.C., N.M. processed clinical data. A.J.C., A.J.G., G.C., W.C. performed sequencing data quality control. D.Chubb., B.N., A.Sosinsky, J.M., G.C., W.C. performed quality control of simple somatic mutations. A.F., G.C., J.H., W.C., A.J.C. called copy number alterations. D.Chubb evaluated microsatellite instability. A.J.C. called structural variants and identified recurrent events. A.F., B.K., A.J.C., and M.N.L. identified recurrent copy number alterations. B.K., C.A.P., D.Church identified driver mutations. A.J.G., D.Chubb., A.J.C., B.K., and A.H. extracted mutational signatures with help from L.B.A. A.J.C. identified biallelic events. R.C., A.J.C., S.T. analysed disrupted pathways. E.L. profiled the immune landscape. R.C. characterised the mitochondrial genome. J.H. analysed metastases. A.J.C. evaluated the genomic impact of prior treatments. H.M.W., B.N. analysed the microbiome. B.N., R.C. performed survival analysis. A.J.C. correlated genomic and clinical features. A. Sud, and B.K performed clinical actionability analyses. B.K. estimated telomere content. B.K. identified extrachromosomal DNA. B.K. inferred driver mutation clonality. A.S., D.Church, D.C.W., I.T., P.Q., R.S.H., T.G. supervised the study. A.J.C. A.J.G., A.F., B.K., C.A.P., D.Chubb, D.Church, E. L., H.M.W., J.H., R.C., R.S.H., I.T. wrote the manuscript. All authors read and approved the final version of this manuscript.

## ACKNOWLEDGEMENTS

R.S.H., I.T and N.L.B. are supported by the Wellcome Trust (214388), T.G. and A.S are supported by the Wellcome Trust (202778/B/16/Z). I.T. is supported by Cancer Research UK (C6199/A27327). R.S.H. is supported by Cancer Research UK (C1298/A8362). A.S. is supported by Cancer Research UK (A22909). T.G. is supported by Cancer Research UK (A19771 and DRCNPG-May21\100001). P.Q. and H.M.W. are supported by Cancer Research UK Grand Challenge Initiative (OPTIMISTICC C10674/A27140). P.Q. is also supported by Yorkshire Cancer Research L386 and is a National Institute of Health Senior Investigator. B.N was funded through the Cancer Research UK Birmingham Centre award (C17422/A25154). C.A.-P. was supported by “la Caixa” Foundation (ID 100010434) fellowship (LCF/BQ/ES18/11670011). I.T. is funded by the Wellcome Trust (214388), Cancer Research UK (C6199/A27327) and the Cancer Research UK Scotland Cancer Centre (CTRQQR-2021\100006). L.B.A was supported by grants from the US National Institutes of Health, including: NIEHS R01ES032547 and NCI R01CA269919. G. Caravagna is supported by the Italian Association for Cancer Research (AIRC) under MFAG 2020-ID 24913. We acknowledge funding from the National Institute of Health (NCI U54 CA217376) to A.S and T.G.. This work was also supported a Wellcome Trust award to the Centre for Evolution and Cancer at the ICR (105104/Z/14/Z). D.N.C. is supported a by Cancer Research UK Advanced Clinician Scientist Fellowship award (C26642/A27963). D.N.C, D.C.W. and A.F. acknowledge support from the Oxford NIHR Comprehensive Biomedical Research Centre. (The views expressed are those of the authors and not necessarily those of the NHS, the NIHR, or the Department of Health.) N.L.-B. acknowledges funding from the European Research Council (consolidator grant 682398) and ERDF/Spanish Ministry of Science, Innovation and Universities – Spanish State Research Agency/DamReMap Project (RTI2018-094095-B-I00) and Asociación Española Contra el Cáncer (AECC) (GC16173697BIGA). IRB Barcelona is a recipient of a Severo Ochoa Centre of Excellence Award from the Spanish Ministry of Economy and Competitiveness (MINECO; Government of Spain) and is supported by CERCA (Generalitat de Catalunya).

This research was made possible through access to the data and findings generated by The 100,000 Genomes Project. The 100,000 Genomes Project is managed by Genomics England Limited (a wholly owned company of the Department of Health) funded by the National Institute for Health Research and NHS England. The Wellcome Trust, Cancer Research UK and the Medical Research Council have also funded research infrastructure. The 100,000 Genomes Project uses data provided by patients and collected by the National Health Service as part of their care and support. We are grateful to all individuals involved.

## GENOMICS ENGLAND RESEARCH CONSORTIUM

J. C. Ambrose^1^, P. Arumugam^1^, E. L. Baple^1^, M. Bleda^1^, F. Boardman-Pretty^1,2^, J. M. Boissiere^1^, C. R. Boustred^1^, H. Brittain^1^, M. J. Caulfield^1,2^, G. C. Chan^1^, C. E. H. Craig^1^, L. C. Daugherty^1^, A. de Burca^1^, A. Devereau^1^, G. Elgar^1.2^, R. E. Foulger^1^, T. Fowler^1^, P. Furió-Tarí^1^, J. M. Hackett^1^, D. Halai^1^, A. Hamblin^1^, S. Henderson^1,2^, J. E. Holman^1^, T. J. P. Hubbard^1^, K. Ibáñez^1,2^, R. Jackson^1^, L. J. Jones^1,2^, D. Kasperaviciute^1,2^, M. Kayikci^1^, L. Lahnstein^1^, L. Lawson^1^, S. E. A. Leigh^1^, I. U. S. Leong^1^, F. J. Lopez^1^, F. Maleady-Crowe^1^, J. Mason^1^, E. M. McDonagh^1,2^, L. Moutsianas^1,2^, M. Mueller^1,2^, N. Murugaesu^1^, A. C. Need^1,2^, C. A. Odhams^1^, C. Patch^1,2^, D. Perez-Gil^1^, D. Polychronopoulos^1^, J. Pullinger^1^, T. Rahim^1^, A. Rendon^1^, P. Riesgo-Ferreiro^1^, T. Rogers^1^, M. Ryten^1^, K. Savage^1^, K. Sawant^1^, R. H. Scott^1^, A. Siddiq^1^, A. Sieghart^1^, D. Smedley^1,2^, K. R. Smith^1,2^, A. Sosinsky^1,2^, W. Spooner^1^, H. E. Stevens^1^, A. Stuckey^1^, R. Sultana^1^, E. R. A. Thomas^1,2^, S. R. Thompson^1^, C. Tregidgo^1^, A. Tucci^1,2^, E. Walsh^1^, S. A. Watters^1^, M. J. Welland^1^, E. Williams^1^, K. Witkowska^1,2^, S. M. Wood^1,2^ & M. Zarowiecki^1^

1. Genomics England, London, UK.

2. William Harvey Research Institute, Queen Mary University of London, London, UK.

