## Supplementary Figures for "Whole genome sequencing of 2,023 colorectal cancers reveals mutational landscapes, new driver genes and immune interactions"

**Supplementary Fig. 1: Overview of structural-variant-calling pipeline.** BAM: binary sequence alignment map, PCAWG: The Pan-Cancer Analysis of Whole Genomes, SV: structural variant.

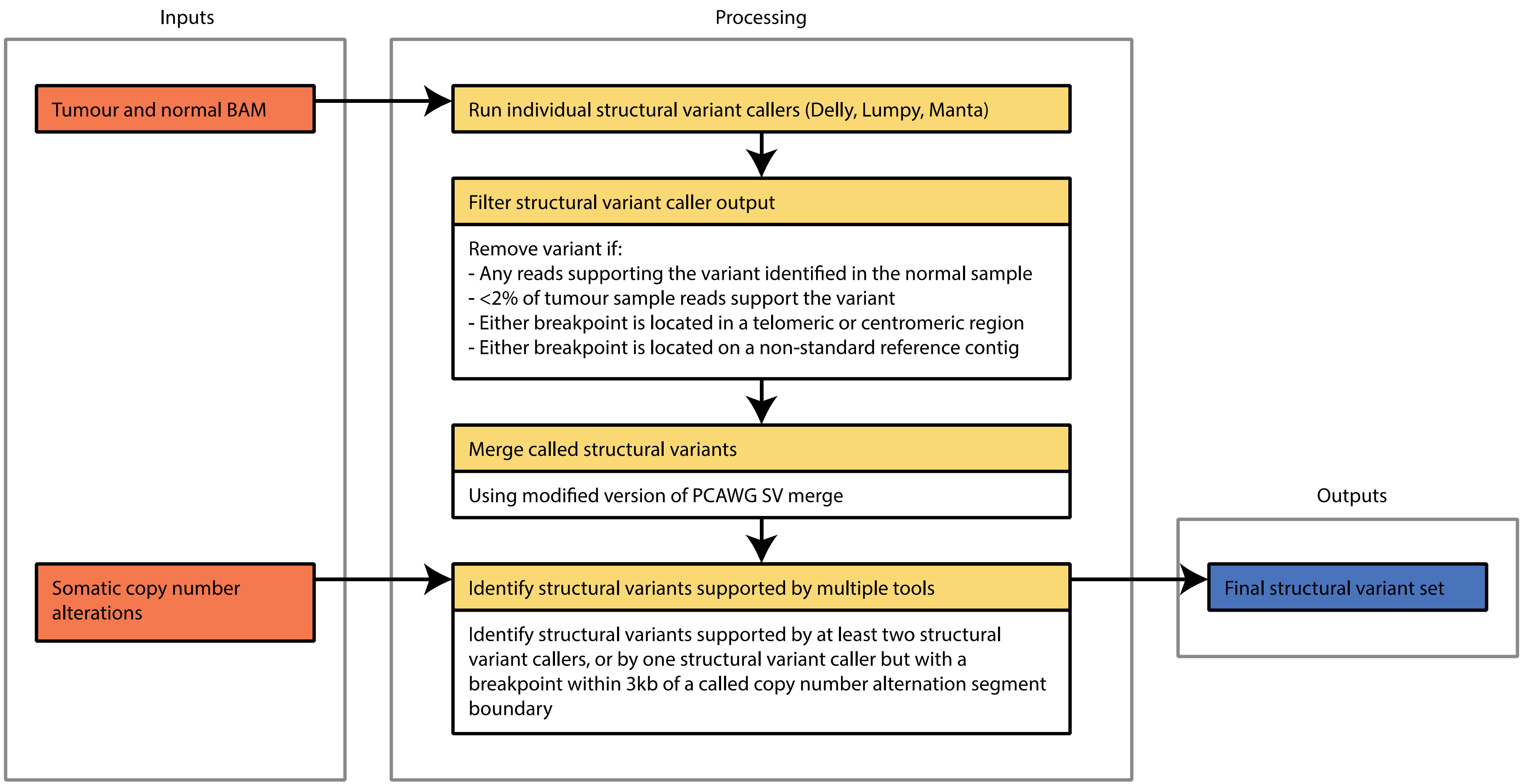

**Supplementary Fig. 2: Association between SNV VAF and numbers of driver mutations called in primary microsatellite stable (MSS) colorectal cancer (CRC) samples (n=1956), stratified by median single nucleotide variant (SNV) variant allele frequency (VAF).** Potential driver mutations were defined as any coding variant annotated as probably pathogenic from 63 driver genes previously identified in MSS CRC [1-4]. Below the dashed line, it is likely that low median VAF results from low cancer cell purity and this causes a failure to identify some driver mutations. Tumour samples below the threshold were therefore excluded from the analysis.

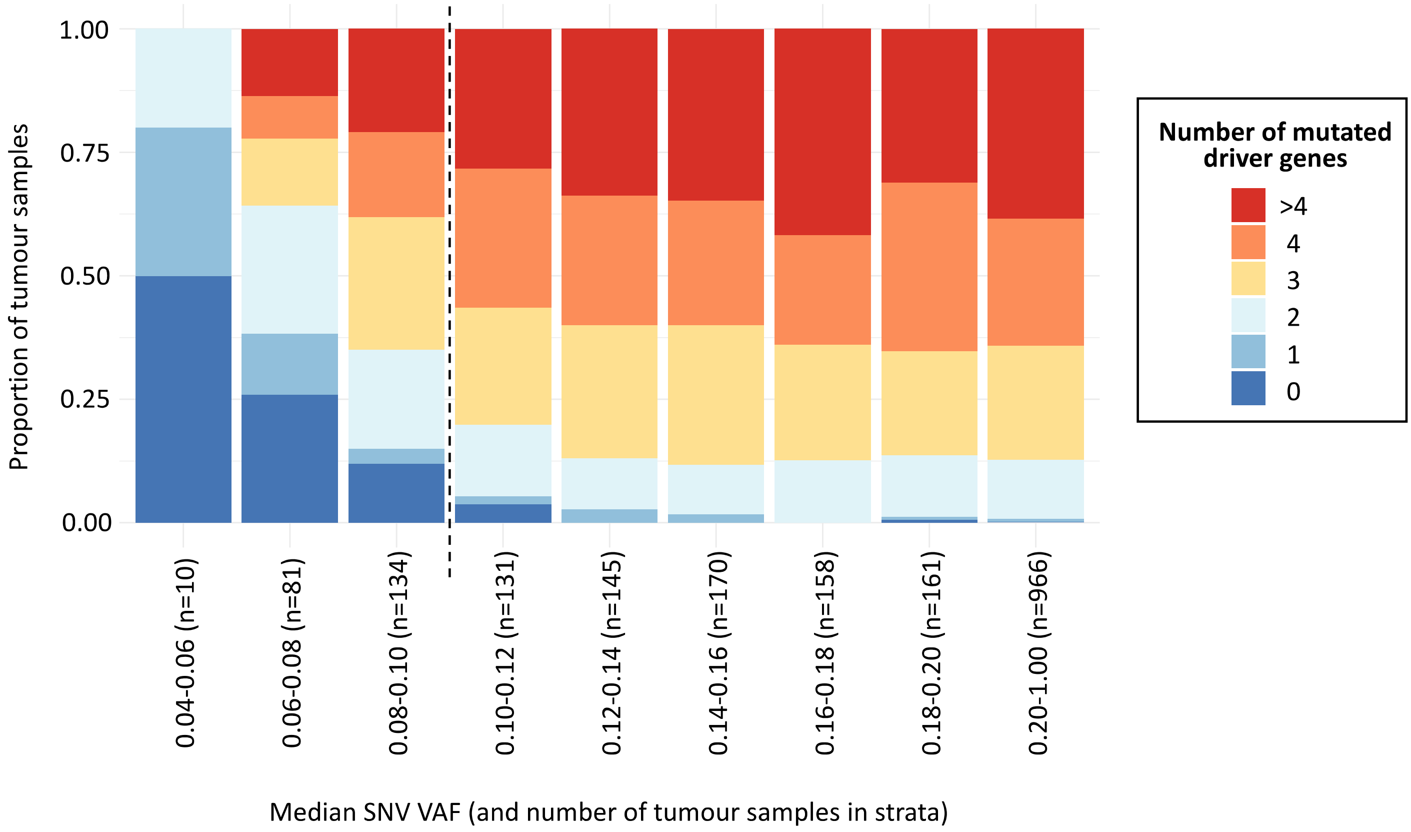

**Supplementary Fig. 3: Removing bias introduced by soft clipping of semi-aligned reads.** An alignment of six reads to a reference sequence containing an A/T variant is shown. Bold black T and red A represent reference and alternate alleles respectively. Soft clipping is represented by strikethrough. Without soft clipping, three reads would support both the reference (T) and alternate (A) alleles, resulting in an unbiased variant allele frequency (VAF) of 3/6=0.5. (**a**) Read R3 is soft clipped until five consecutive matches with the reference are obtained. After clipping, only two reads support the alternate allele (A), whilst three reads support the reference allele (T), resulting in a biased VAF of 2/5=0.4. (**b**) FixVAF clips all reads by five bases, regardless of whether they contain a variant site or support a reference or alternate allele. Reads supporting both the reference and alternate alleles are now clipped by five bases. In this example, FixVAF would compute a VAF of 2/4=0.5, and therefore remove bias.

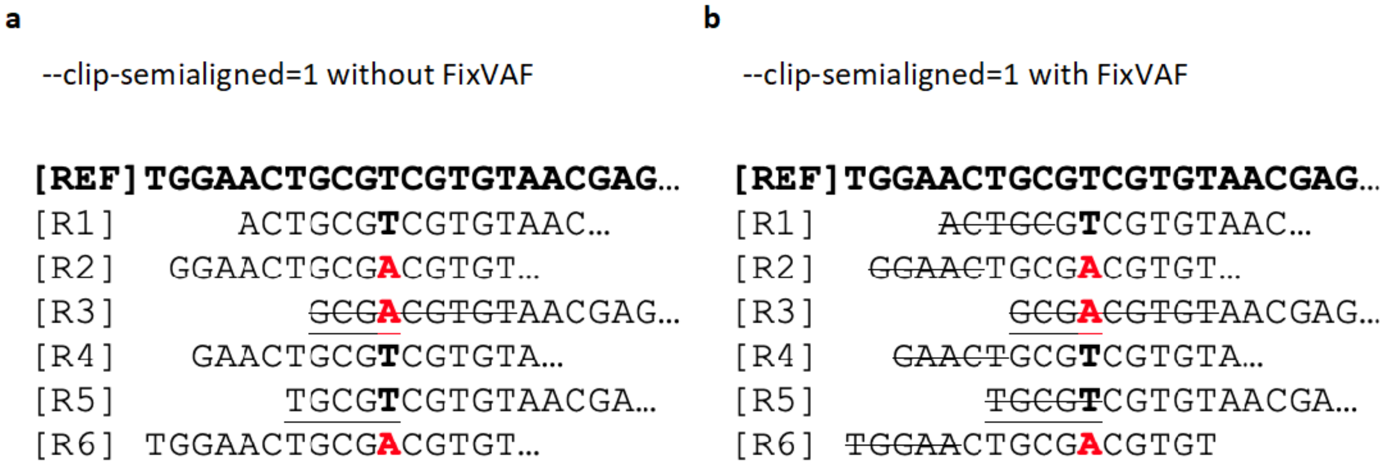

**Supplementary Fig. 4: Overview of copy number aberration calling pipeline.** BAM: binary sequence alignment map, CCF: cancer cell fraction, SNV: single nucleotide variant, vcf: variant call format.

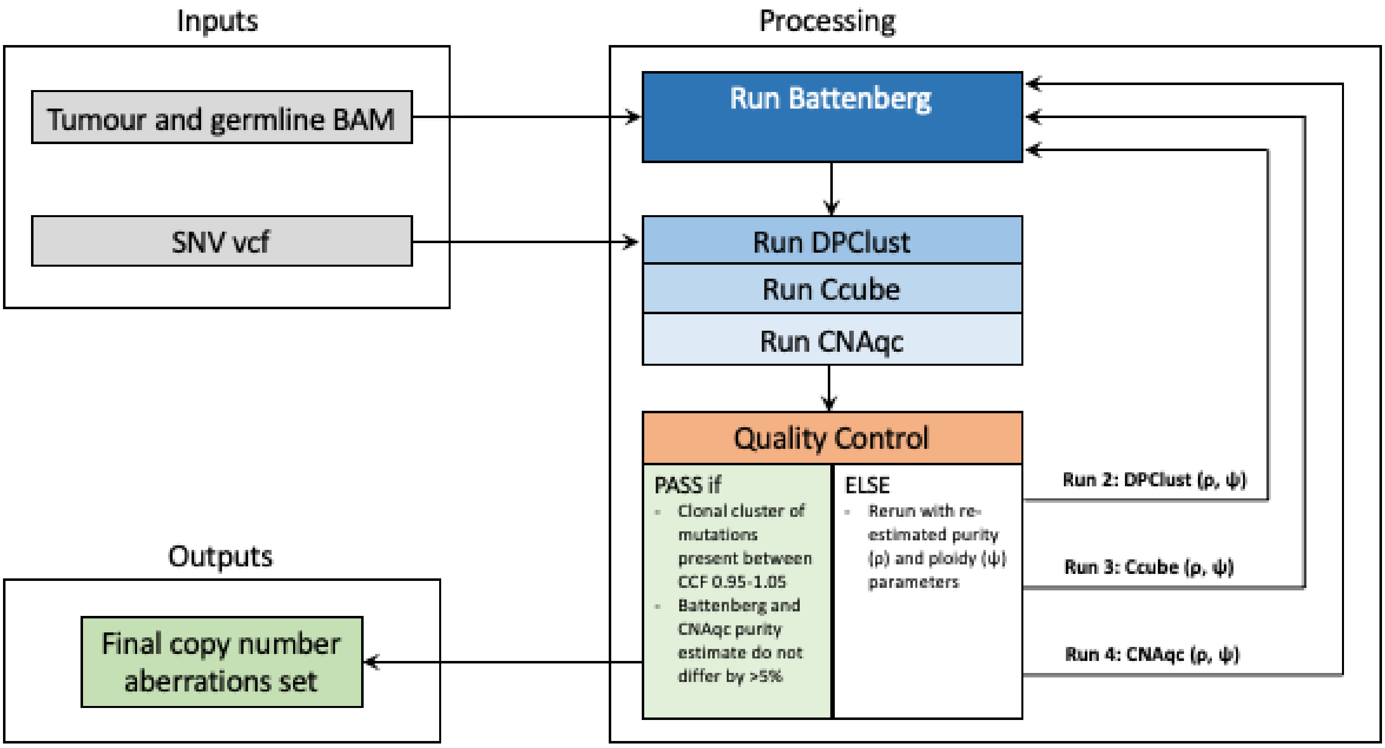

**Supplementary Fig. 5: Number of mutations attributed to SBS93 in tumours with and without *TP53* mutations.** MSS: microsatellite stable; MSI: microsatellite unstable. Significance tested using two-tailed Wilcoxon rank-sum test.

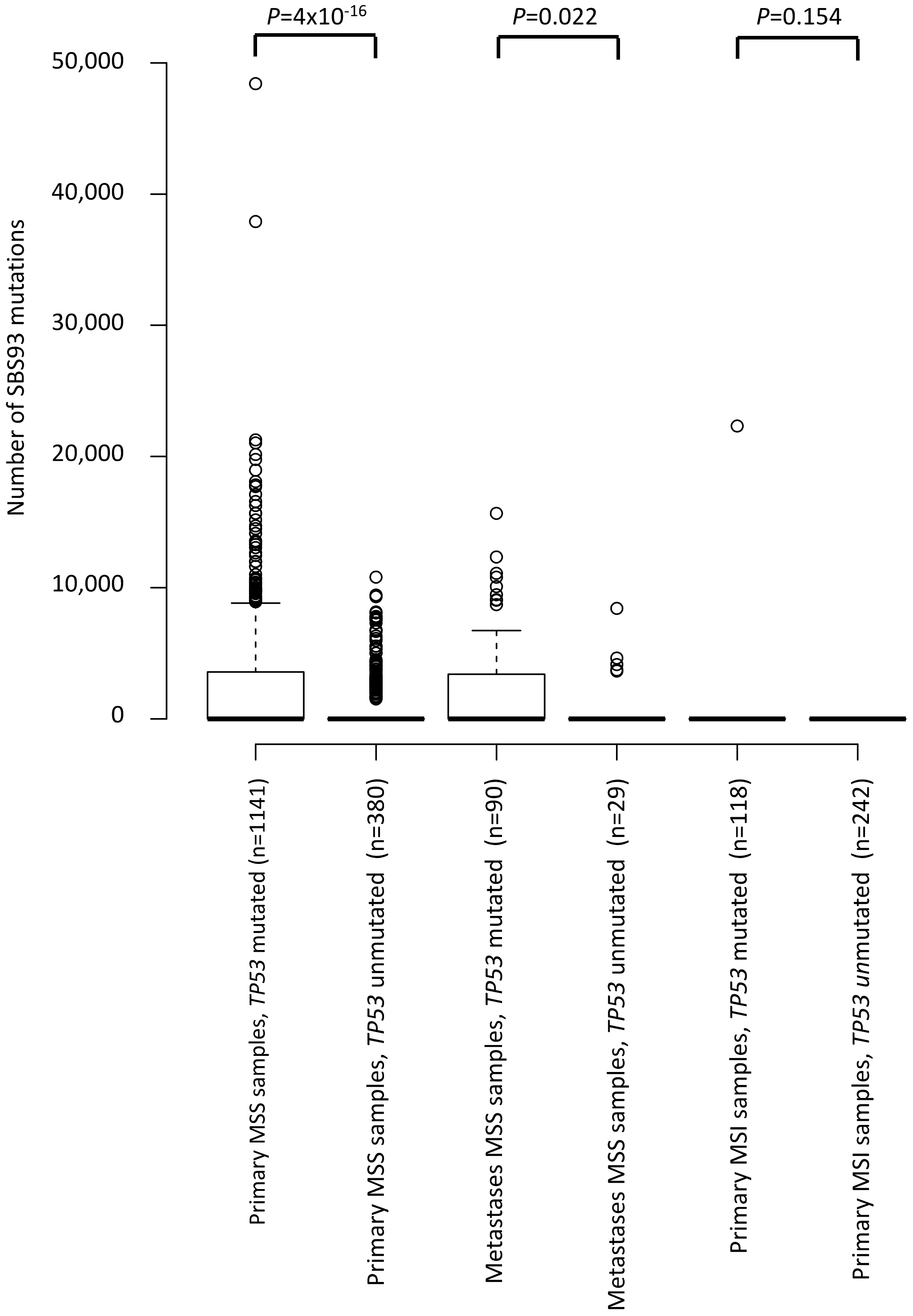

**Supplementary Fig. 6: Structural variant (SV) signature extraction.** (**a**) Multimodal SV size and replication timing distributions. Dashed lines represent thresholds used to categorize variants for signature extraction. (**b**) To assess stability of SV signature extraction using the hierarchical Dirichlet process (HDP), the cohort was randomly split into halves and signatures extracted independently from each. Nine signatures extracted from the cohort halves showed high similarity between halves (red and blue grey; cosine similarity >0.9) and high similarity with signatures extracted from the full cohort and were therefore included in subsequent analyses. MSS: microsatellite stab le; MSI:  microsatellite unstable.

**a**

**b**

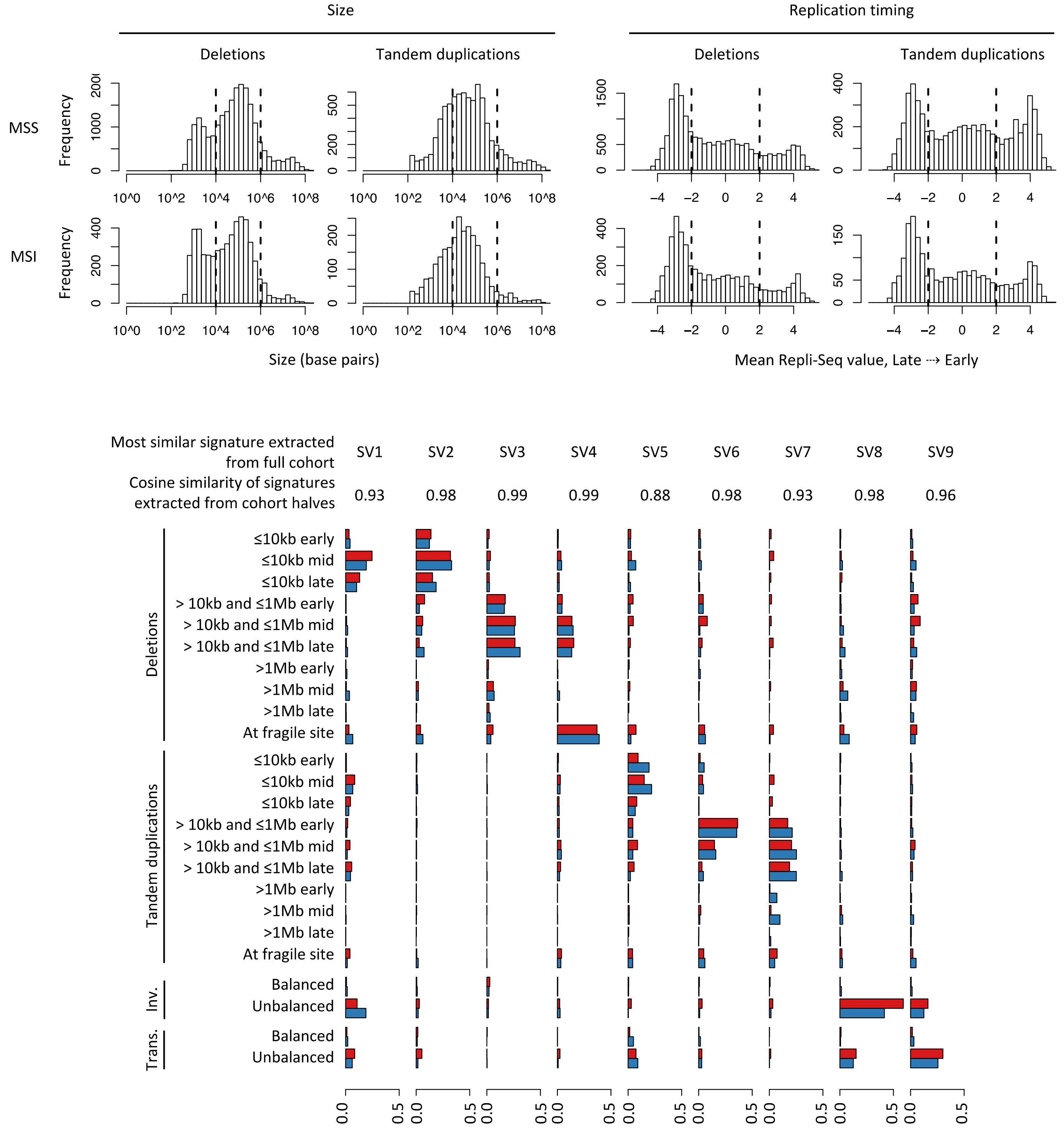

**Supplementary Figure 7: CN signatures. (i)** Deconvoluting the 4 de novo signatures using the set of 19 COSMIC CNV signatures, identified 6 contributing COSMIC signatures. CNV48A is a heterogeneous signature, dominated by heterozygous segments of 3-8 copies. It is decomposed in to three Cosmic signatures, CN15 – previously associated with HRD and TD, CN6 previously associated with chromothripsis and CN18 which has a currently unexplained aetiology. CNV48B is comprised primarily of heterozygous segments of 3-4 copies with a length of >40MB it is deconvoluted in to a single cosmic signature CN2, associated with tetraploidy. CNV48C is dominated by heterozygous segments with a copy number of 2 and is decomposed to CN1, indicative of a diploid state. CNV48D is dominated by LOH segments with a CN of 1 and heterozygous segments with a CN of 2 and to a lesser extent 3-4, it deconvoluted in to CN9 which has previously been associated with chromosomally instable diploid tumours. For each de-novo CN signature on the top left of each plot the contributing COSMIC signatures are provided on the right alongside the final refitted signature on the bottom left. A) CNV48A is comprised of 42.18% CN15, 29.72% CN6 and 28.1% CN18. B) CNV48B is comprised of CN2 C) CNV48C is comprised of CN1. D) CNV48D is comprised of CN9. **(ii)** Selection plot showing the Mean Sample Cosine difference and Average Stability for 1-30 copy number signature de-novo extractions. The accepted solution contained four de-novo signatures.

**(i)**

A) CNV48A

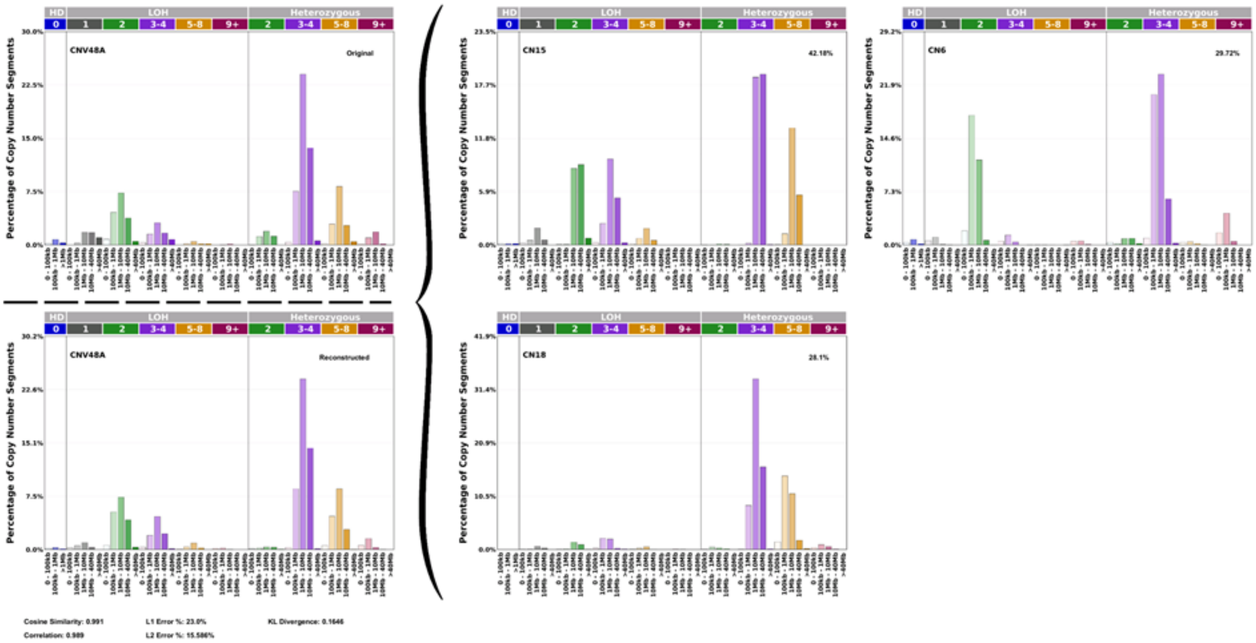

B) CNV48B

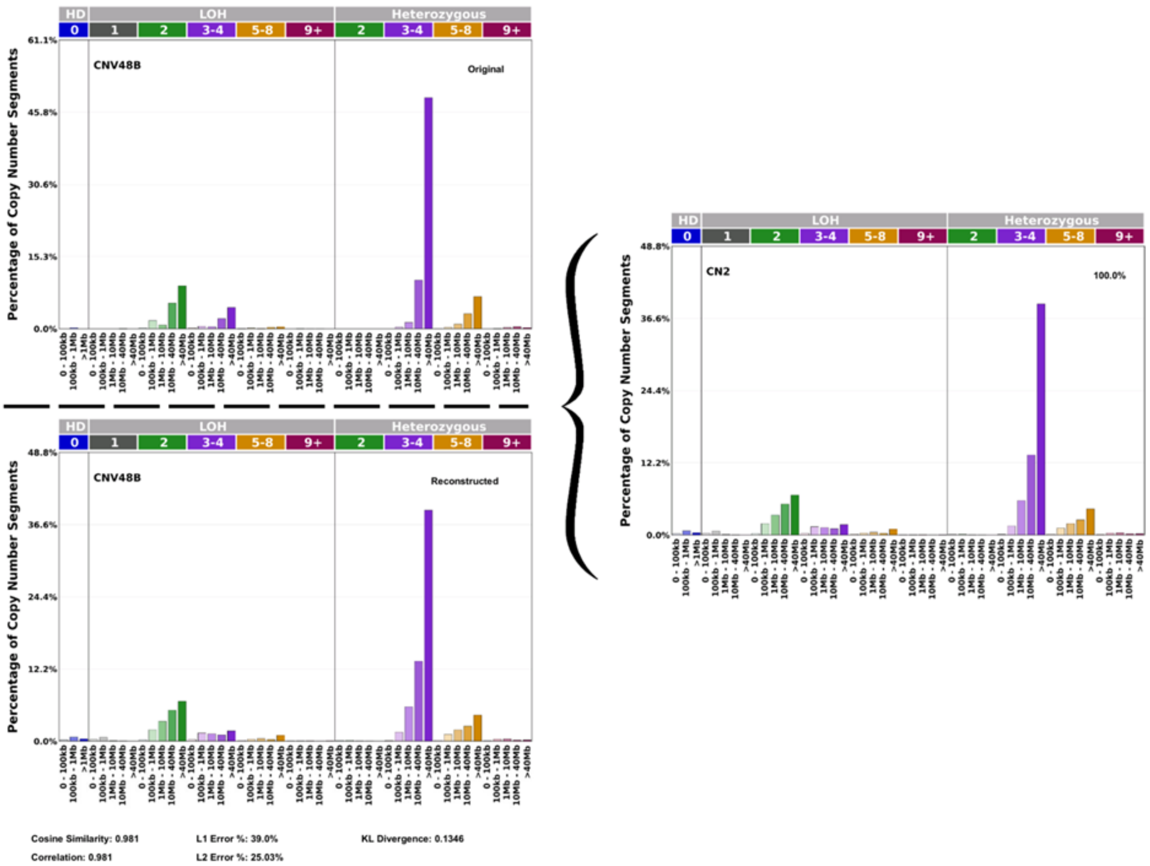


C) CNV48C

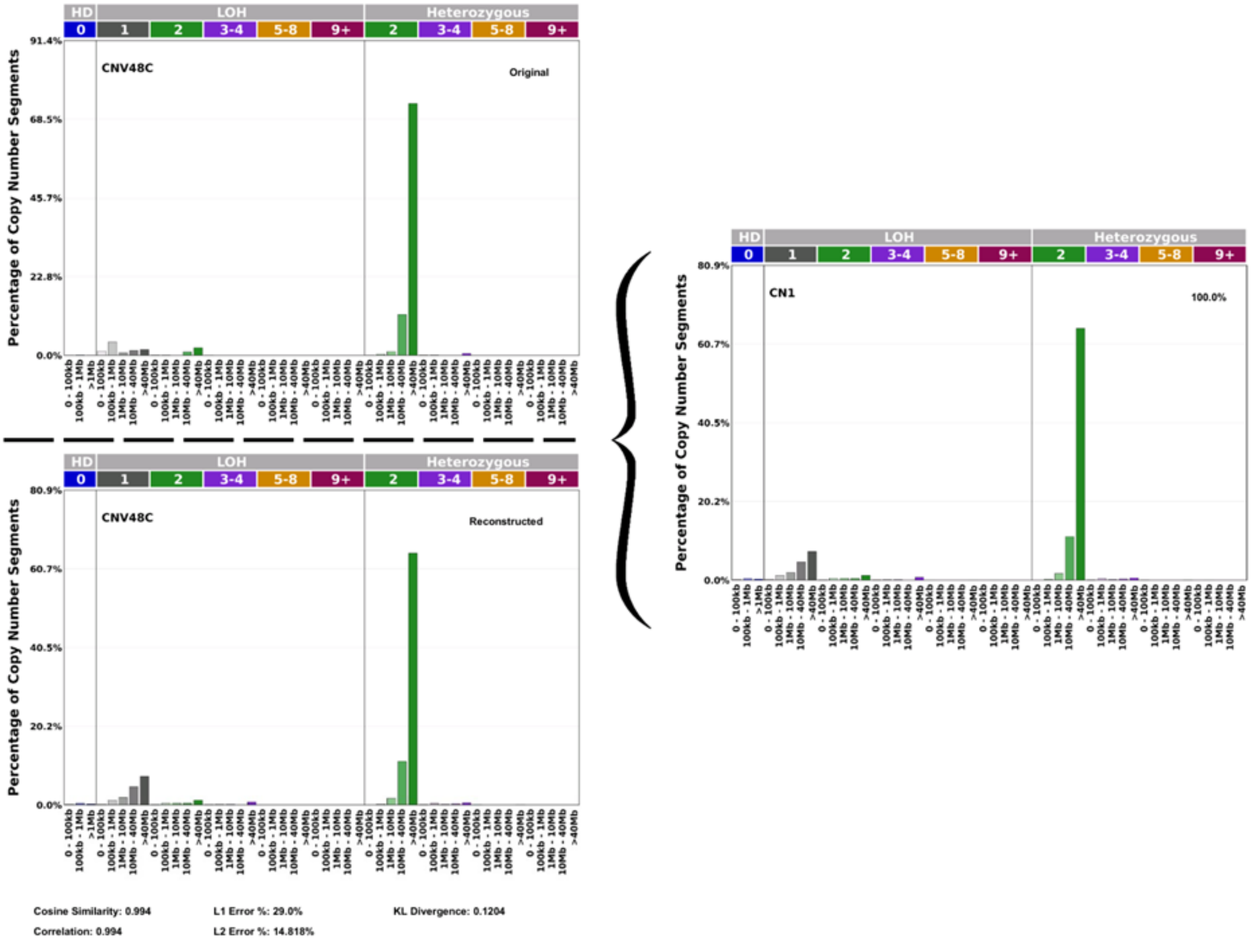

D) CNV48D

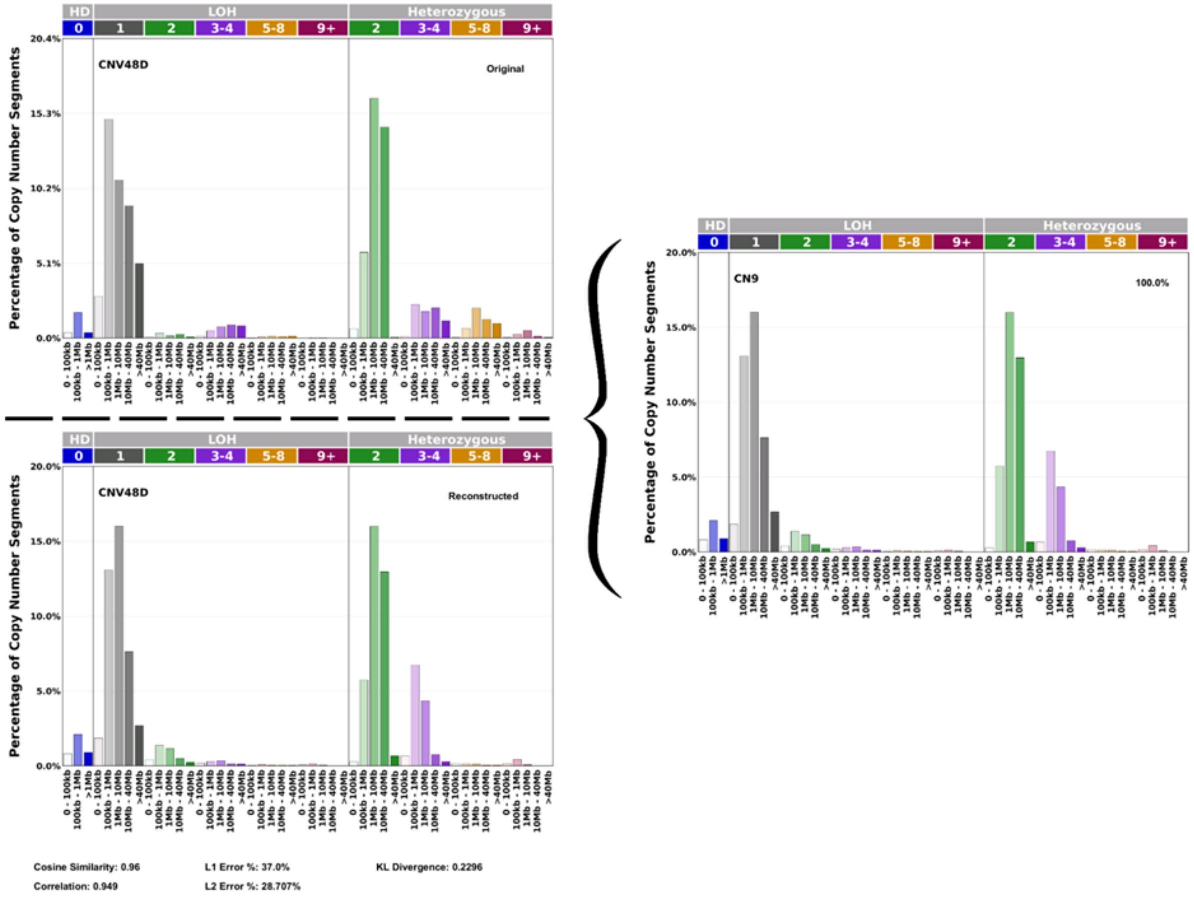

CNV48D
Original
Reconstructed
CN9
100.0%
HD
LOH
Heterozygous
Percentage of Copy Number Segments
Cosine Similarity: 0.96
Correlation: 0.949
L1 Error %: 37.0%
L2 Error %: 28.707%
KL Divergence: 0.2296


**(ii)**

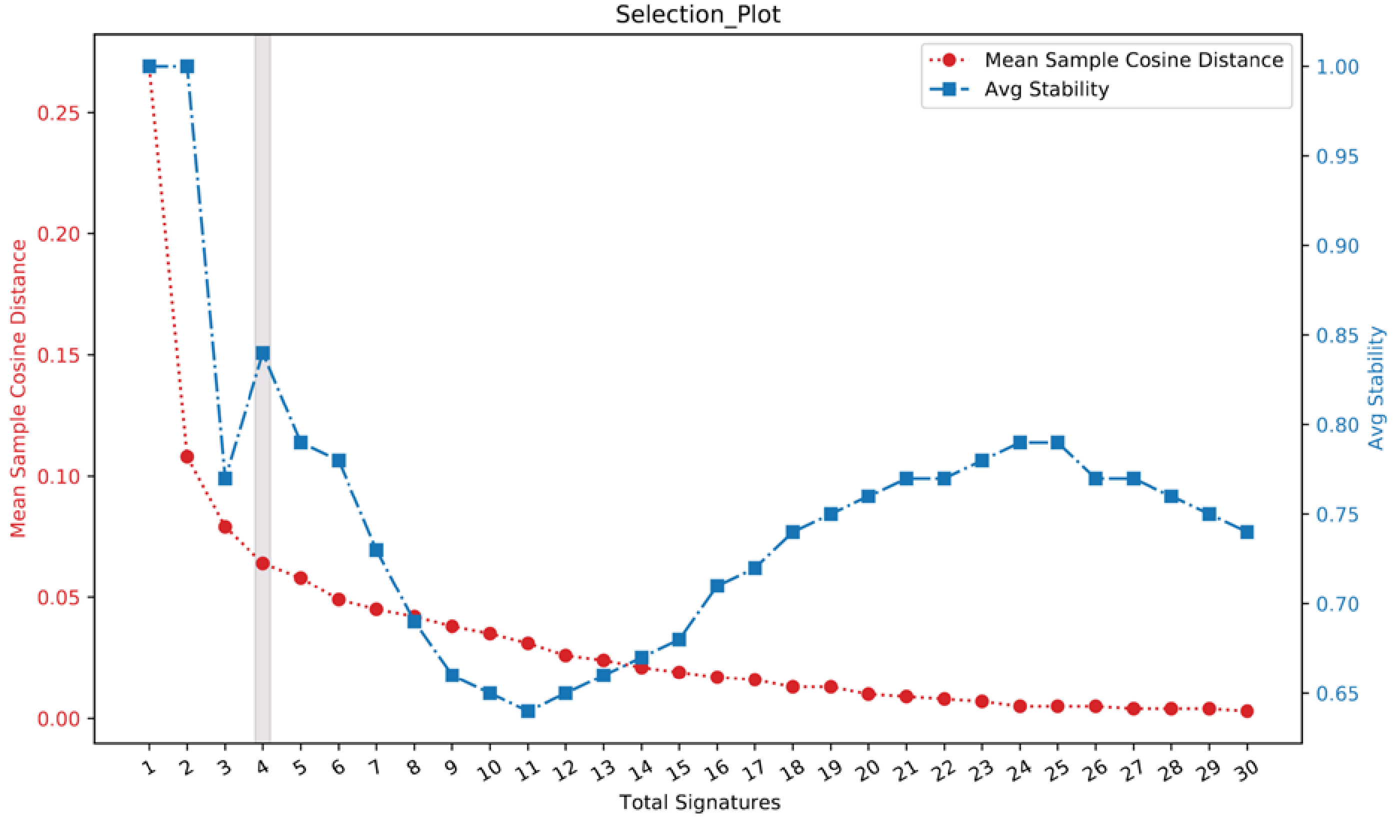

Selection_Plot
Mean Sample Cosine Distance
Avg Stability
Total Signatures

**Supplementary Fig. 8: Mutation status of *BRCA1/BRCA2* in tumours with and without predicted homologous recombination deficiency (HRD).** HRD was predicted in 1765 tumours with profiled copy number alterations using HRDetect with a probability threshold of 0.7. Germline single nucleotide polymorphisms and small insertion and deletions (indels) defined as moderate or high impact by Variant Effect Predictor (VEP) and reported as pathogenic or likely pathogenic by ClinVar (v1.20) were considered. Somatic single nucleotide variants and indels defined as moderate or high impact by VEP were considered. The mutation status of *BRCA1* and *BRCA2* was combined for each tumour. LOH: loss of heterozygosity.

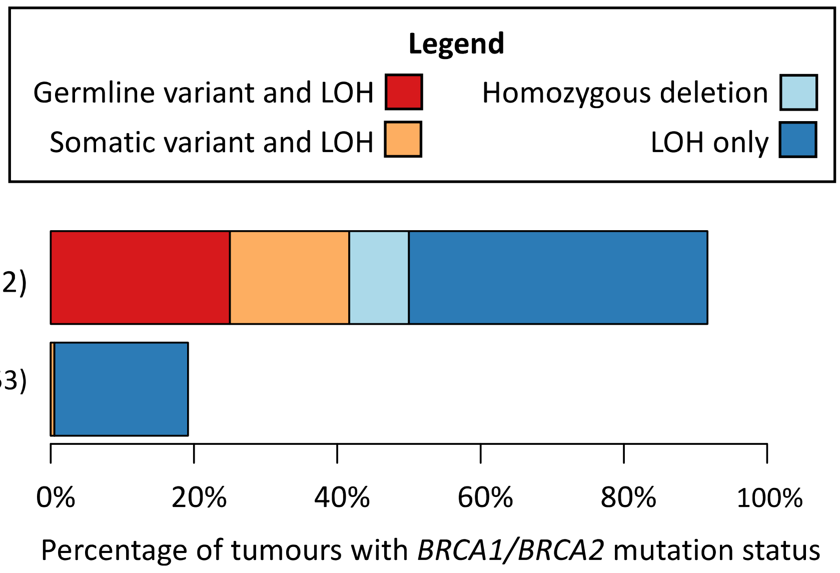

**Supplementary Figure 9: Distribution of per-tumour driver mutation counts by CRC subtype.** Oncogenic mutations from 185 candidate driver genes were included in the per-tumour counts. The Kruskal-Wallis test was used to estimate the significance of differences in driver mutation counts between subtypes.

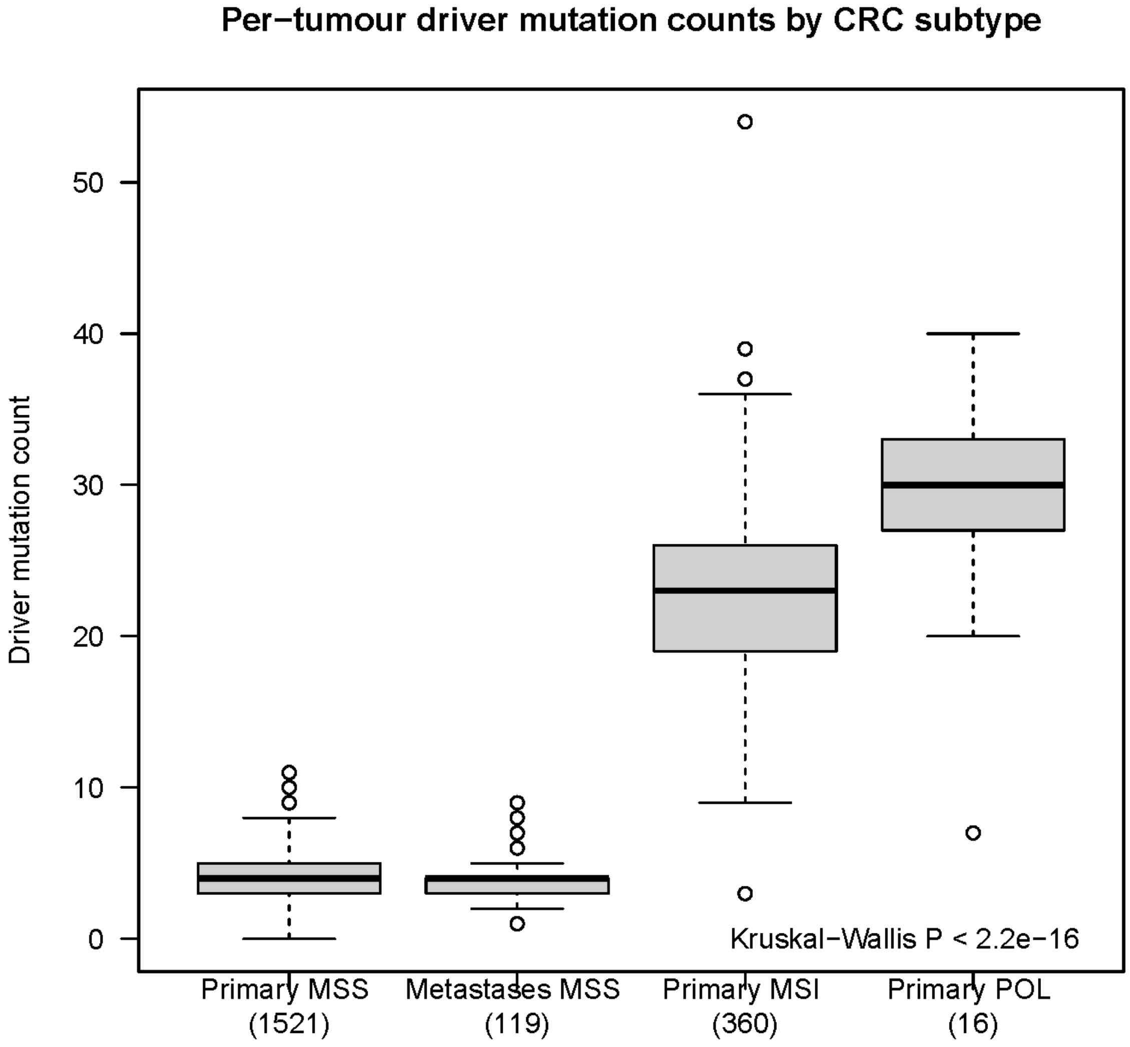

**Supplementary Fig. 10: Non-synonymous mutations in selected driver genes identified in primary microsatellite stable (MSS) tumours.** Coloured circles represent mutations predicted as oncogenic and are annotated with their predicted consequence. Faint circles represent variants of unknown significance.

**AKT1 (ENST00000554581)**

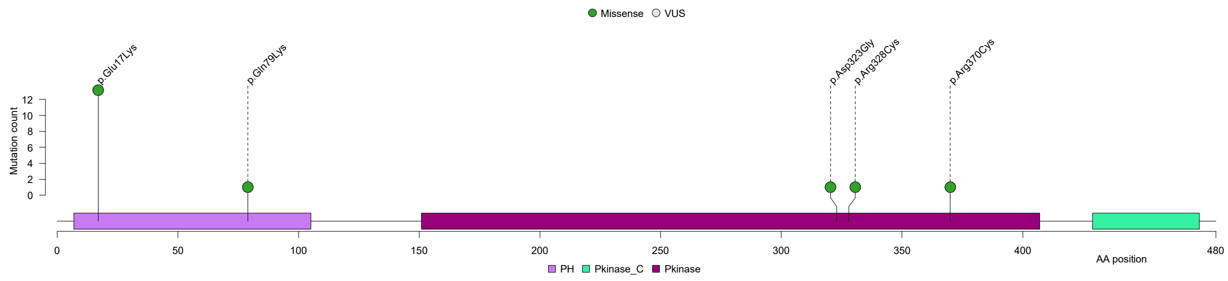


**ARID2 (ENST00000334344)**

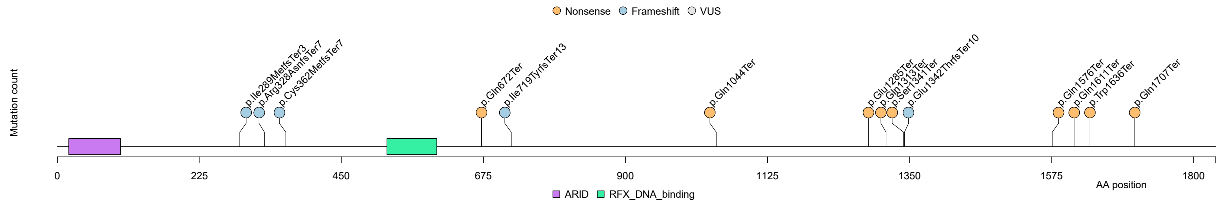


**CDKN2A (ENST00000498124)**

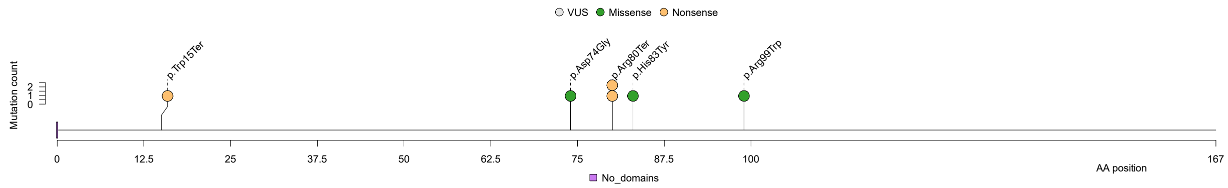


**CTNND1 (ENST00000399050)**

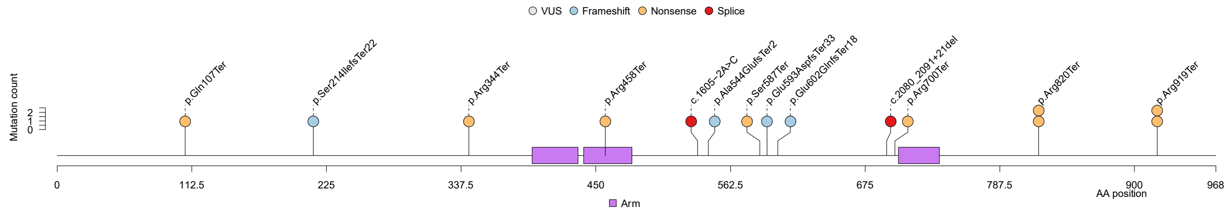

**IDH1 (ENST00000415913)**

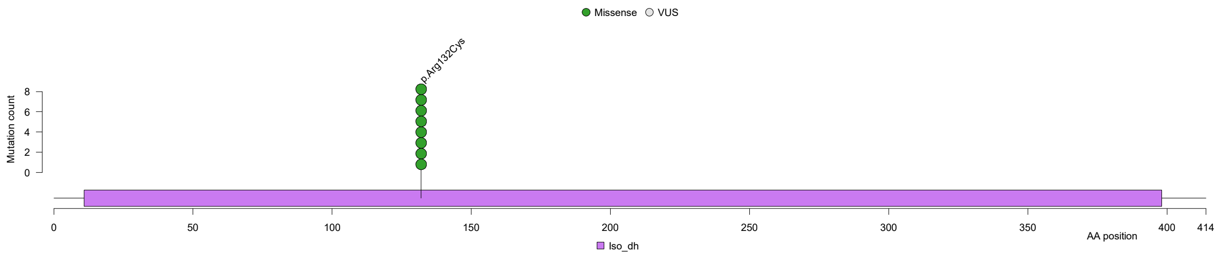


**RAF1 (ENST00000442415)**

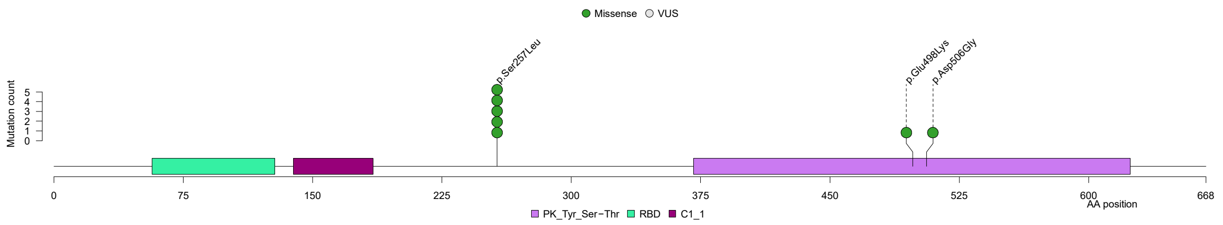


**RASGRF1 (ENST00000419573)**

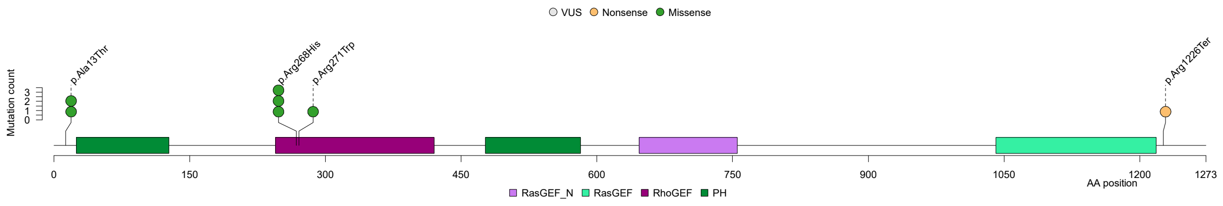


**PIK3R1 (ENST00000521381)**

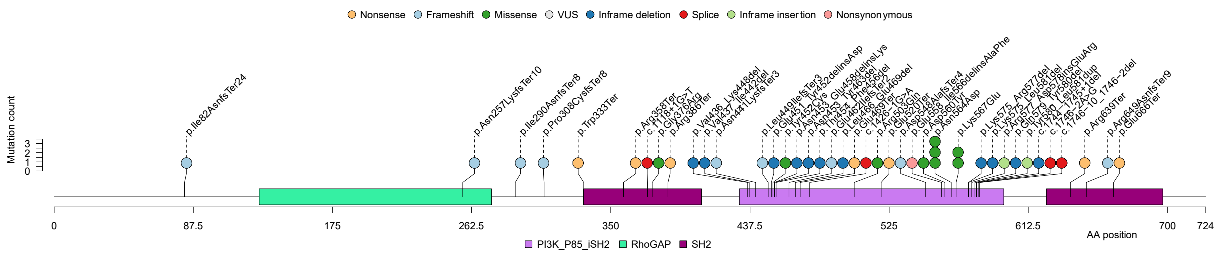

**Supplementary Fig. 11: Non-synonymous mutations in selected driver genes identified in primary microsatellite unstable (MSI) tumours.** Coloured circles represent mutations predicted as oncogenic and are annotated with their predicted consequence. Faint circles represent variants of unknown significance.

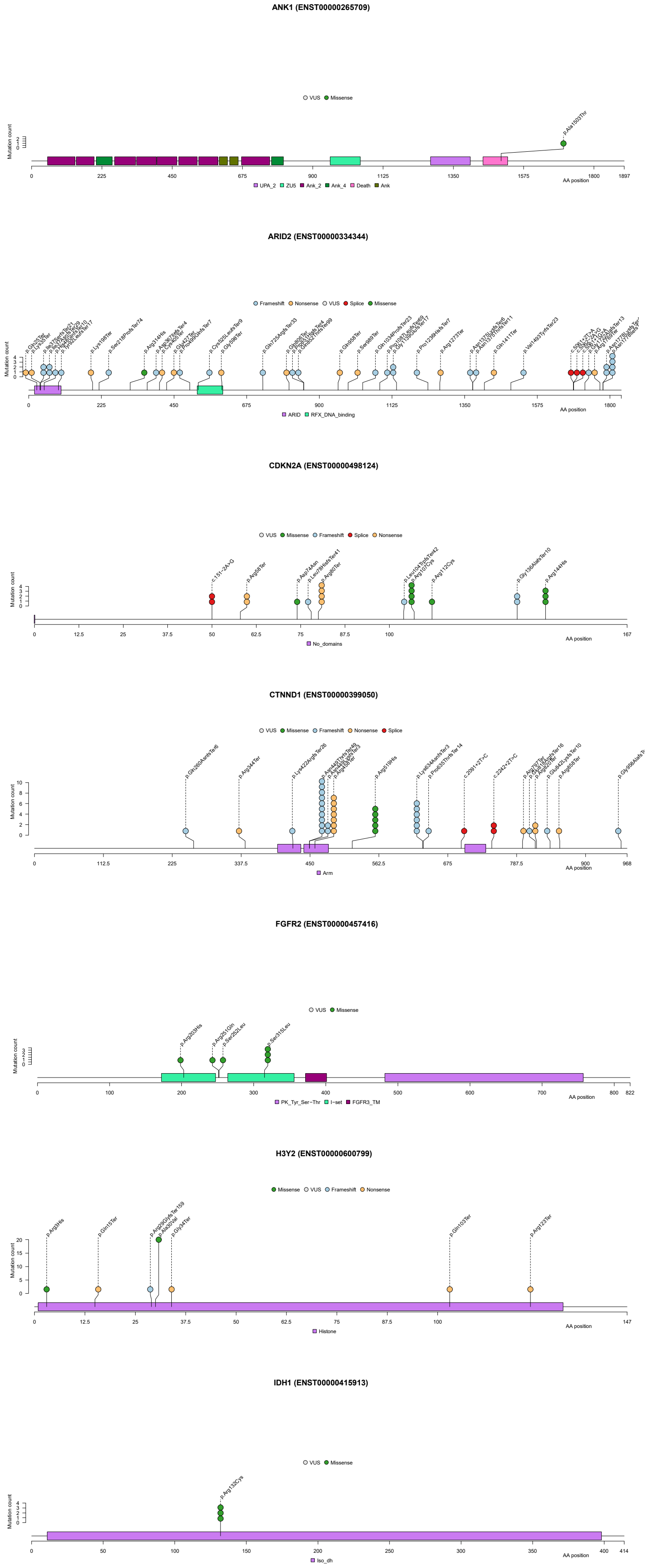

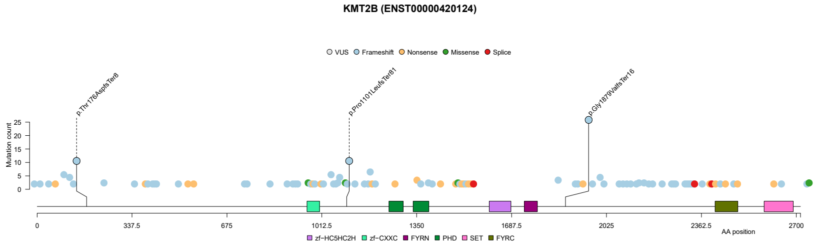
KMT2B (ENST00000420124)
VUS Frameshift Nonsense Missense Splice
AA position
zf−HC5HC2H zf−CXXC FYRN PHD SET FYRC

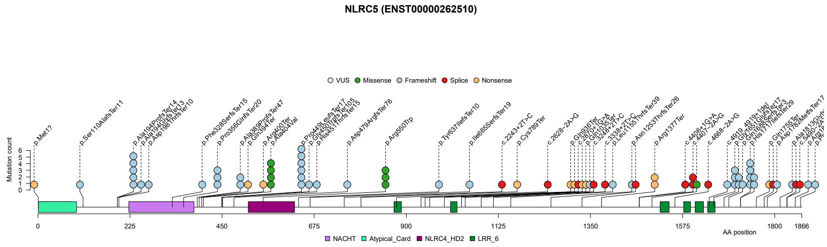
NLRC5 (ENST00000262510)
VUS Missense Frameshift Splice Nonsense
AA position
NACHT Atypical_Card NLRC4_HD2 LRR_6

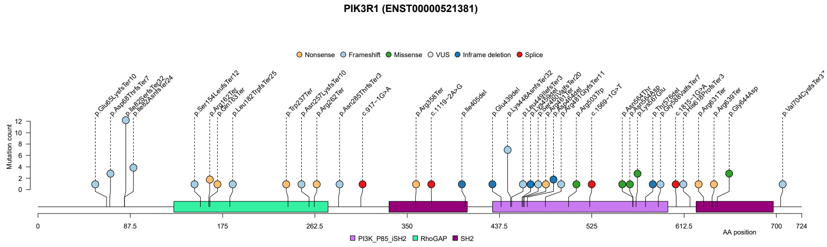
PIK3R1 (ENST00000521381)
Nonsense Frameshift Missense VUS Inframe deletion Splice
AA position
PI3K_P85_iSH2 RhoGAP SH2

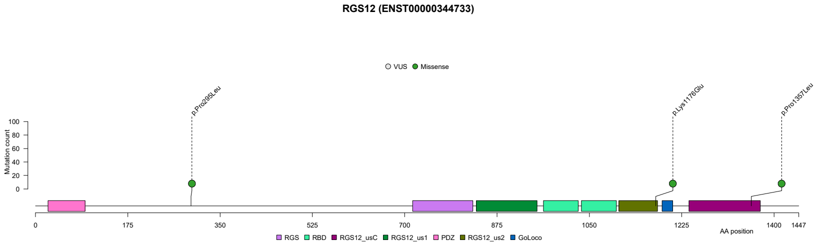
RGS12 (ENST00000344733)
VUS Missense
AA position
RGS RBD RGS12_usC RGS12_us1 PDZ RGS12_us2 GoLoco

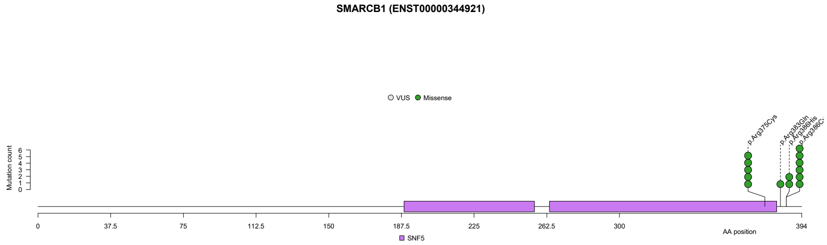
SMARCB1 (ENST00000344921)
VUS Missense
AA position
SNF5

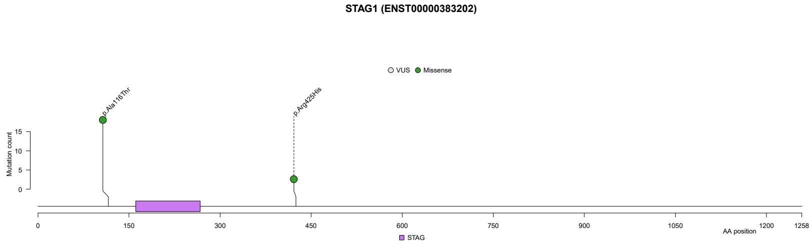
STAG1 (ENST00000383202)
VUS Missense
AA position
STAG

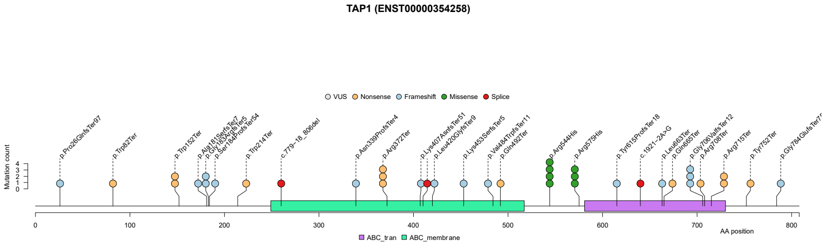
TAP1 (ENST00000354258)
VUS Nonsense Frameshift Missense Splice
AA position
ABC_tran ABC_membrane

**Supplementary Fig. 12: Non-synonymous mutations in selected driver genes identified in primary POL tumours.**

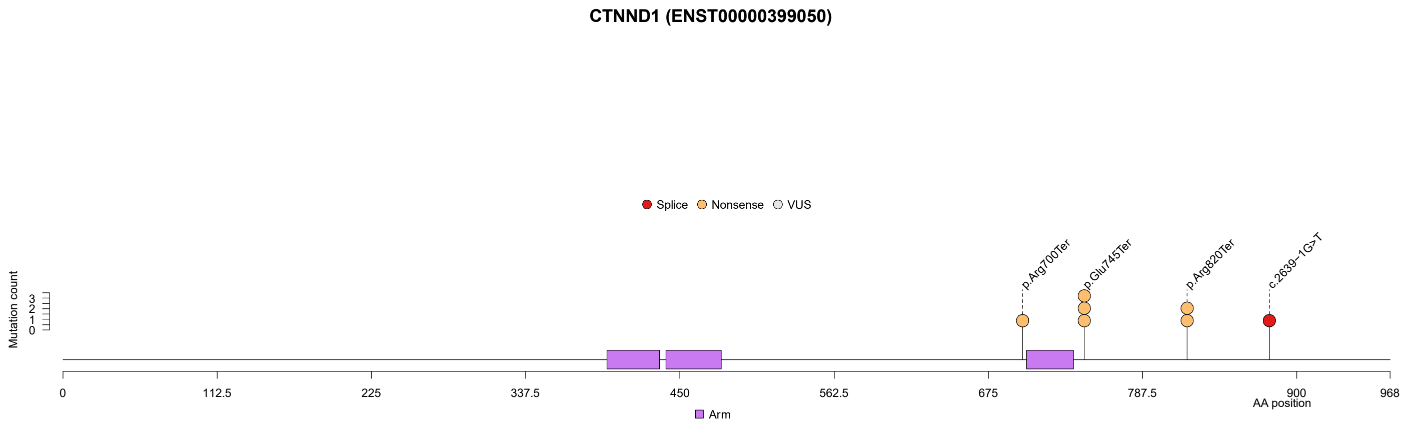

**Supplementary Fig. 13: Quantile-quantile plots of OncodriveFML results for non-coding elements in primary MSS tumours.** Red circles indicate non-coding elements with $Q<0.01$. The overdispersion of the test statistic for driver status revealed by the QQ plots suggests numerous non-coding drivers. However, we remain cautious about declaring driver status for any gene or element based on genomic data alone. For example, analysis of the *CSMD3* distal promoter in primary MSS samples revealed two recurrently mutated positions at chr8:113437866 (n=7, 0.5%) and chr8:113437869 (n=11, 0.7%). The chr8:113437866 hotspot exclusively comprised by GTT>GGT transversions. The chr8:113437869 hotspot was characterised by CTT>CGT (n=5), CTT>CCT (n=4) and CTT>CAT (n=2) substitutions. Each was respectively located 3bp and 1bp immediately upstream of a CTCF binding motif predicted by the JASPAR database [5] and supported by CTCF ChIP data in colon [6] . This is consistent with the significant functional impact signals found at the CTCF binding site overlapping the hotspots (chr8:113437400-113438400). In line with previous observations, five of 12 CTT>CGY and three of four CTT>CCT mutations were attributable to SBS17b and SBS17a, respectively [7,8] . CTCF binding sites are known to be enriched in mutations, creating passenger mutation clusters and hotspots [9]). Based on this, we cannot exclude the possibility that these two hotspots in the *CSMD3* distal promoter are passenger mutations.

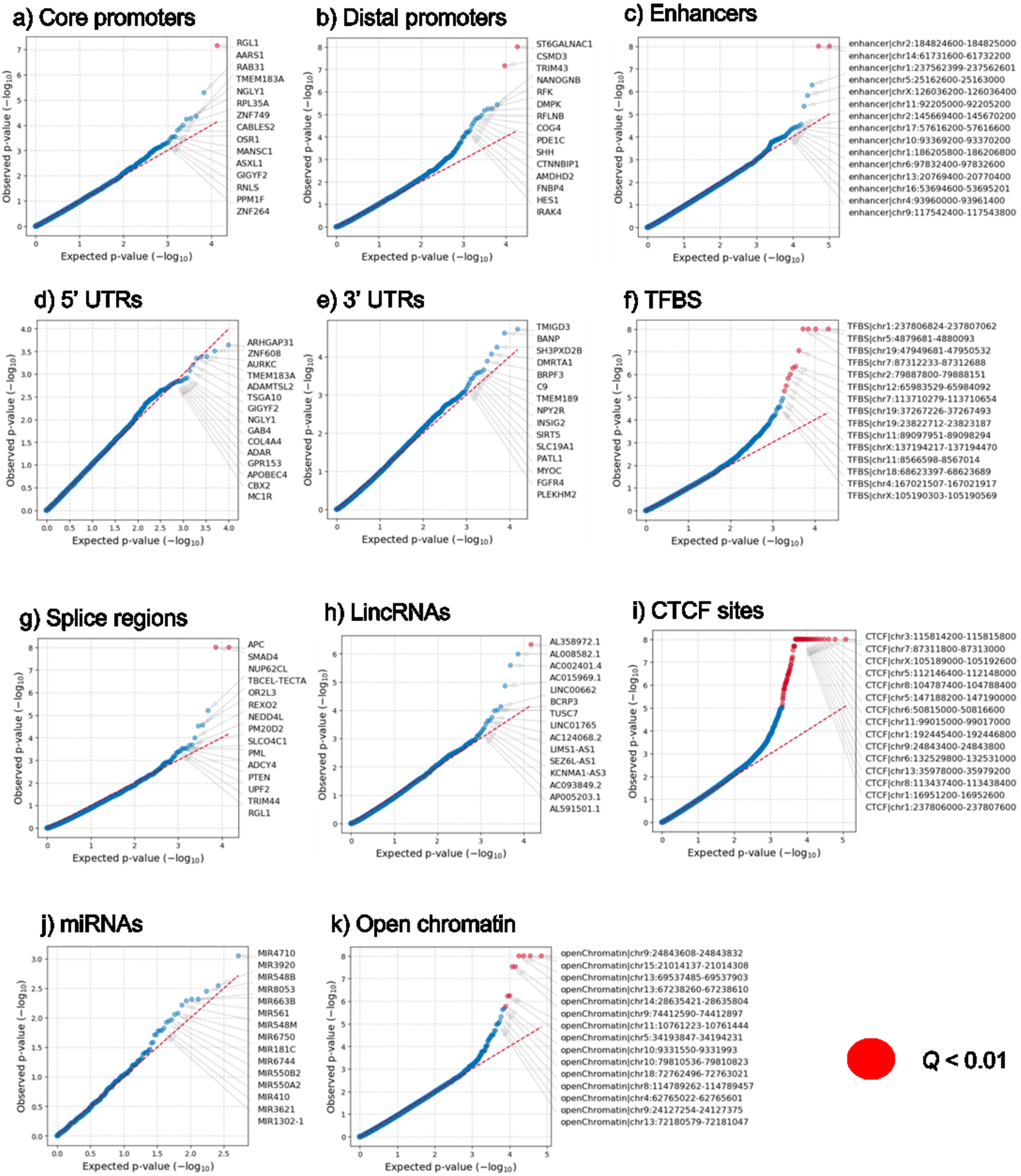

a) Core promoters
Observed p-value (−log10)
Expected p-value (−log10)
RGL1
AARS1
RAB31
TMEM183A
NGLY1
RPL35A
ZNF749
CABLES2
OSR1
MANSC1
ASXL1
GIGYF2
RNLS
PPM1F
ZNF264
b) Distal promoters
ST6GALNAC1
CSMD3
TRIM43
NANOGNB
RFK
DMPK
RFLNB
COG4
PDE1C
SHH
CTNNBIP1
AMDHD2
FNBP4
HES1
IRAK4
c) Enhancers
enhancer|chr2:184824600-184825000
enhancer|chr14:61731600-61732200
enhancer|chr1:237562399-237562601
enhancer|chr5:25162600-25163000
enhancer|chrX:126036200-126036400
enhancer|chr11:92205000-92205200
enhancer|chr2:145669400-145670200
enhancer|chr17:57616200-57616600
enhancer|chr10:93369200-93370200
enhancer|chr1:186205800-186206800
enhancer|chr6:97832400-97832600
enhancer|chr13:20769400-20770400
enhancer|chr16:53694600-53695201
enhancer|chr4:93960000-93961400
enhancer|chr9:117542400-117543800
d) 5' UTRs
ARHGAP31
ZNF608
AURKC
TMEM183A
ADAMTSL2
TSGA10
GIGYF2
NGLY1
GAB4
COL4A4
ADAR
GPR153
APOBEC4
CBX2
MC1R
e) 3' UTRs
TMIGD3
BANP
SH3PXD2B
DMRTA1
BRPF3
C9
TMEM189
NPY2R
INSIG2
SIRT5
SLC19A1
PATL1
MYOC
FGFR4
PLEKHM2
f) TFBS
TFBS|chr1:237806824-237807062
TFBS|chr5:4879681-4880093
TFBS|chr19:47949681-47950532
TFBS|chr7:87312233-87312688
TFBS|chr2:79887800-79888151
TFBS|chr12:65983529-65984092
TFBS|chr7:113710279-113710654
TFBS|chr19:37267226-37267493
TFBS|chr19:23822712-23823187
TFBS|chr11:89097951-89098294
TFBS|chrX:137194217-137194470
TFBS|chr11:8566598-8567014
TFBS|chr18:68623397-68623689
TFBS|chr4:167021507-167021917
TFBS|chrX:105190303-105190569
g) Splice regions
APC
SMAD4
NUP62CL
TBCEL-TECTA
OR2L3
REXO2
NEDD4L
PM20D2
SLCO4C1
PML
ADCY4
PTEN
UPF2
TRIM44
RGL1
h) LincRNAs
AL358972.1
AL008582.1
AC002401.4
AC015969.1
LINC00662
BCRP3
TUSC7
LINC01765
AC124068.2
LIMS1-AS1
SEZ6L-AS1
KCNMA1-AS3
AC093849.2
AP005203.1
AL591501.1
i) CTCF sites
CTCF|chr3:115814200-115815800
CTCF|chr7:87311800-87313000
CTCF|chrX:105189000-105192600
CTCF|chr5:112146400-112148000
CTCF|chr8:104787400-104788400
CTCF|chr5:147188200-147190000
CTCF|chr6:50815000-50816600
CTCF|chr11:99015000-99017000
CTCF|chr1:192445400-192446800
CTCF|chr9:24843400-24843800
CTCF|chr6:132529800-132531000
CTCF|chr13:35978000-35979200
CTCF|chr8:113437400-113438400
CTCF|chr1:16951200-16952600
CTCF|chr1:237806000-237807600
j) miRNAs
MIR4710
MIR3920
MIR548B
MIR8053
MIR663B
MIR561
MIR548M
MIR6750
MIR181C
MIR6744
MIR550B2
MIR550A2
MIR410
MIR3621
MIR1302-1
k) Open chromatin
openChromatin|chr9:24843608-24843832
openChromatin|chr15:21014137-21014308
openChromatin|chr13:69537485-69537903
openChromatin|chr13:67238260-67238610
openChromatin|chr14:28635421-28635804
openChromatin|chr9:74412590-74412897
openChromatin|chr11:10761223-10761444
openChromatin|chr5:34193847-34194231
openChromatin|chr10:9331550-9331993
openChromatin|chr10:79810536-79810823
openChromatin|chr18:72762496-72763021
openChromatin|chr8:114789262-114789457
openChromatin|chr4:62765022-62765601
openChromatin|chr9:24127254-24127375
openChromatin|chr13:72180579-72181047
Q < 0.01

**Supplementary Fig. 14: Hotspots of simple structural variants (SVs) identified in primary microsatellite unstable cancers (*n*=292).** Coloured lines represent numbers of samples with a SV breakpoint of each class in 1Mb genome regions. Hotspots are annotated with their cytoband, the number of contained genes (in brackets) and any candidate gene. SVs at fragile sites are not included.

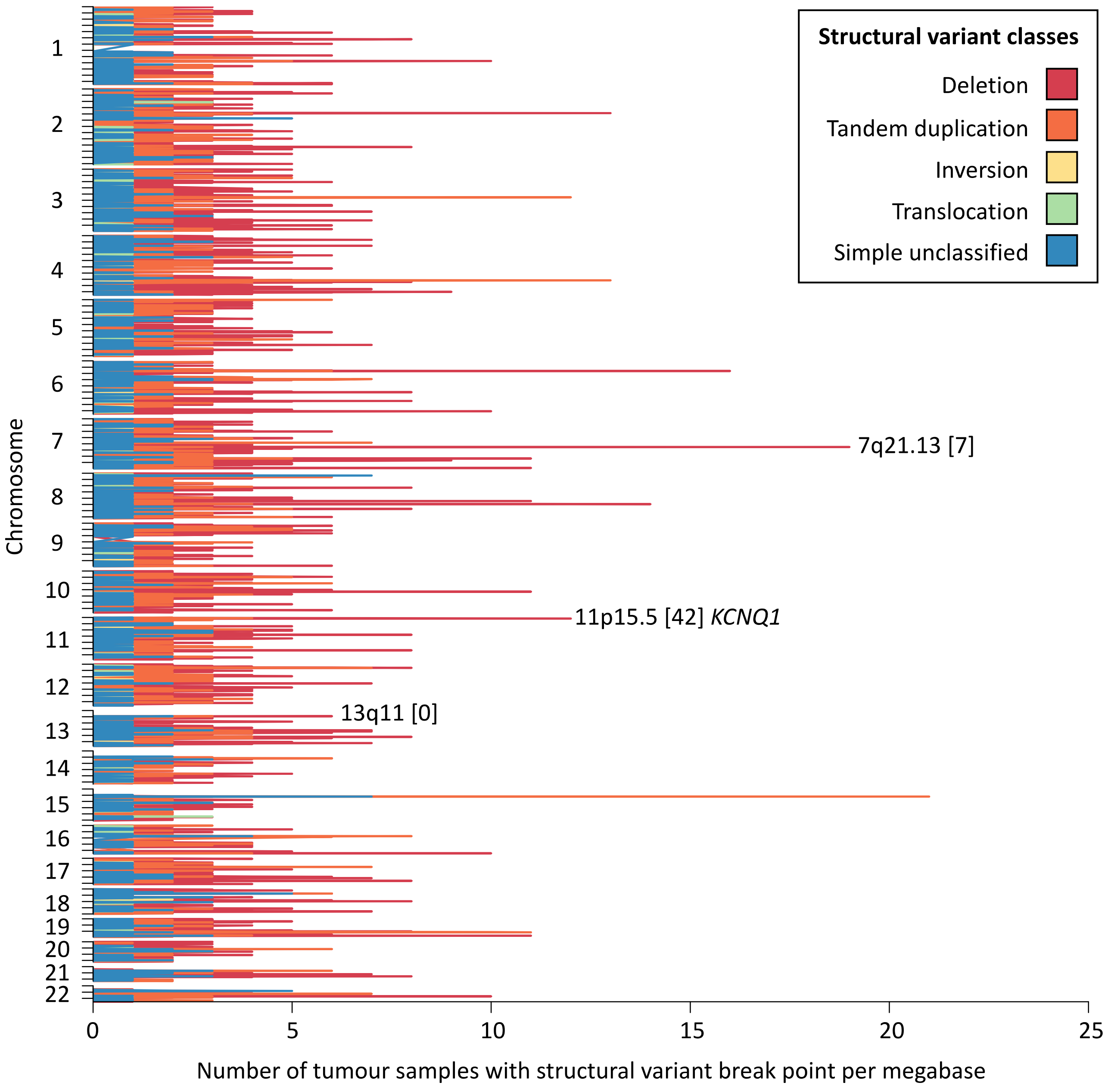

**Supplementary Fig. 15. Hotspots of simple structural variants (SVs) identified at 17q24.3 overlapping a *SOX9* promoter-interacting regulatory element in primary microsatellite stable (MSS) cancers (*n*=1354).** Upper track arcs represent simple SVs. Second-from-top track shows mean GC-corrected log ratio between tumour and normal read coverage (logR) as computed by Battenberg (Nik-Zainal *et al.*, 2012, Cell). Higher and lower logR values indicate tendencies for copy number gains and losses amongst the considered tumour samples. Third-from-top track shows chromosomal interactions identified in HT29 cells using promoter capture Hi-C (Orlando *et al.*, 2018, Nature Genetics). Fourth-from-top shows histone mark signals. Vertical lines represent hotspot start and end positions.

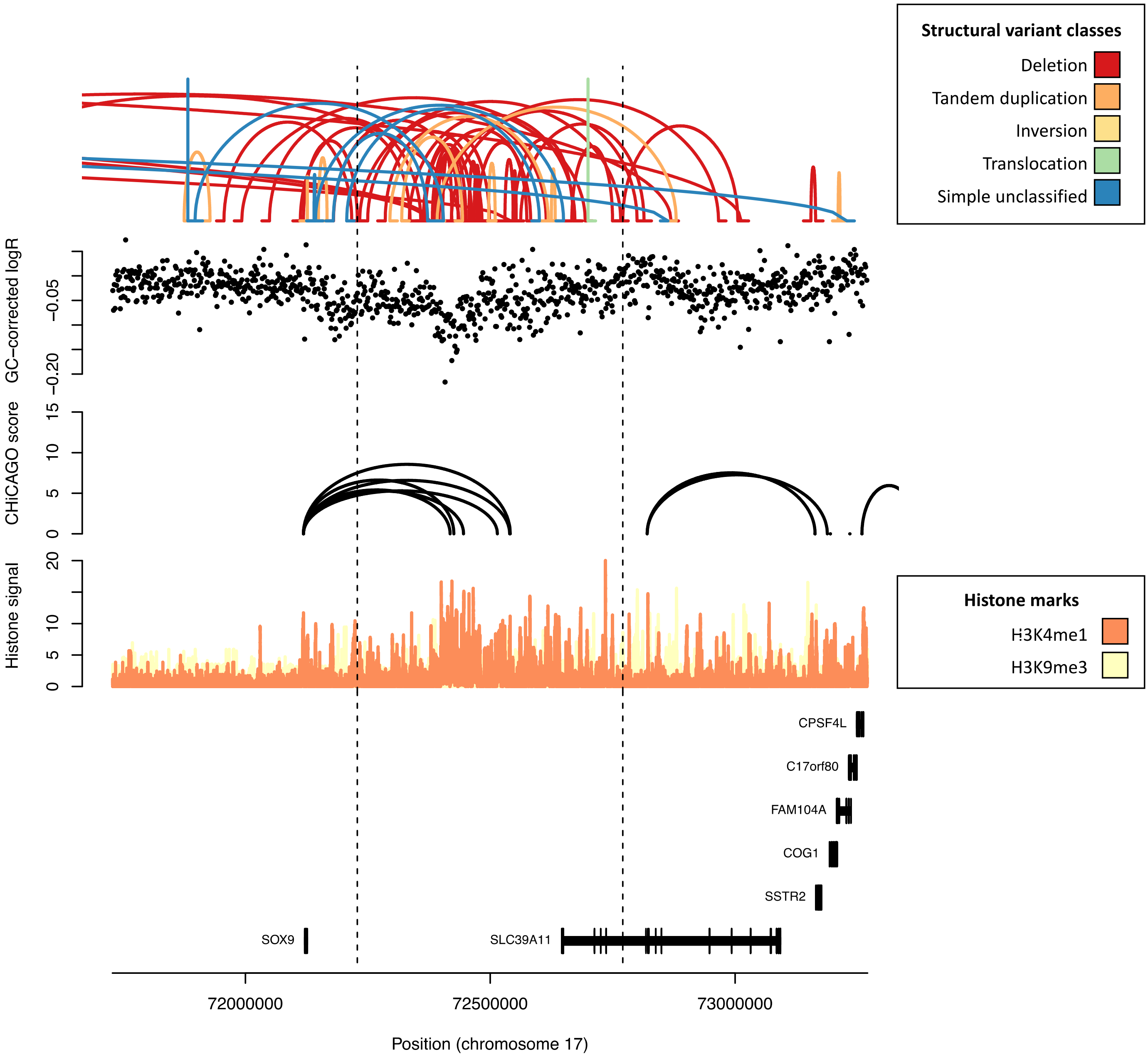

**Supplementary Fig. 16: Classification of tumour samples into diploid and tetraploid (genome-doubled), upper. Hierarchical clustering of all CRCs based on copy number states identifies WGD/non-WGD split, and suggests two WGD subtypes, lower**. WGD classification method is described in **Online Methods**.

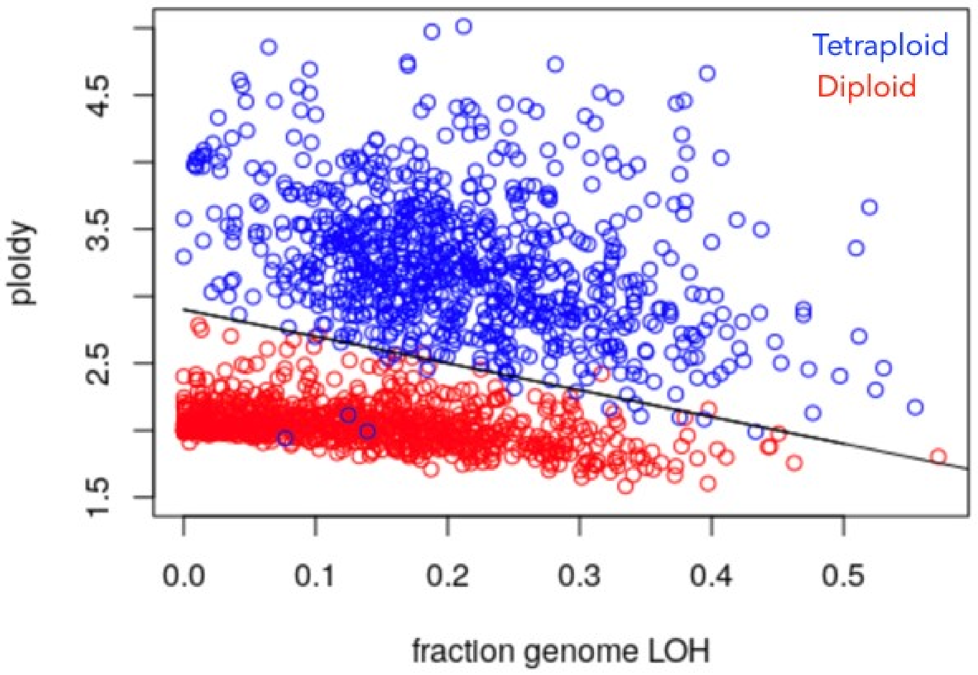


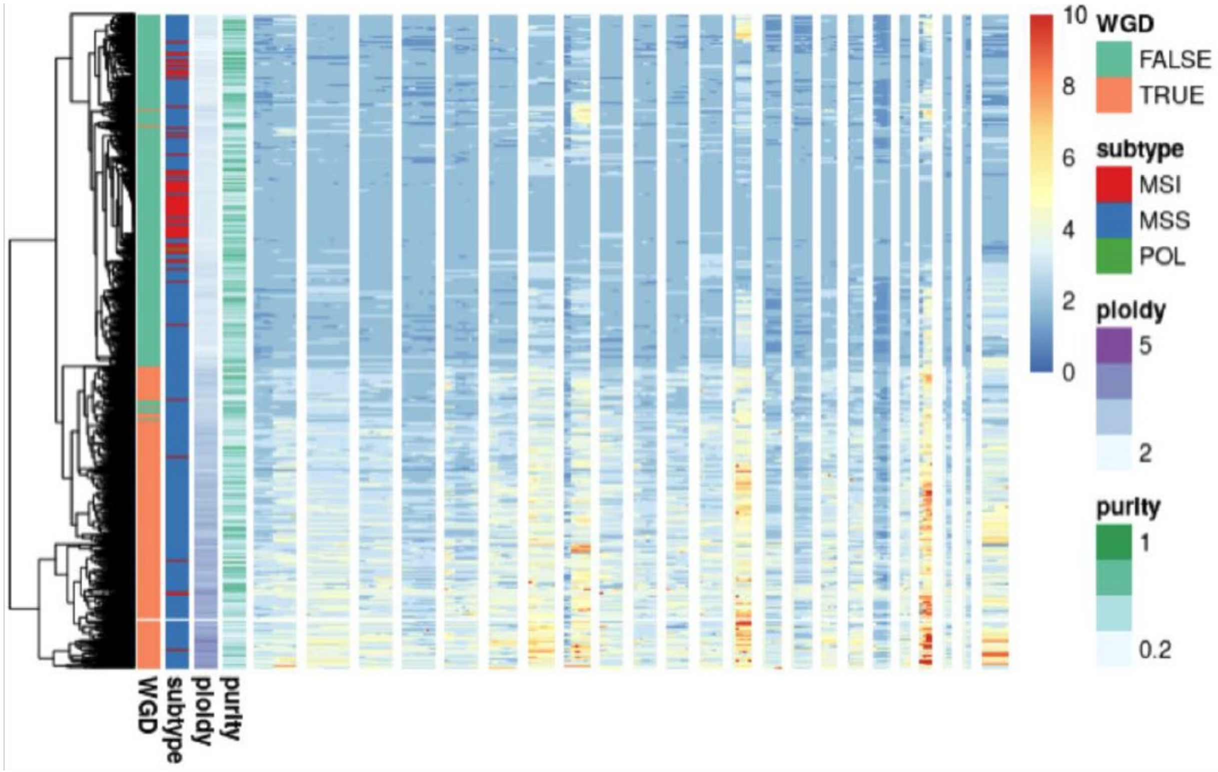

**Fig. 17. Number of copies gained in primary MSS cancers.** Shown are chromosome arms with recurrent arm-level gains identified by GISTIC ($Q < 0.05$).

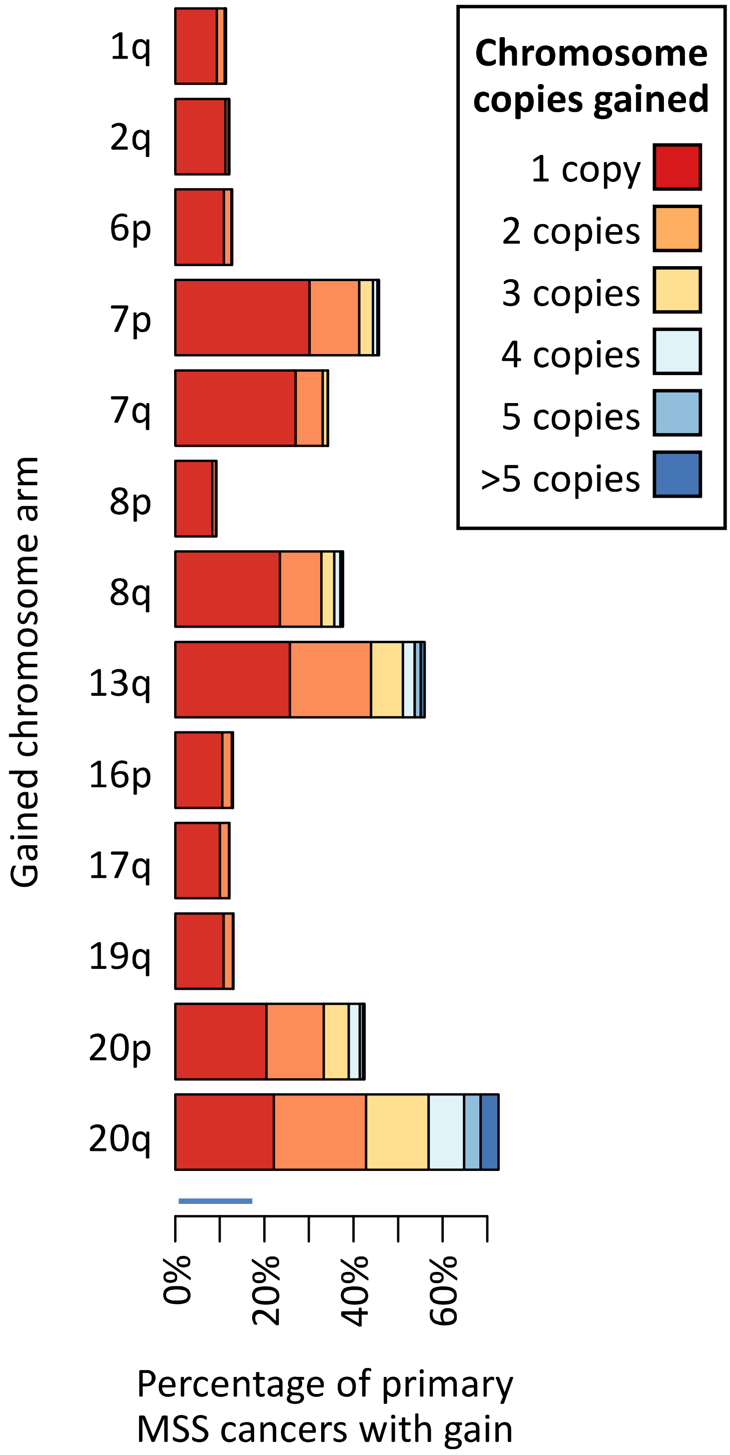

**Supplementary Fig. 18. Extrachromosomal DNA (ecDNA) detected across CRC subtypes and its contribution to common oncogene amplification.** A) Counts of tumours carrying at least one ecDNA amplicon across tumour subtypes (e.g. tumour counted as “Circular” if ≥ 1 circularised amplicon detected, otherwise “BFB” if ≥ 1 BFB amplicon detected until “No amp” where no valid amplicon detected). (B-D) ecDNA classification of commonly amplified oncogenes in Primary MSS, Primary MSI and Metastases MSS tumours respectively. Classification was restricted to gene amplifications with a total copy number ≥ 5 in diploid tumours and ≥ 10 in tetraploid tumours (i.e. “big gains”). Primary POL cohort not shown as no genes were amplified. Counts taken from Supplementary Table 29.

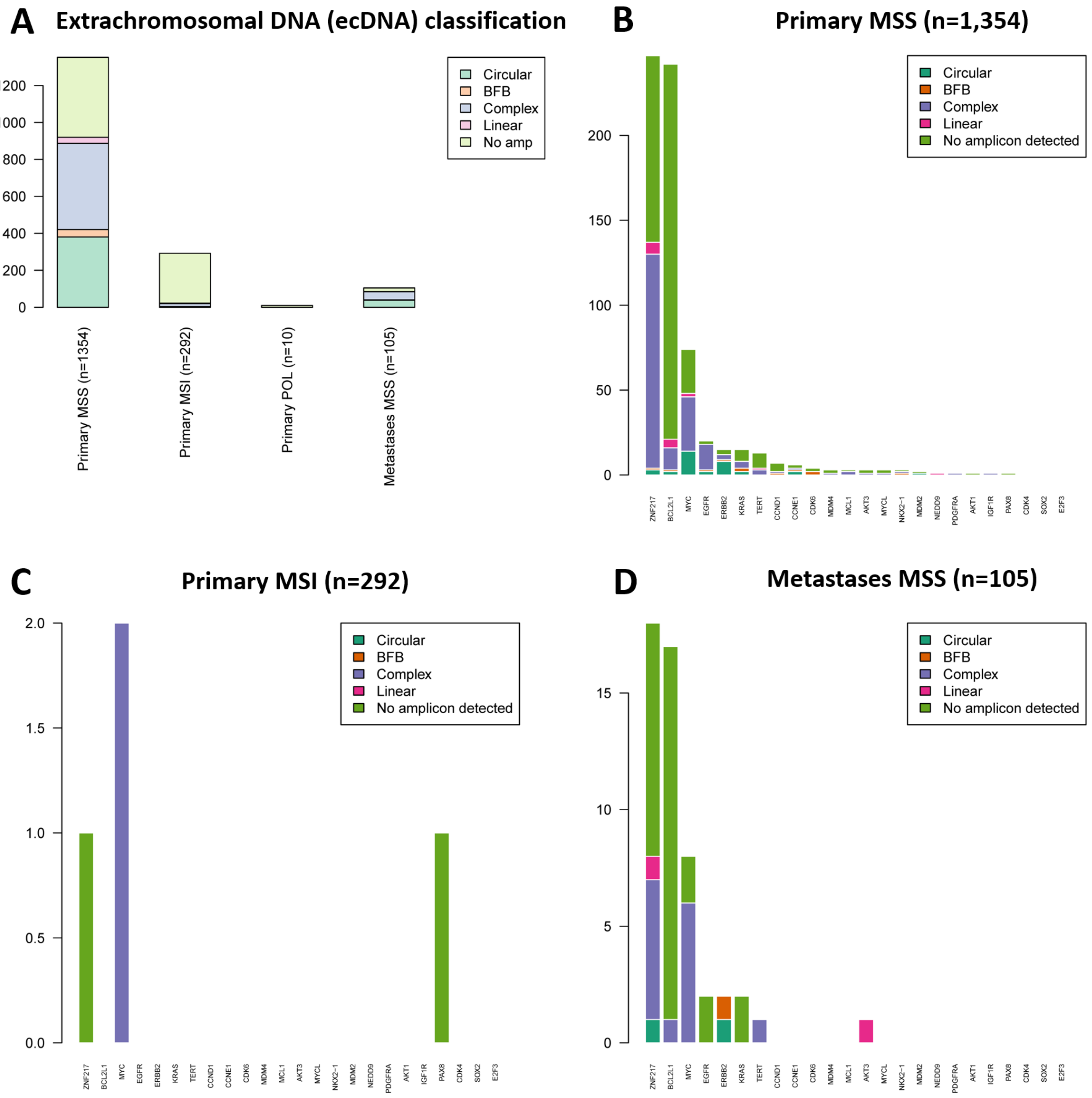

**Supplementary Fig. 19: Distributions of mitochondrial DNA (mtDNA) mutations**. Number of SNVs of the 12 substitution classes with light strand as reference. The data indicate strand bias in the mutational processes as regards: (**a**) heavy and light mtDNA strands; (**b**) transcribed and non-transcribed strands of mitochondrial protein-coding genes; (**c**) transcribed and non-transcribed strands of protein-coding genes transcribed from the heavy strand; (**d**) transcribed and non-transcribed strands of protein-coding genes transcribed from the light strand; and (**e**) heavy and light strands between the two origin of replication sites. In each case, the strand assignment is nominal since the strand that suffered the mutation cannot be determined.

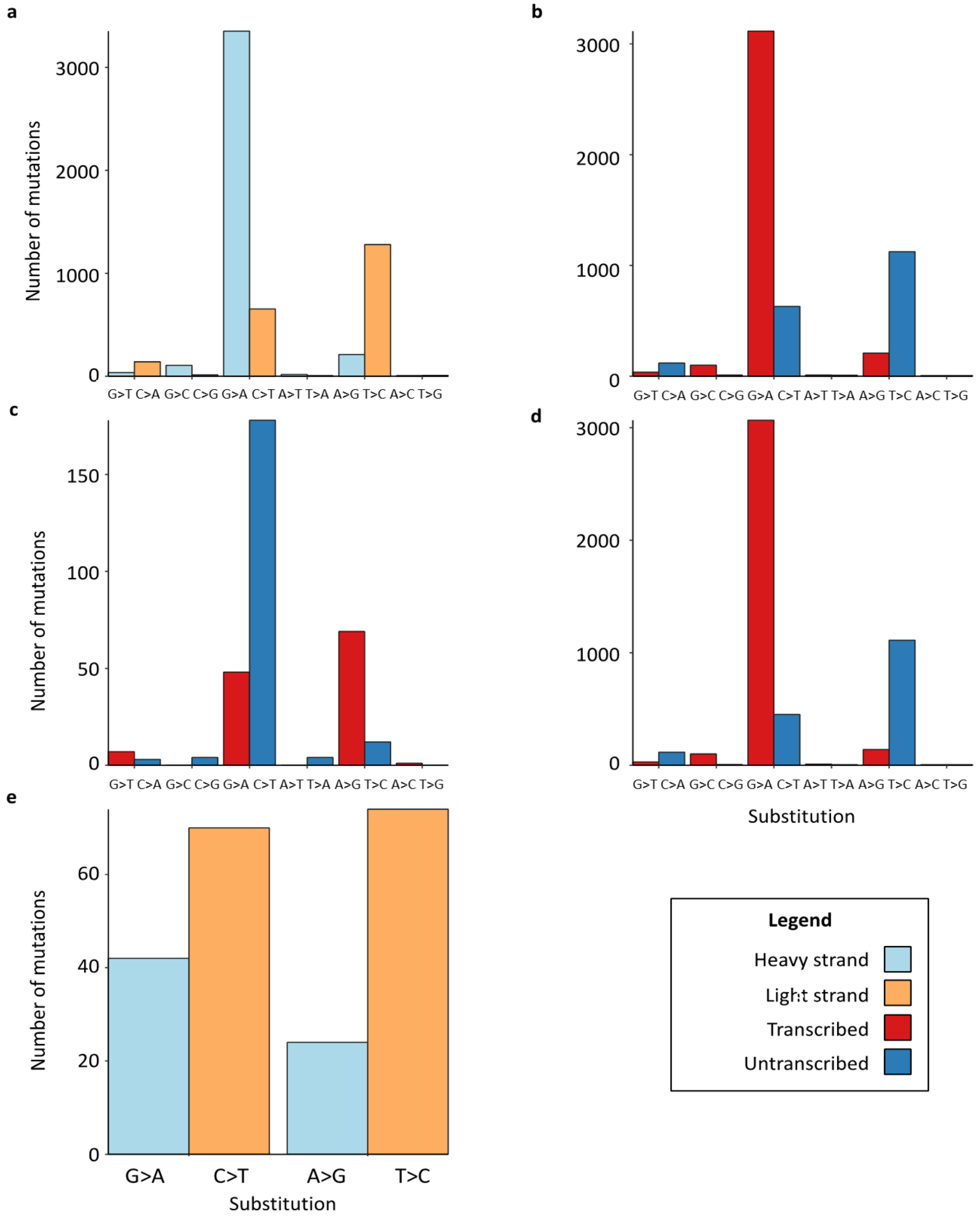

**Supplementary Fig. 20. Neoantigen burden metric comparisons**. Lower and upper triangle and diagonal cells represent pairwise scatter plots, correlation statistics and density plots respectively (green: MSS cancers; red: MSI; yellow: POL). NA mutation: number of unique mutations giving rise to one or more neoantigens; NA peptide: number of unique neoantigen peptides; NA total: number of antigenic HLA-peptide interactions detected; Expr NA total: number of antigenic HLA-peptide interactions detected in genes expressed in ≥10% TCGA CRCs. Measures presented on log10 scale.

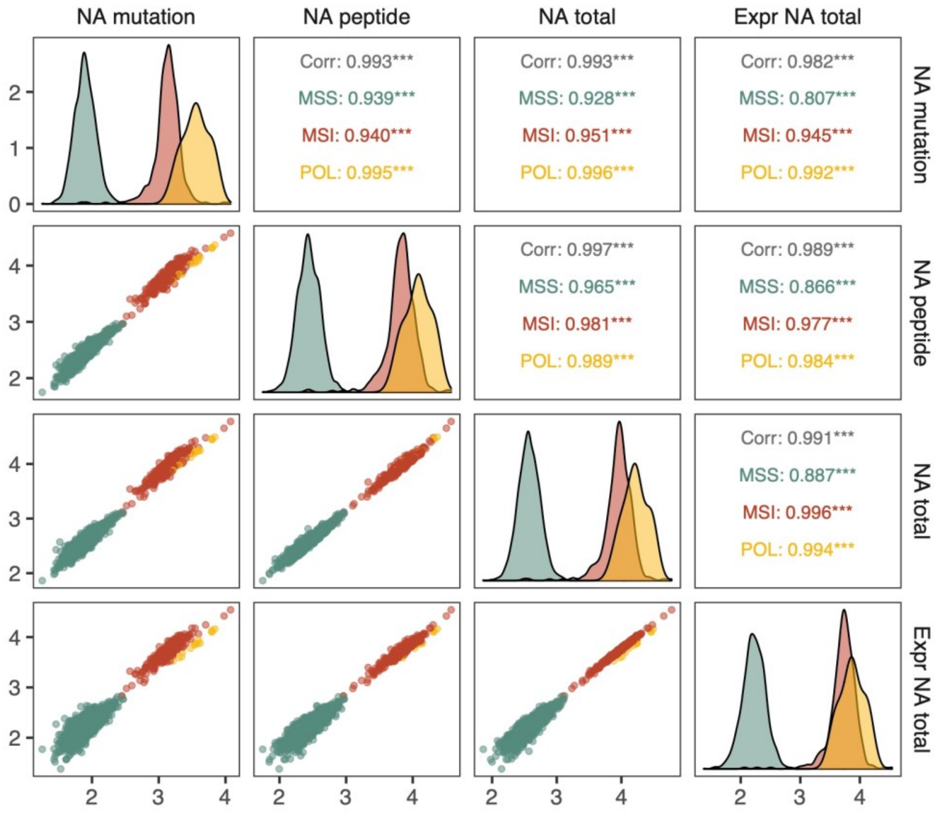

**Supplementary Fig. 21. Heatmap and frequency chart of the 20 most common antigenic (a) SNV and (b) frameshift mutations.** Mutations are shown in order of decreasing frequency across the CRC set. Colours indicate if the mutation is antigenic (dark blue), has escaped antigenicity through HLA alteration (purple) or is non-antigenic (light blue). The molecular subtype of each cancer is shown above the heatmap (green: MSS, red: MSI, yellow: POL, pink: POL&MSI). Frameshift mutations are shown for MSI cancers only.

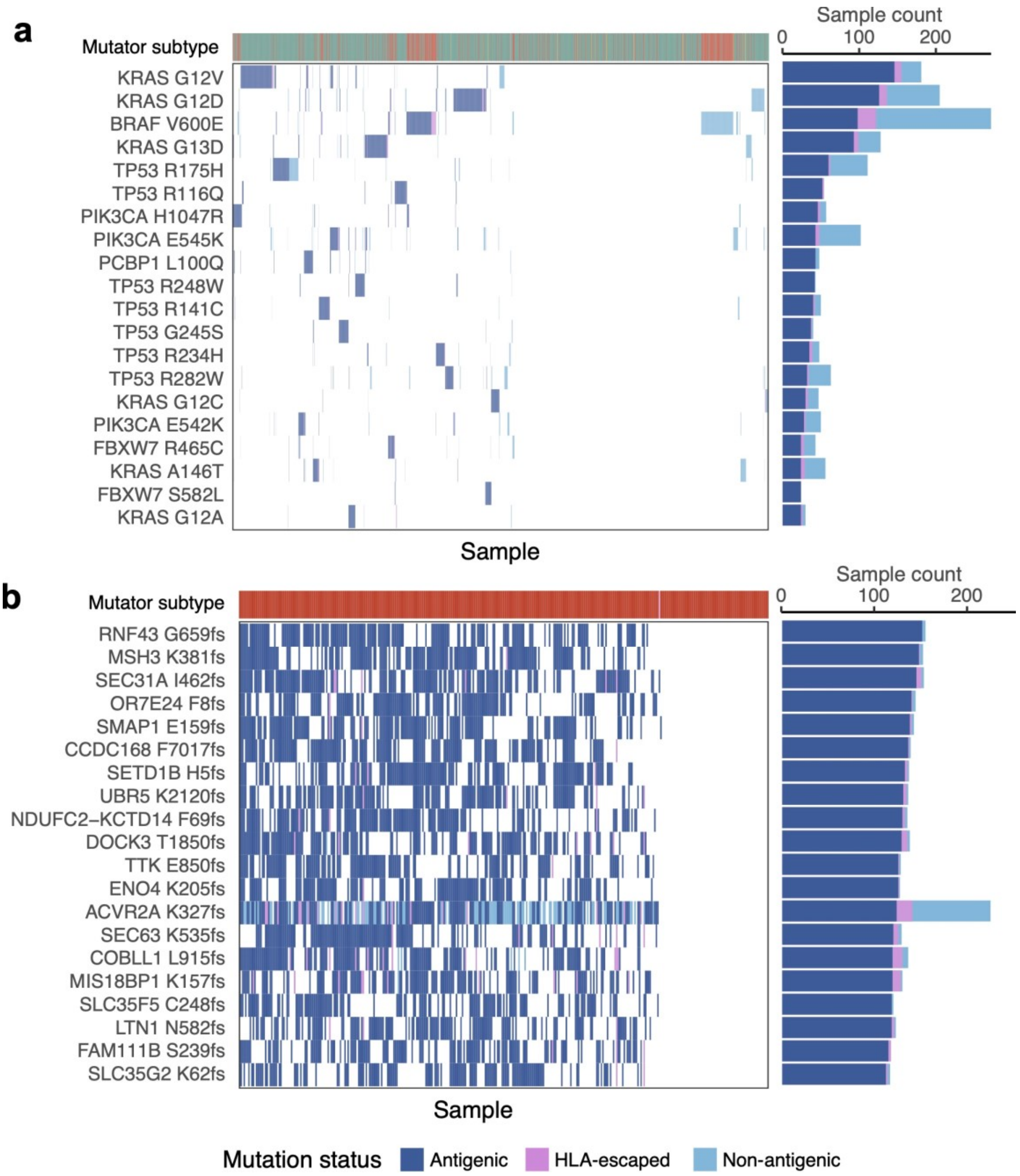

**Supplementary Fig. 22. Immunogenic potential of CRC driver mutations.** (**a**) Median PHBR of driver mutations across the whole CRC cohort, plotted against the number of cancers the mutations are observed in. Grey dots represent individual mutations, red dots show the median for mutations at the same frequency. Correlation coefficient is computed using Spearman's rank correlation. (**b**) Median PHBR of driver mutations that are shared between CRC subtypes, computed separately for cancers of each subtype. Lines connect PHBR values of the same mutation across subtypes. P-values of paired Wilcoxon tests comparing primary MSS to other subtypes are shown above. (**c**) PHBR of all mutations in all cancers that were not observed (in grey) compared to mutations that were observed (in red).

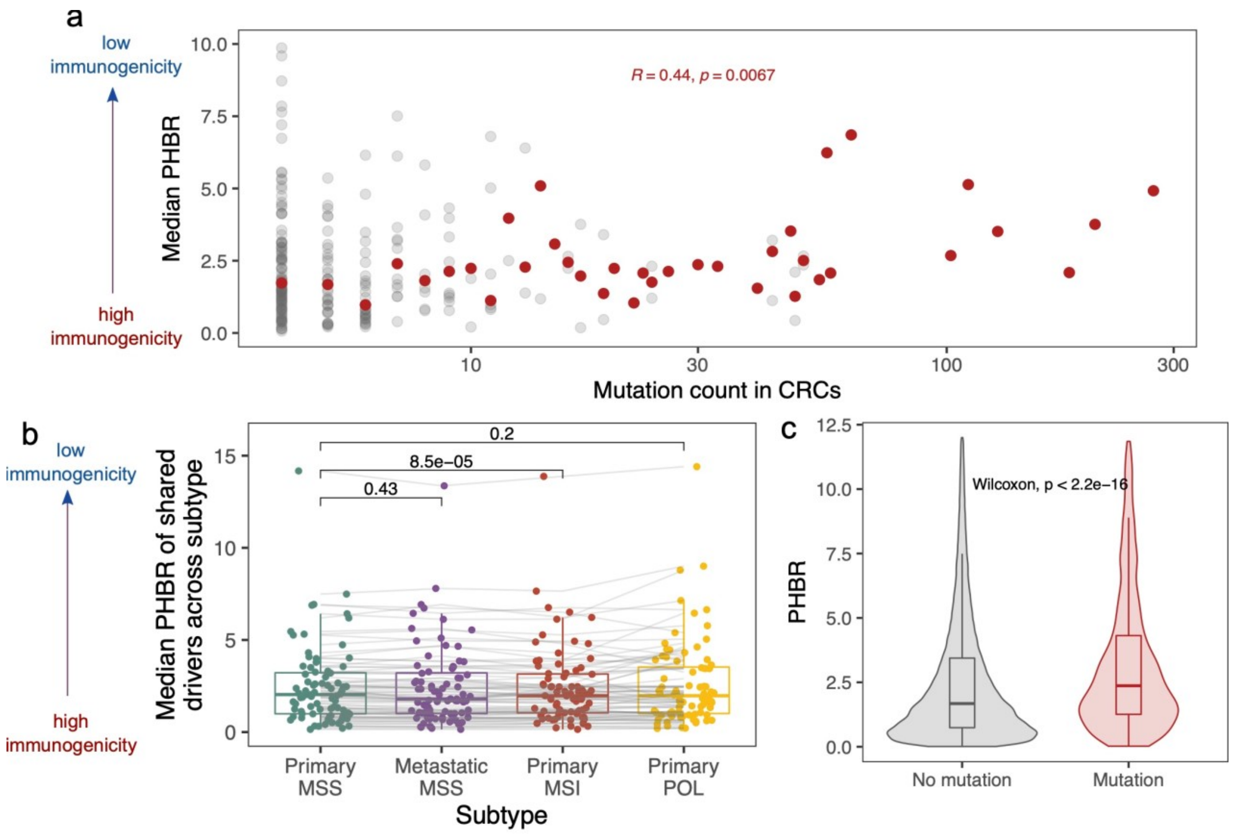

**Supplementary Fig. 23. Effect of immune escape on neoantigen burden in a multiple regression model including clinical variables.** Green and red squares represent neoantigen burden fold changes in MSS and MSI cancers respectively, with whiskers showing 95% confidence intervals. Escape is defined as having HLA LOH or a mutation in the HLA or other antigen presenting genes (see Online Methods). Fold change is shown compared to Non-escaped, Male, Stage A/B, Proximal colon, Primary, untreated samples.

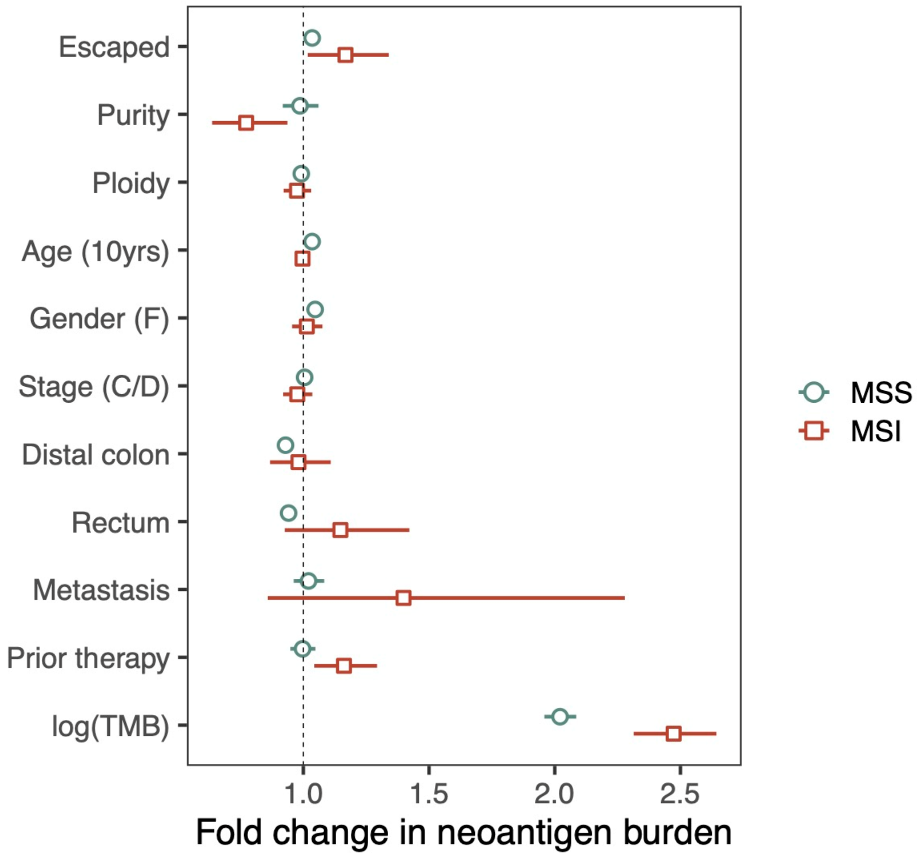

**Supplementary Fig. 24: Site of primary tumour and number of variants attributed to mutational signatures in primary microsatellite stable (MSS) tumours.** Shown are mutational signatures associated with age at sampling (10-year bins) at a Bonferroni-corrected *P*-value of 0.05 using multiple linear regression considering age at sampling, sex, primary tumour site, stage, grade and sample purity. *n*: number of tumour samples from site.

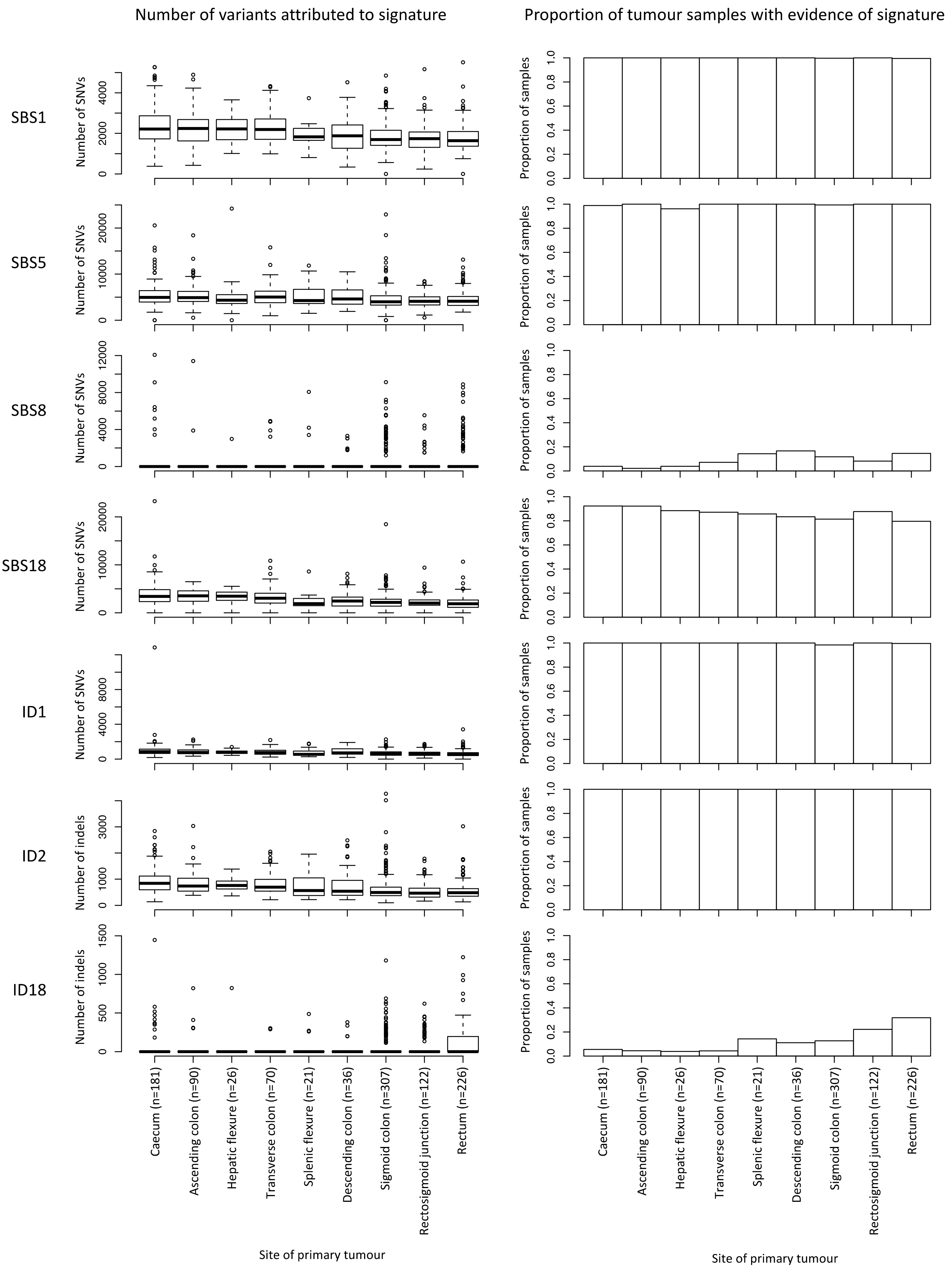

**Supplementary Fig. 25: Signature activity in normal crypt epithelial cells from the right, transverse and left colon.** Where multiple crypts from the same colon region had been sampled in a single participant, the median number and proportion of variants attributed to each signature was considered. Data from Lee-Six *et al.* (2019, Nature). IDA closely resembles ID18. P-values were computed using two-sided Wilcoxon rank sum tests. *n*: number of participants, Trans: transverse colon.

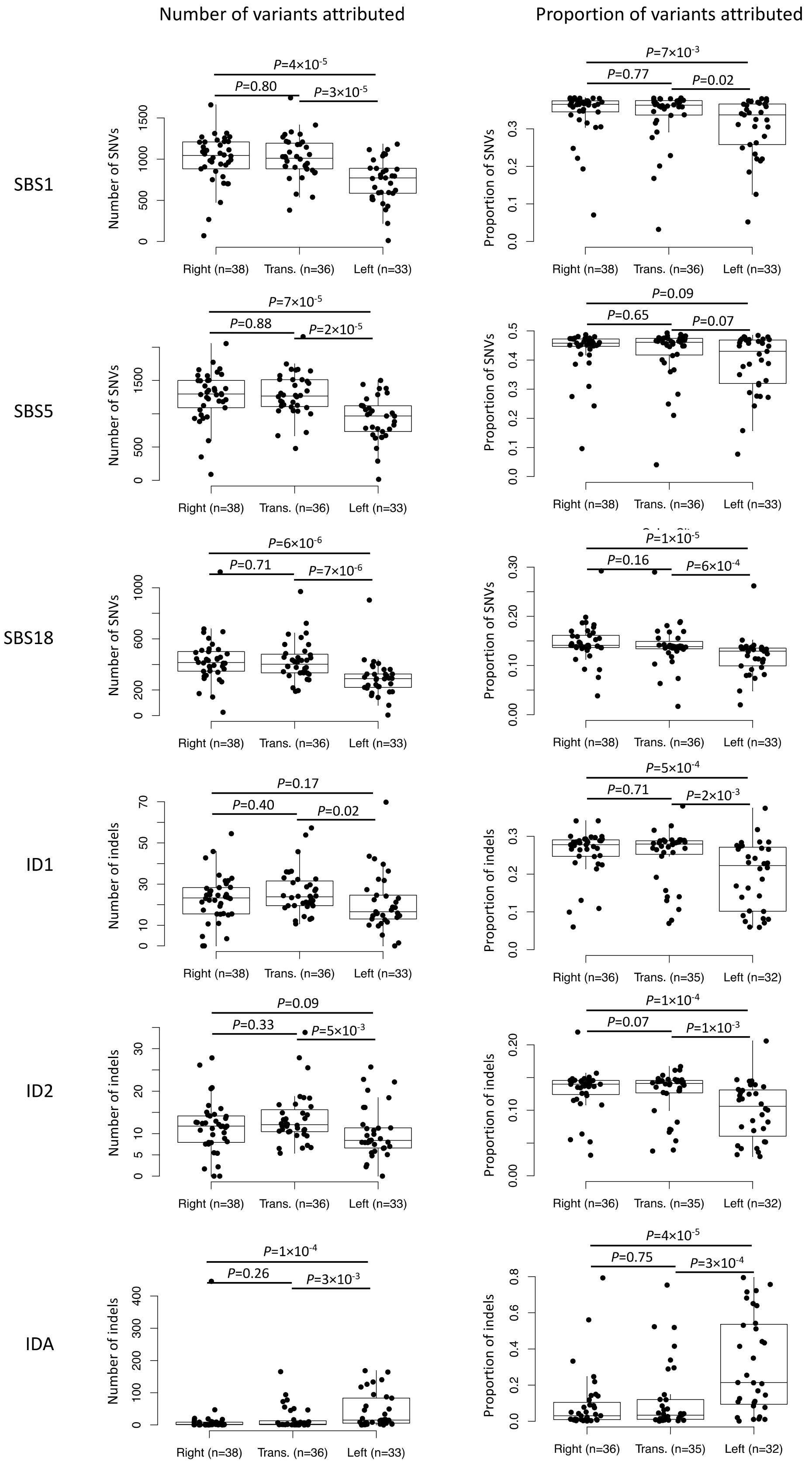

**Supplementary Fig. 26: Age at sampling and number of variants attributed to mutational signatures in primary microsatellite stable (MSS) tumours.** Shown are mutational signatures associated with age at sampling (10 year bins) at a Bonferroni-corrected *P*-value of 0.05 using multiple linear regression considering age at sampling, sex, primary tumour site, stage, grade and sample purity. The Yeo-Johnson extension to the Box-Cox-transformation was applied to variant numbers.

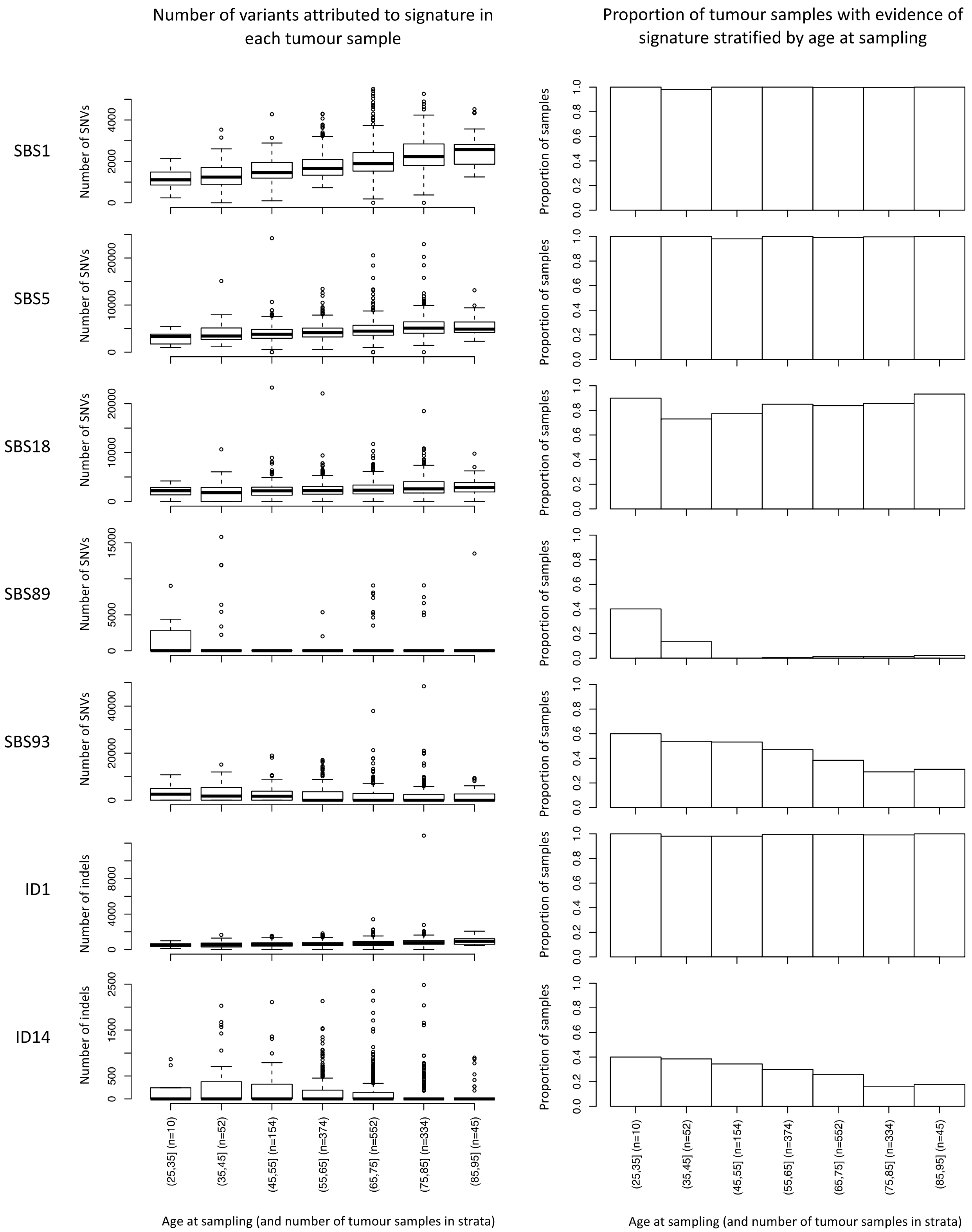

**Supplementary Fig. 27. Genomic comparison of primary and metastatic MSS colorectal cancer. (a)** Total mutation burden was not significantly different between primary and metastatic tumours. **(b)** Tumour ploidy was significantly increased in metastatic tumours. **(c)** Trace along the genome showing, for each 1Mb window, the difference in proportion of mono-allelic lossGain/LOH between metastatic and primary tumours. Black stars indicate significance controlled for multiple testing: * FDR<0.05, ** FDR<0.01, *** FDR<0.001. **(d)** Trace along the genome showing, for each 1Mb window, the difference in mean ploidy-adjusted copy number between metastatic and primary tumours. Red stars indicate significance controlled for multiple testing: * FDR<0.05, ** FDR<0.01, *** FDR<0.001.

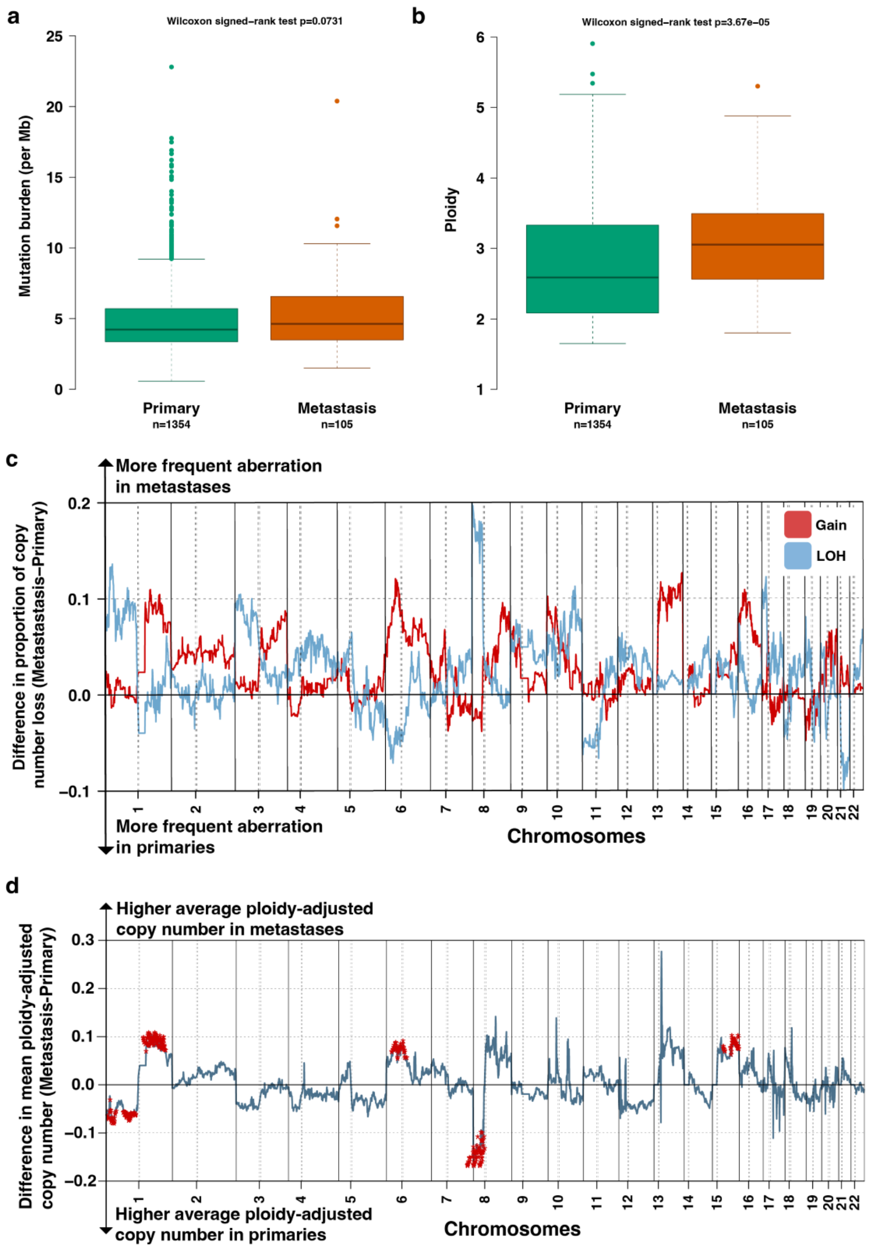

**Supplementary Fig. 28. Immune comparison of primary and metastatic MSS colorectal cancer. (a)** The proportion of samples that were immune-escaped was not significantly different between primary and metastatic tumours. **(b)** The total number of neoantigens per sample was not significantly different between primary and metastatic tumours.

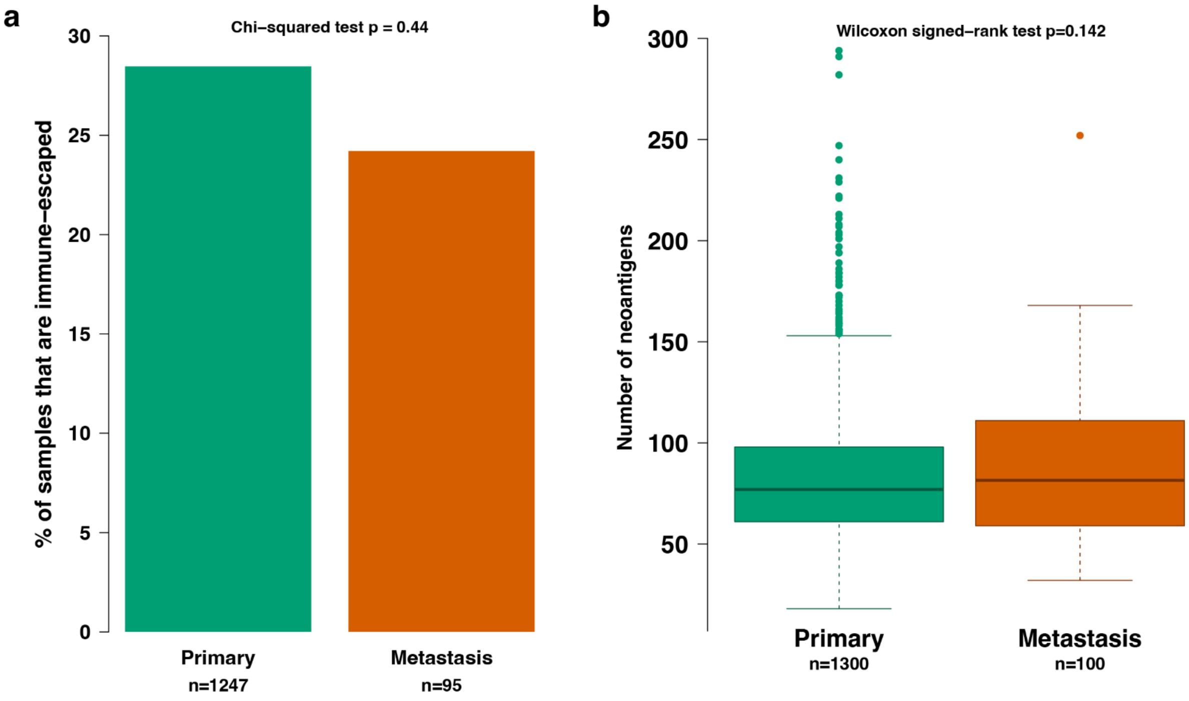

**Fig. 29. Microbiome decontamination process.** Tumour and blood prevalence of all species are shown, according to methods based on The Cancer Microbiome Atlas. Orange points indicate taxa thought to be contaminants due to presence in both blood and tumour samples. Outlined points indicate species previously associated with CRC.

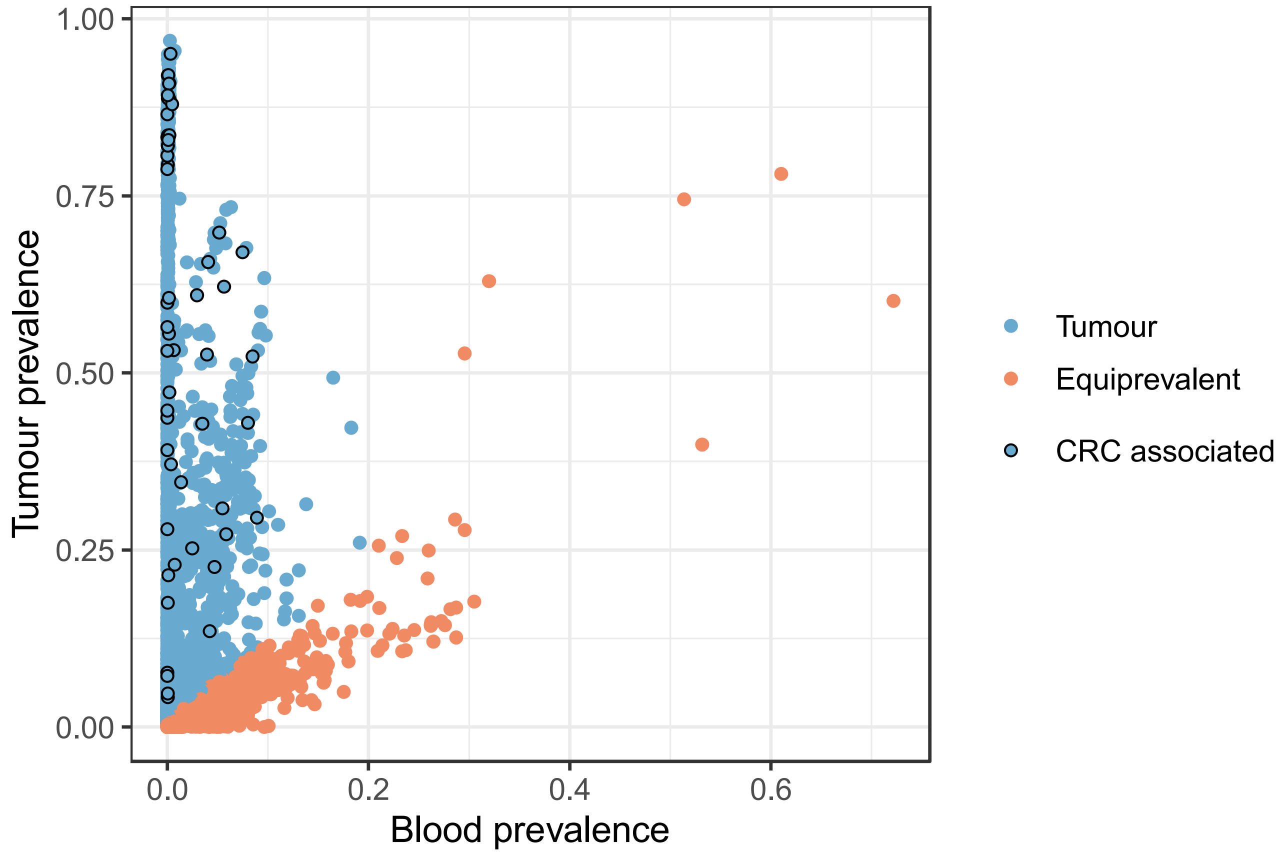

**Supplementary Fig. 30. Mean relative abundance of microbial genera for the four main CRC subgroups.**

The most abundant 20 genera are shown. Other taxa are summed as “Others” for ease of visualisation.

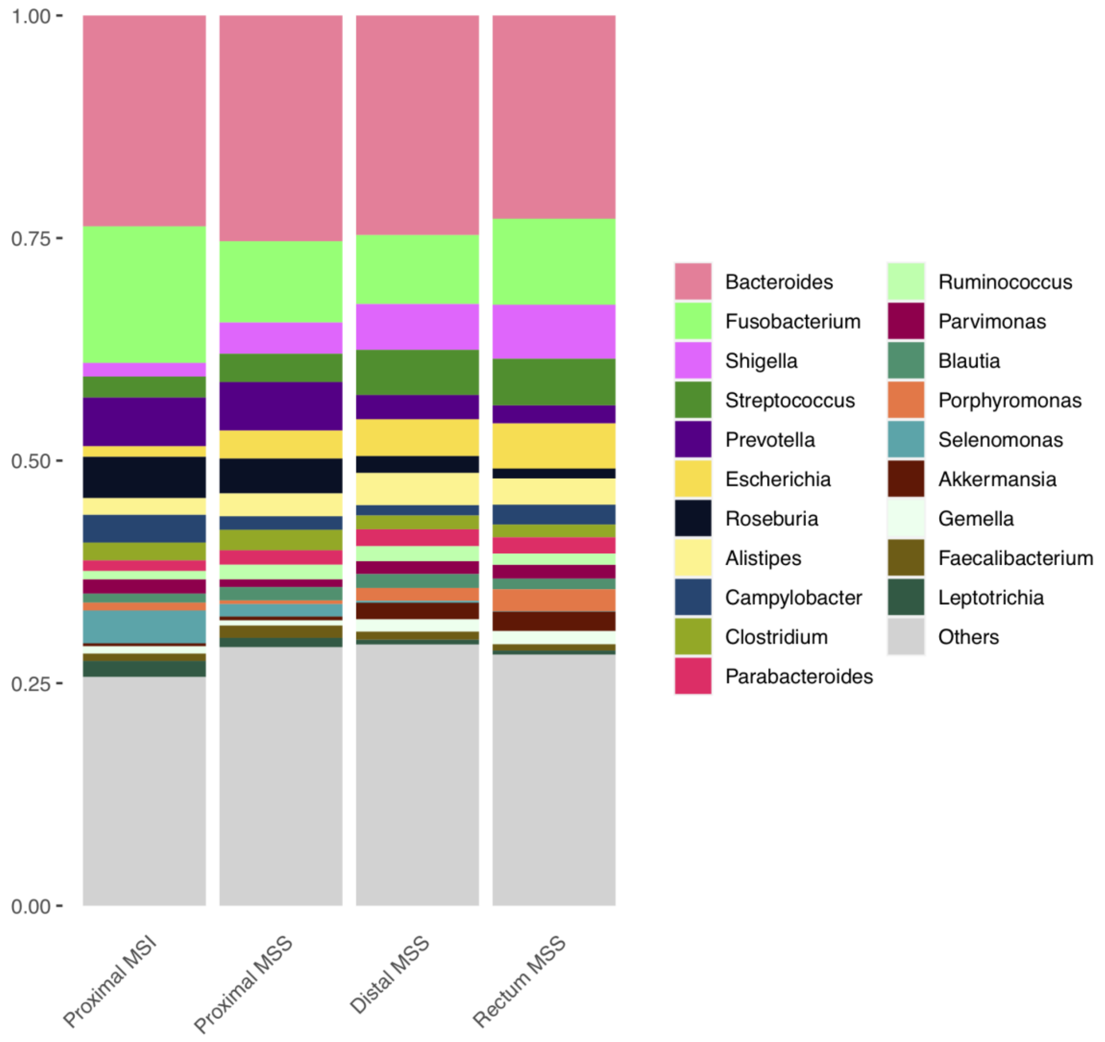

**Supplementary Fig. 31. Bacterial load (A) and Shannon diversity index (B) for different CRC groupings.** Primary tumours are compared to metastases. Within primary tumours, microsatellite instability (MSI) status and anatomy are compared. The 33 distal and rectal MSI cancers are not included as the small cohort sizes do not allow meaningful comparisons. Mann-Whitney P-values for pairwise comparisons are displayed.

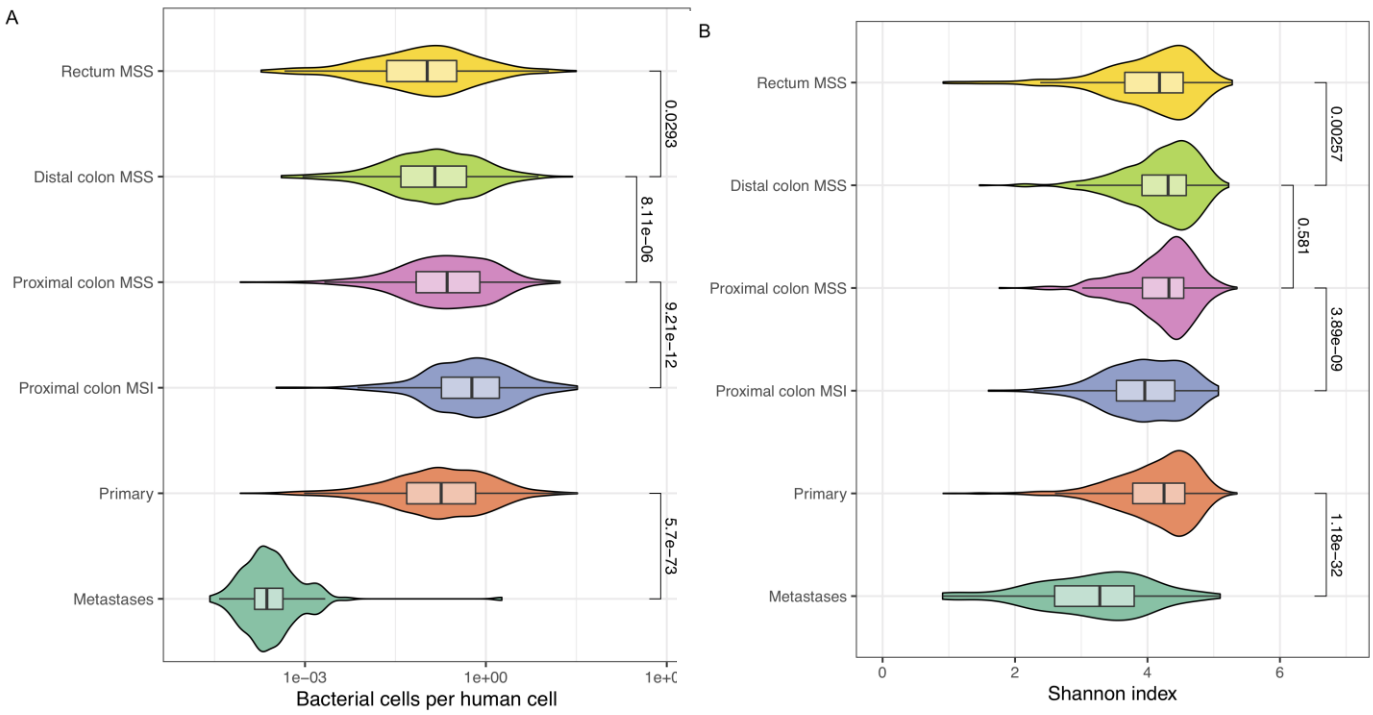

**Supplementary Fig. 32. Adonis PERMANOVA results comparing Bray-Curtis distances against various clinical and genomic factors.** R-squared is the percentage of diversity linked to each factor. Adonis P-value is indicated by symbol: * P<0.05. ** P<0.01. *** P<0.001

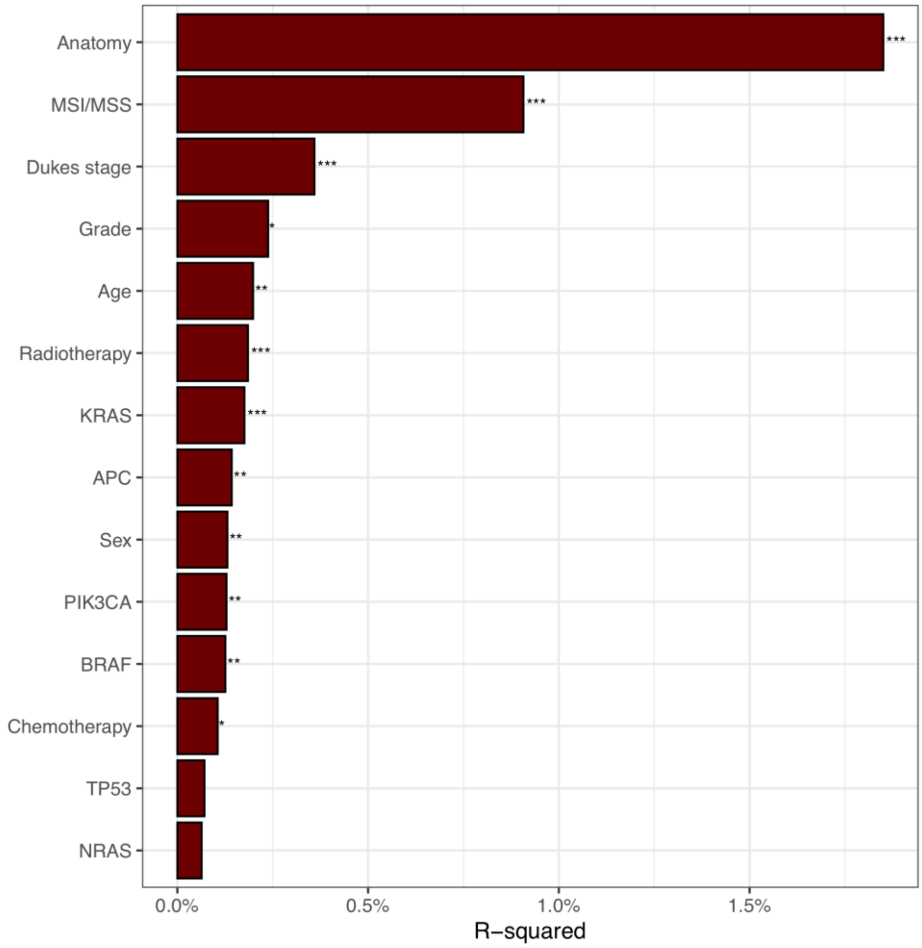

**Supplementary Fig. 33. Example taxa distributions**. Examples of taxa significantly associated by MaAsLin2 with metadata categories. (**a**) *Akkermansia* and (**b**) *Fusobacterium* are shown. Univariate Mann-Whitney *P*-values are shown, as these plots do not include distal or rectal MSI tumours, while multivariate MaAslin2 *P*-values are calculated from all samples.

**a**

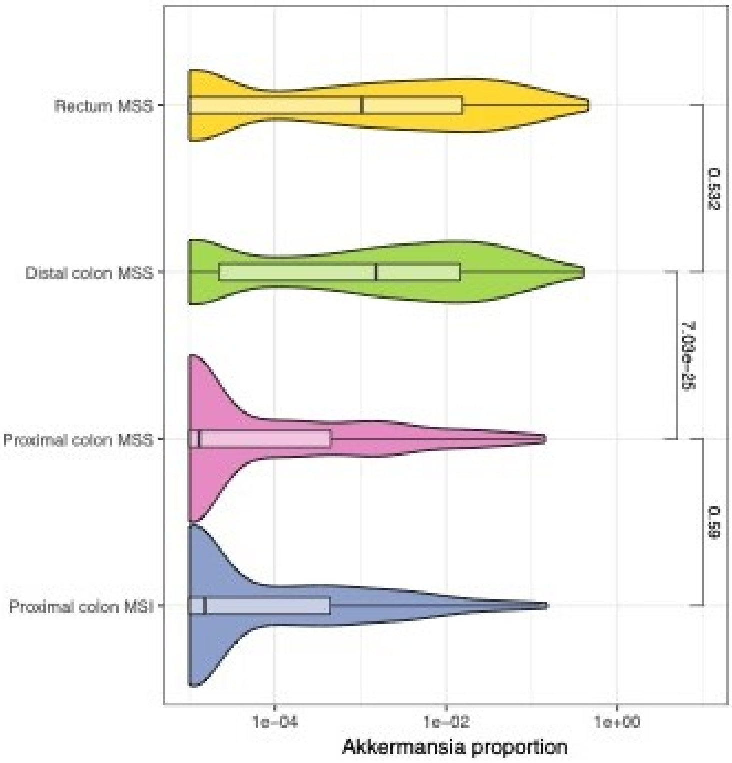


**b**

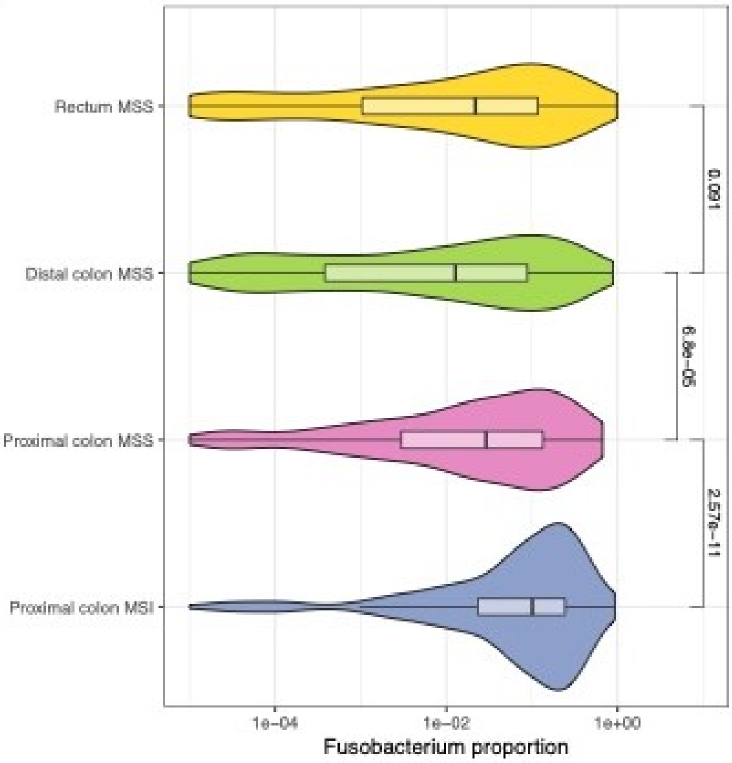

**Supplementary Fig. 34: *E.coli* distribution for PKS-positive and -negative MSS CRCs.** *E.coli* proportions in tumours with either ID18 or SBS88 contributing to 5% or more of the mutational burden, compared to tumours with no PKS contribution, are shown by anatomical location. No MSI tumours were PKS positive by these thresholds. Mann-Whitney p-values comparing PKS-positive and -negative tumours for each location are shown.

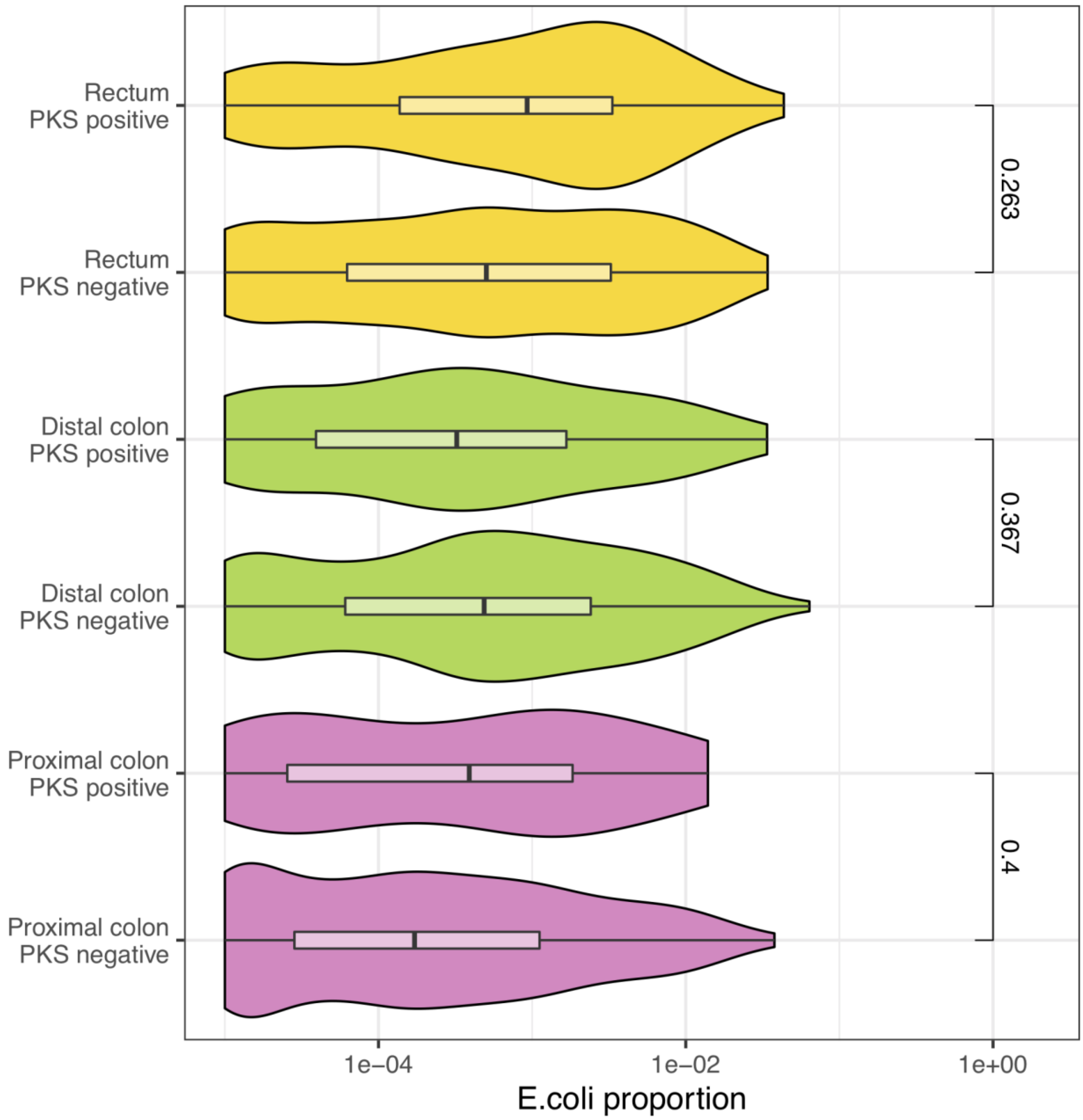

1. Giannakis, M. *et al.* Genomic Correlates of Immune-Cell Infiltrates in Colorectal Carcinoma. *Cell Rep* **15**, 857-865 (2016).
2. Grasso, C.S. *et al.* Genetic Mechanisms of Immune Evasion in Colorectal Cancer. *Cancer Discov* **8**, 730-749 (2018).
3. Network, T.C.G.A. Comprehensive molecular characterization of human colon and rectal cancer. *Nature* **487**, 330-7 (2012).
4. Seshagiri, S. *et al.* Recurrent R-spondin fusions in colon cancer. *Nature* **488**, 660-4 (2012).
5. Castro-Mondragon, J.A. *et al.* JASPAR 2022: the 9th release of the open-access database of transcription factor binding profiles. *Nucleic Acids Res* **50**, D165-d173 (2022).
6. Hammal, F., de Langen, P., Bergon, A., Lopez, F. & Ballester, B. ReMap 2022: a database of Human, Mouse, Drosophila and Arabidopsis regulatory regions from an integrative analysis of DNA-binding sequencing experiments. *Nucleic Acids Res* **50**, D316-d325 (2022).
7. Guo, Y.A. *et al.* Mutation hotspots at CTCF binding sites coupled to chromosomal instability in gastrointestinal cancers. *Nat Commun* **9**, 1520 (2018).
8. Katainen, R. *et al.* CTCF/cohesin-binding sites are frequently mutated in cancer. *Nat Genet* **47**, 818-21 (2015).
9. Arnedo-Pac, C., Muinos, F., Gonzalez-Perez, A. & lopez-Bigas, N. Signatures 1 and 17 show increased propensity to create mutational hotspots in the human genome. *In preparation.* (2022).
