## Supplementary material for "Whole genome sequencing of 2,023 colorectal cancers reveals mutational landscapes, new driver genes and immune interactions": Methods

**ONLINE METHODS**

### 1. Sample collection

Samples were obtained as part of the 100,000 Genomes Project (100KGP) cancer programme, an initiative for high throughput tumour sequencing for NHS cancer patients ^1,2^. Patient recruitment was organised by 13 Genomic Medicine Centres (GMCs) and their affiliated hospitals across England. All patients provided written informed consent. Tissue collection, sample preparation, DNA extraction and quantification were undertaken locally, followed by transfer of DNA to a central national biorepository. Whole genome sequencing of paired tumour-constitutional (whole blood-derived) DNA was performed by Illumina, and processed BAM files were then transferred to Genomics England, who performed additional processing, quality checking and data storage.

### 2. Whole genome sequencing and somatic variant calling

Sequencing, mapping and variant calling were generally performed as described ^3^, although we used a less stringent variant allele frequency to enable analysis of sub-clonal mutations.

#### 2.1. Sequencing and alignment

Samples were prepared using an Illumina TruSeq DNA PCR-free library preparation kit and sequenced on a HiSeq X, generating 150 base pair (bp) paired-end reads. Tumour and constitutional DNAs were sequenced to average depths of 100x and 33x respectively. Poor sequencing quality outliers were identified using principal component analysis and removed, based on the following quality metrics: percentage of mapped reads; percentage of chimeric DNA fragments; average insert size; AT/CG dropout; and unevenness of local coverage. Illumina’s North Star pipeline (v2.6.53.23) was used for the primary whole genome sequencing (WGS) analysis. Sequence reads were aligned to the *Homo sapiens* GRCh38Decoy assembly using Isaac (v03.16.02.19) ^4^. Overall, PCR-free tumour and germline sequencing data for 2,492 fresh-frozen colorectal cancer (CRC) samples were obtained from the 100kGP main programme version 8 release and taken into our analysis.

#### 2.2. Single nucleotide variant and indel calling

Single nucleotide variant (SNV) and small insertion and deletion (indel) calling was performed using Strelka (v2.4.7). In addition to the default Strelka filters, we applied the following exclusion filters:

- Variants with a germline allele frequency (AF) >1% in the full Genomics England dataset.
- Variants with a population germline AF >1% in the gnomAD database ^5^.
- Somatic variants with frequency >5% in the Genomics England cancer dataset. A 5% cut-off was chosen based on the frequency of recurrent non-synonymous variants in Cancer Gene Census genes ^6^.
- Variants overlapping simple repeats as defined by Tandem Repeats Finder ^7^.
- Indels in regions with high levels of sequencing noise where >10% of the base calls in a window extending 50bps either side of the indel have been filtered out by Strelka due to the poor quality.
- Indels within 10bps of 100kGP or Gnomad v3 germline indel with AF > 1%.
- Variants in regions of poor mappability where the majority of overlapping 150bp reads do not map uniquely to the variant position.
- SNVs resulting from systematic mapping and calling artefacts present in both tumour and normal 100kGP sample sets. We tested whether the ratio of tumour allele depths at each somatic SNV site was significantly different to the ratio of allele depths at this site in a panel of normal samples (PoN) using Fisher’s exact test. The PoN was composed of a cohort of 7,000 non-tumour genomes from the Genomics England dataset. At each genomic site only individuals not carrying the relevant alternative allele were included in the count of allele depths. The mpileup function in bcftools (v1.9) was used to count allele depths in the PoN. To replicate Strelka filters duplicate reads were removed and quality thresholds set at mapping quality ≥5 and base quality ≥5. All somatic SNVs with Fisher’s exact test phred score <80 were filtered, with the threshold determined by optimising precision and recall calculated from a TRACERx truth set ^8^.

#### 2.3. Removing alignment bias introduced by soft clipping of semi-aligned reads

The Isaac --clip-semialigned parameter invokes the soft clipping of read ends until five consecutive bases are matched with the reference genome. This soft clipping therefore results in the loss of support for alternate alleles occurring within five bases of each read end, leading to artefactually low VAFs. To address allelic bias introduced by this clipping, we introduced FixVAF to soft clip all reads by 5bps at each end, regardless of whether any of the bases are variant sites or whether the reads support reference or alternate alleles (**Supplementary Fig. 3**) ^9^. Reads containing small insertions and deletions at variant positions were ignored.

#### 2.4. Identifying microsatellite instability

Tumours with microsatellite instability (MSI) were identified using MSINGS ^10^ following the previously described procedure for background model generation (<https://github.com/sheenamt/msings/blob/master/Recommendations_for_custom_assays>). A set of 132 tumours with known MSI status (106 MSS, 26 MSI) was randomised into test and training sets of 53 MSS and 13 MSI cases (i,e. two sets of 66 cases). Microsatellite sites were generated using MISA ^11^. Only sites overlapping regions of good mappability were considered. Sites measured as unstable in >5 MSS test tumours and sites not unstable in ≥1 test MSI tumours were removed. The background model produced using the training set was able to perfectly distinguish between MSI and MSS samples in the test set using default MSINGs settings and was then applied to the full CRC cohort.

#### 2.5. Identifying pathogenic germline variants

Tumours with pathogenic somatic or germline variants in *POLE* or *POLD1* were identified considering the 22 known pathogenic variants reported by Rayner, et al. ^12^. In total, 18 tumours (17 MSS, 1 MSI) had a pathogenic germline (n=1) or somatic (n=17) *POLE* variant and these were considered as a separate POL group in all subsequent analyses. All of the highest mutational burden tumours were either MSI or had a known pathogenic *POLE* variant, indicating that no pathogenic polymerase proofreading domain mutations were missed. Tumours with pathogenic *POLE* variants also exhibited high SBS10a and SBS10b activity, which are established indicators of *POLE* exonuclease domain mutations ^13^.

#### 2.6. Copy number aberration calling

Somatic copy number aberrations (CNAs) were called using a framework implemented in the CleanCNA R-package. Genome-wide subclonal CNAs were first called using Battenberg v2.2.8 ^14^. To check the quality of these CNA calls we applied DPClust ^14^ and CNAqc ^15^ to the CNA profiles and SNV VAFs. DPClust clusters variants by their cancer cell fraction (CCF), whilst CNAqc compares observed and expected peaks in SNV VAF distributions to assess CNA calling accuracy. A sample was classified as ‘pass’ if it met both of the following criteria, and ‘fail’ otherwise:

1. A clonal cluster of SNVs (0.95 ≤ CCF ≤ 1.05) was identified by DPClust. This clonal cluster was required to have either the highest CCF of all SNV clusters or contain the largest number of SNVs. SNV clusters containing < 1% of all sample SNVs were removed before assessment.
2. The difference in purity estimates from Battenberg and CNAqc was < 5%. CNAqc estimates sample purity considering peaks in SNV VAF distributions in genome regions with one of five copy number states (1:0, 1:1, 2:0, 2:1, 2:2).

CNAs were profiled a maximum of four times per sample and the procedure was stopped if both criteria were met. After a failure, CNA were re-called using Battenberg with re-estimated sample purity and tumour ploidy. After the first fail purity and ploidy were re-estimated using information from DPClust, where *CCF_top_* is the CCF of the SNV cluster with the greatest CCF:

$$p_{new}=\rho_{old}CCF_{top}$$

$$\Psi_{new}= \frac{\left( \rho_{old}\psi_{old} \right)+2(\rho_{new}-\rho_{old})}{\rho_{new}}$$

After the second fail purity and ploidy were re-estimated using Ccube ^16^, and after the third and fourth fails purity and ploidy were re-estimated using CNAqc. If a sample failed after four re-runs, then it was removed from downstream analyses reliant on CNA. Passing CNA profiles were produced for 1,765/2,023 samples.

#### 2.7. Structural variant calling

Structural variants (SVs; also referred to as chromosomal rearrangements) represent two reference positions (referred to as rearrangement breakpoints) that are non-adjacent in the reference genome and juxtaposed in a specific orientation. We identified somatic rearrangements using a graph-based consensus approach comprising Delly ^17^, Lumpy ^18^ and Manta ^19^ whilst also considering support from CNAs (**Supplementary Fig. 1**). Rearrangements were first called using the three individual callers with default parameters. Delly was run with post-filtering of somatic SVs using all normal samples, as described in the Delly documentation. Rearrangements from the three individual callers were further filtered if any reads supporting the variant were identified in the matched normal, if <2% of tumour reads supported the variant, or if either variant breakpoint was in a telomeric or centromeric region or on a non-standard reference contig (*i.e.,* not chromosomes 1-22, X or Y). Remaining rearrangements were merged with a modified version of PCAWG Merge SV, which uses a graph-based approach to identify and merge rearrangements identified by multiple callers, allowing a maximum 400bp difference in breakpoint position to account for variant calling ambiguity ^20^. Rearrangements were included in the final data set if they were identified by at least two callers, or by a single caller but with a breakpoint within 3kb of a CNA segment boundary. SVs were only called in the 1,765/2,023 samples with CNA profiles passing QC criteria.

Somatically acquired long interspersed nuclear element (LINE-1) retrotransposition events were identified using xTea ^21^. Other retrotransposition categories, including Alu elements, SINE-VNTR-Alu elements and processed pseudogene, collectively comprise ≤3% of retrotransposition events across human cancers ^22^ and were therefore not considered. Retrotransposition events are mechanistically distinct from other SV-generating events ^22^ and retrotransposition events were therefore excluded from our SV analyses. SVs identified using the graph-based consensus approach were classified as part of a retrotransposition event and excluded if: (1) xTea identified a transduced region within 10kb of either SV break point in the same sample, or (2) xTea identified a transduced region within 10kb of either SV break point in ≥1% of CRC samples. A 10kb threshold was chosen as the majority of somatically acquired transductions span regions <10kb from a canonical LINE-1 element ^23^.

Putative kinase gene fusions were identified considering the following genes: *ALK*, *BRAF*, *EGFR*, *ERBB2*, *ERBB4*, *FGFR1*, *FGFR2*, *FGFR3*, *KIT*, *MET*, *NTRK1*, *NTRK2*, *NTRK3*, *RET*, and *ROS1* ^24^. Fusions were required to involve the kinase domain of the 3’ gene and have correct strand orientation.

### 3. Clinical data

Clinical data were obtained from the Genomic Medicine Centres (GMCs), NHS Digital (NHSD) and Public Health England’s National Cancer Registration and Analysis Service (PHE-NCRAS), through the Genomics England Research Environment as part the 100kGP main programme version 10 release. Survival data were obtained from the 100kGP main programme version 13 release. Tumour samples sequenced by Genomics England were matched to their respective PHE-NCRAS records using the date of tumour sampling reported by Genomics England and dates of biopsy/treatment reported by PHE-NCRAS, allowing a maximum discrepancy of seven days.

Clinical data included sex, age at tumour sampling, date of cancer diagnosis, date of last reported follow-up and date of death, tumour histology, tumour type (primary, recurrence of primary or metastases), anatomical site sampled, anatomical site of primary tumour, Dukes stage, and tumour grade (differentiation). For some variables, data were obtained from multiple sources (GMC, NHSD, PHE-NCRAS) and any conflicts between these sources were resolved by individual inspection. If Dukes staging was not available, it was inferred from TNM staging if reported. Anatomical site of primary tumour was reported at different resolutions by the different data sources (*e.g.,* one source may report site as proximal colon, whilst another may report it as caecum). To resolve and standardize site, we therefore constructed an anatomical ontology based on ICD-10-CM codes and assigned sample terms to this ontology. This allowed us to consider anatomical site at two levels of resolution: less specific (colon; proximal colon; distal colon; rectum) and more specific (caecum; ascending colon; hepatic flexure; transverse colon; splenic flexure; descending colon; sigmoid colon, rectosigmoid colon; rectum). The proximal colon comprised the caecum, ascending colon, hepatic flexure, and transverse colon, the distal colon comprised the splenic flexure, descending colon, and sigmoid colon, and the rectosigmoid junction was considered part of the rectum.

Germline mutations in the Mendelian CRC predisposition genes (*APC, MSH2, MLH1, MSH6, MUTYH, SMAD4, BMPR1A, GREM1, STK11, NTHL1, POLE* and *POLD1*) were explored in the sequenced constitutional DNA. Disease-causing changes were identified based on ClinVar annotation as *Pathogenic* or *Likely Pathogenic,* with the exception of *POLE* and *POLD1* which used the method described in section 2.5. Evidence of pathogenic bi-allelic changes was required to diagnose the recessive conditions (*MUTYH, NTHL1, MBD4*) and no such cases were found.

### 4. Sample selection

Since tumour sample purity and sequencing data quality affect the sensitivity and precision of variant calling ^25^ we excluded samples using the following quality control (QC) procedures (**Supplementary Table 2**).

- Tumour samples were excluded if cross-contamination of the tumour sample was >1%, as estimated by VerifyBamID ^26^.
- Tumour samples were excluded if cross-contamination of the matched germline sample was >1%, as estimated by VerifyBamID.
- Estimating tumour sample purity is particularly difficult when purity is low. We therefore used the distribution of single-nucleotide variant (SNV) variant allele frequencies (VAFs) to identify low purity samples, as a low average SNV VAF can be indicative of low sample purity ^27^. Tumour samples with a median SNV VAF <0.1 were excluded, with this threshold chosen based on the smaller numbers of potential driver variants observed in microsatellite stable (MSS) CRC samples when compared to all MSS CRC samples (**Supplementary Fig. 2**). Here, driver mutations were defined as any potentially pathogenic coding variant called in 63 driver genes previously identified in MSS CRC ^28-31^.
- Tumour samples were excluded if <100 SNVs were called, as this number is below the smallest number of SNVs previously reported in CRC whole genomes ^20^ and therefore suggestive of low sample purity or sequencing data quality.
- Tumour samples were excluded if many mutations were associated with a likely artefactual mutational signature ^13^.

In total 286/2492 (11.5%) of tumour samples were excluded based on the above criteria.

Tumour samples were also excluded if essential clinical data was missing, or there were unresolvable conflicts between the sources from which clinical data were obtained (GMCs, NHSD, PHE-NCRAS) (**Supplementary Table 2**). In total 183/2206 (8.3%) of tumour samples that passed tumour sample purity and sequencing data QC were excluded based on clinical data, using the following criteria:

- GMC, NHSD and PHE-NCRAS reported conflicting years of birth.
- Sex reported by GMC, NHSD and/or PHE-NCRAS did not match sex inferred from sequencing data.
- GMC, NHSD and PHE-NCRAS did not report tumour histology or reported conflicting histology.
- Tumour was not classified as a colorectal adenocarcinoma.
- Missing or conflicting data meant it was unclear whether the primary tumour or a metastasis was sampled.
- Multiple primary tumours or multiple metastases from a single individual were sequenced. In these instances, the primary tumour sample or metastases sample with the highest purity was included, and all other primary tumour samples or metastases samples were excluded. This procedure was completed after all other exclusion criteria had been applied. Primary tumours and metastases were considered separately.

Based on these criteria, 2,023 CRC adenocarcinoma samples were suitable for analysis (**Supplementary Table 2**). This cohort comprises 1,898 primary tumours, 122 metastases and 3 recurrences of primary tumours from 2,017 patients. Six patients had both a primary tumour and a metastasis sample sequenced and included. Clinical data completeness is detailed in **Supplementary Table 20**.

### 5. Single nucleotide variant and indel drivers

#### 5.1. Mutation annotation

Somatic mutations were annotated to Ensembl (v101, GRCh38) using Variant Effect Predictor (VEP) ^32^. The following parameters were used: *vep -i <input_vcf> --assembly GRCh38 –no_stats –cache –offline –symbol –protein -o <output> --vcf –canonical –dir <ref_dir> --hgvs –hgvsg –fasta <GRCh38_fasta> --plugin CADD,<CADD_score_file> --plugin UTRannotator,<GRCh38_uORF_reference>*

The CADD score file was obtained using CADD (v1.6) ^33-35^ with scores attained for all SNV and indel mutations using the CADD software (https://github.com/kircherlab/CADD-scripts) before being utilised by the VEP CADD plugin.

#### 5.2. Protein-coding driver identification

Protein-coding driver genes were identified using the IntOGen pipeline (downloaded February 2021) ^36^.

##### 5.2.1. Pre-processing of input mutations

Somatic mutations passing the filtering criteria described previously were subject to initial sample and mutation pre-processing. In the case of multiple tumours from the same patient, the primary tumour was used. Within each cohort (*i.e.* Primary MSS, Primary MSI, Primary POL, Metastases MSS) tumours were flagged for exclusion from downstream driver gene identification if they contained >10,000 mutations and had an outlier mutation count, defined as *upper quartile + (1.5 × interquartile range)*. Mutations present in a Hartwig Consortium PoN were also excluded ^37^. Unless otherwise specified, mutations were mapped to canonical protein-coding transcripts from Ensembl (v101, GRCh38).

##### 5.2.2. Driver identification methods

Seven driver gene identification methods were run through the IntOGen pipeline:

1. dNdSCV ^38^ is designed to detect genes under positive selection that show an excess of nonsynonymous (missense, nonsense, essential splice) mutations after correction for local trinucleotide context. In the primary POL cohort the parameter “*max_coding_muts_per_sample = Inf*” was used, due to the high proportion of hypermutated tumours.
2. OncodriveFML ^39^ aims to detect driver genes that show an enrichment of mutations with high functional impact. CADD scores were used as measure of functional impact ^33-35^.
3. OncodriveCLUSTL ^40^ is a method designed to detect driver genes that are enriched for linear mutation clusters. In the primary POL cohort, pentamer signatures were used rather than trinucleotide signatures, due to the improved performance of the pentanucleotide-based background models compared to that of trinucleotides in these tumours.
4. cBaSE ^41^ aims to detect driver genes under positive selection that exhibit a significant mutation count bias after correction by tri-nucleotide context.
5. MutPanning ^42^ is designed to detect driver genes that exhibit enrichment of mutations with unusual nucleotide contexts compared to a background model.
6. HotMaps3D ^43^ detects driver genes containing missense mutations that are spatially clustered together in the 3-dimensional structure of the protein. Protein structures were downloaded from The Protein Data Bank (PDB) in March 2020 ^44^.
7. smRegions ^45^ detects genes containing an enrichment of nonsynonymous mutations in regions of interest, such as protein domains, after correcting for trinucleotide context. This analysis utilised information from protein family (Pfam) domains which were mapped to Ensembl (v101) canonical transcripts.

##### 5.2.3. Combination of driver identification methods

The results of the 7 driver identification methods were combined similarly as described ^46^. Briefly, the driver combination procedure considered the top-100 ranked genes and their associated *P* and *Q-*values in each of the seven driver identification methods. Somatically mutated genes assigned as Tier 1 or Tier 2 in the COSMIC Cancer Gene Census (CGC) ^47^ were designated as the “truth” set of known drivers. Through comparison of the relative enrichment of CGC genes in the top ranked gene lists, a per-method weighting was obtained. Per-method ranked lists were combined using Schulze’s voting method to generate a consensus ranking, with combined *P*-values estimated using a weighted Stouffer Z-score method.

Driver candidates were then classified into the following tiers:

- Tier 1. Candidates where the consensus ranking was higher than the ranking of the first gene with Stouffer *Q*>0.05. These represent high confidence drivers.
- Tier 2. Candidates not meeting the criteria for Tier 1, but which are CGC genes and show a combined Stouffer *Q_CGC_<*0.25. These represent a set of “rescued” known cancer drivers.
- Tier 3. Candidates not meeting the criteria for Tier 1 or Tier 2 but with Stouffer *Q*<0.05. These represent lower confidence drivers.
- Tier 4. Candidates not meeting criteria for Tier 1 or Tier 2 and with Stouffer *Q*>0.05. These represent candidates that are not likely to be drivers.

##### 5.2.4. Post-processing of candidate drivers

Candidate driver genes were filtered based on the following annotations:

1) AUTOMATIC FAIL. A candidate driver gene would be excluded from further consideration if annotated with at least one of the following:
   a) *TIER4*. Categorised as Tier 4 by the combination procedure.
   b) *1_METHOD*. Only significant (Q<0.1) in one of the seven methods (non-CGC genes).
   c) *EXPRESSION*. Gene has very low or no expression in relevant TCGA tumour type.
   d) *OLFACTORY_RECEPTOR*. Gene is in list of olfactory receptor genes.
   e) *KNOWN_ARTIFACT*. Gene is in a list of known artifacts or long genes (e.g., TTN).
2) *MANUAL REVIEW*. If a gene is not excluded based on any *AUTOMATIC FAIL* filters, it is retained as a candidate driver:
   a) *GERMLINE*. Non-Tier 1-CGC gene has ≥1 mutations per sample and oe_syn/ms/lof > 1.5 based on GnomAD (v2.1) constraint metric estimates.
   b) *SAMPLE_3_MUTS*. Non-CGC gene where there are ≥3 mutations in ≥1 tumour.
   c) *LITERATURE*. Non-CGC gene where there are no literature annotations according to CancerMine ^48^.
3) *AUTOMATIC PASS*. Is not flagged by any *AUTOMATIC FAIL* or *MANUAL REVIEW* filters.

Candidate driver roles were assigned on the basis of dN/dS ratios for missense (*wmis*) and nonsense (*wnon*) mutations for the given gene derived from dNdSCV (<https://bitbucket.org/intogen/intogen-plus/src/master/core/intogen_core/postprocess/drivers/role.py>):

- A “distance” metric was calculated by $distance=\frac{\left( wmis-wnon \right)}{\sqrt{2}}$
- Candidate drivers with distance >0.1 represent those with an excess of missense to nonsense mutations and are therefore considered *oncogenes*.
- Candidate drivers with distance <0.1 represent those with an excess of nonsense to missense mutations and are therefore considered *tumour suppressor genes* (*TSGs*).
- Otherwise, the role of the candidate driver is unclear and considered *ambiguous*.

In the case of multiple cohorts being run representing subsets of a given tumour type, a consensus role was designated comparing between each subtype role:

- Oncogene if ≥1 cohort and no other cohorts assigned as TSG.
- TSG if in ≥1 cohort and no other cohorts assigned as oncogene.
- Ambiguous otherwise.

Gene candidates were annotated by their overlap with any IntOGen cohorts from a previous IntOGen pan-cancer analysis (1 February 2020) as well as from a pan-cancer TCGA analysis ^49^.

Following post-processing, nine candidate genes were excluded as likely false-positives: *ANK1* (not expressed in CRC), *ANO2* (olfactory calcium channel), *CACNA1E* (driven by strand bias artefact mutation p.Ile95Leu), *CHD2* (detected mutations inconsistent with known role), *CSMD3* (not expressed in CRC), *FSIP2* (not expressed in CRC), *KRT12* (not expressed in CRC), *LILRB3* (not expressed in CRC), and *MYH11* (myosin heavy chain protein).

#### 5.3. Non-coding driver identification

##### 5.3.1. Defining sets of non-coding regions

Regions from candidate non-coding elements overlapping coding sequence (CDS) or exon regions from canonical protein-coding transcripts were removed using bedops (v2.4.39) ^50^.

The following sets of non-coding regions were defined:

1. Core promoters (n=19,283). Defined based on transcription start site (TSS) of canonical protein-coding transcripts: 200bp < TSS < 50bp. CDS regions were removed.
2. Distal promoters (n=19,296). Defined based on TSS of canonical protein-coding transcripts: 2kb < TSS. CDS regions were removed.
3. Five-prime untranslated regions (5’-UTRs; n=18,613). Defined based on canonical protein-coding transcripts. CDS regions were removed.
4. Three-prime untranslated regions (3’-UTRs; n=18,806). Defined based on canonical protein-coding transcripts. CDS regions were removed.
5. LincRNAs (n=16,510). Based on exon regions from transcripts annotated as lincRNAs in Ensembl (v101). Exon regions from canonical protein-coding transcripts were removed.
6. miRNAs (n=1,793). Based on regions from transcripts annotated as miRNAs in Ensembl (v101). Exon regions from canonical protein-coding transcripts were removed.
7. Non-canonical splice regions (n=18,163). Defined from regions extending 30bp into the intron from essential splice donor or acceptor sites in canonical protein-coding transcripts. Exon regions from canonical protein-coding transcripts were removed.
8. Enhancers (n=130,996). Defined from Ensembl v101 regulatory elements annotated as “Enhancer”. Exon regions from canonical protein-coding transcripts were removed.
9. Open chromatin regions (n=95,344). Defined from Ensembl (v101) regulatory elements annotated as “Open chromatin”. Exon regions from canonical protein-coding transcripts were removed.
10. CTCF sites (n=173,711). Defined from Ensembl (v101) regulatory elements annotated as “CTCF sites”. Exon regions from canonical protein-coding transcripts were removed.
11. TF binding sites (n=29,259). Defined from Ensembl (v101) regulatory elements annotated as “TF binding sites”. Exon regions from canonical protein-coding transcripts were removed.

##### 5.3.2. Detecting non-coding drivers

Potential non-coding driver mutations were identified in non-hypermutated primary MSS tumours (n=1,442). OncodriveFML was run on sets of non-coding regions according to the following amended parameters from the protein-coding analysis: “indel-max” indels are treated as a set of substitutions, with the functional impact of the indel mutation being the maximum of all the substitutions, and the background simulated as substitutions. A *Q*<0.01 threshold was considered as significant.

#### 5.4. SNV mutations exhibiting extreme strand bias

SNV mutations that otherwise pass filtering criteria as previously detailed were further scrutinised by whether they exhibited excessive strand bias (Strelka INFO field SNVSB>10). This highlighted many missense mutations causing a recurrent missense change in *CACNA1E* (p.Ile95Leu) that exhibited excessive strand bias and therefore were likely to be false-positive mutation calls. It is therefore likely that *CACNA1E* itself is a false-positive driver candidate.

#### 5.5. Driver mutation annotation

Nonsynonymous mutations in the 682 gene transcripts considered by OncoKB (v3.3) were annotated using the OncoKB API ^51^. In the first instance, the HGVSg identifier was used; in the rare instances that this failed, a combination of gene symbol, consequence and HGVSp were used to map mutations to OncoKB annotations.

##### 5.5.1. Annotation of oncogenic mutations

Nonsynonymous mutations in candidate driver genes were annotated as pathogenic if any of the following criteria were met:

1. The mutation is annotated by OncoKB as “Oncogenic”, “Likely Oncogenic” or “Predicted Oncogenic”.
2. The driver is classified as an oncogene, the mutation consequence is missense, and the mutation is recurrent (seen in ∈3 tumours in cohort).
3. The driver is classified as a TSG or ambiguous and either:
   a. Consequence is protein-truncating (splice acceptor, splice donor, frameshift, stop lost, stop gained or start lost).
   b. Consequence is missense and mutation is recurrent (seen in ∈3 tumours in cohort).

For *POLE*, oncogenic annotations were restricted to missense mutations in the exonuclease domain (amino acid residues 268-471).

Nonsynonymous mutations not meeting these criteria were considered as variants of uncertain significance (VUS).

##### 5.5.2. Lollipop plots of driver gene mutations

Lollipop plots of driver gene mutations were generated using the trackViewer R-package ^52^. Pfam protein domains mapping to the Ensembl (v101) canonical transcripts were plotted. The protein position was taken from the first position in the HGVSp annotation, apart from splice donor and acceptor mutations where the codon nearest to the HGVSc transcript position was assigned as the protein position.

##### 5.5.3. Timing driver mutations

The relative evolutionary timings of candidate driver mutations were obtained using MutationTimeR (<https://github.com/gerstung-lab/MutationTimeR>) ^53^. Copy number input for MutationTimeR was prepared from Battenberg segmentation files, with the clonal frequency of each segment taken as the tumour purity. In the case of subclonal calls, the clonal frequency was calculated by multiplying the tumour purity by the clonal fraction. The clusters input for MutationTimeR was prepared from DPClust cluster estimates. The VAF proportion was calculated by multiplying the estimated cluster CCF by the tumour purity. Superclonal clusters (CCF>1.1) were removed. VCF input for MutationTimeR was obtained from the small somatic SNV/indel variant VCFs which had been filtered as previously described. For SNVs, alt and ref depths were obtained using FixVAF (<https://github.com/danchubb/FixVAF>). For indels, ref and alt depths were obtained from Tier2 Strelka TAR and TIR fields respectively. Only mutations within Battenberg copy-number segments were retained (note: for male XY tumours with only 1 copy of the X chromosome copy number information is restricted to the pseudoautosomal region (PAR) and Battenberg was not run on the Y chromosome).

MutationTimeR was run with 1,000 bootstraps. For tumours previously defined as having undergone whole genome doubling (WGD), the parameter “isWgd” was set to TRUE. Mutations were then classified into estimated simple clonal states (as per Fig.1a of^53^):

- “Clonal [early]” – Mutation on ≥ 2 copies per cell
- “Clonal [late]” – Mutation on 1 copy per cell, no retained allele
- “Clonal [NA]” – Mutation on 1 copy per cell, either on amplified or retained allele
- “Subclonal” – Mutation on < 1 copy per cell

#### 5.6. Mutational signature attribution

SeqInfo VCFs produced as part of SigProfilerMatrixGenerator ^54^ were used to map somatic mutations from input VCFs to their SBS96, DBS78 or ID83 contexts and then to the final SigProfilerExtractor COSMIC (v3.2) decomposed signature probabilities.

#### 5.7. Annotation of double base substitution mutations

Per-tumour VCFs containing DBS mutations, either directly called originally by Strelka, or originally called by Strelka as two adjacent SNVs and reconstructed as DBS mutations, were created and mutation consequences re-calculated using VEP as above.

### 6. Patterns of somatic copy number alteration

#### 6.1. Whole genome duplication classification

Tumours were classified as whole genome duplicated (WGD) considering the average genome copy number state ($\psi_{ave}$):

$$\psi_{ave}=\frac{\sum_{i=1}^{S} L_{i}(C_{i}^{Maj}+C_{i}^{Min})}{\sum_{i=1}^{S} L_{i}}$$

where, $S$ is the number of copy number genome segments, $C_{i}^{Maj}$ and $C_{i}^{Min}$ are the major and minor allele copy numbers for genome segment $i$, and $L_{i}$ is the base pair length of genome segment $i$. If there is evidence of sub clonal alteration, then the copy number states corresponding to the largest tumour cell fraction are considered. Tumours are classified as WGD as follows ^55^:

$$\left\{ \begin{aligned} WGD, &if 2.9-2H< \psi_{ave} \\ Not WGD, &otherwise \end{aligned} \right.$$

Where, $H$ is the fraction of the genome with a minor allele copy number of 0.

#### 6.2. Classification of copy number alterations

Individual copy number alternations were grouped into six categories: homozygous deletion (HD), loss of heterozygosity (LOH) inckuding copy-neutral LOH, other loss (OLOSS), no change (NOC), gain (GAIN) and amplification (AMP). Classification considers whether a tumour has undergone WGD.

Where subclonal CNAs existed, the copy number state corresponding to the largest cell fraction was used. Classification into one of the six categories overlaps significantly between non-WGD and WGD tumours, with differences relating to total copy number. Differences include:

- In non-WGD tumours, segments are classified as LOH if one allele has a copy number state of 0 and the total copy number (t_CN_) ≤2. In WGD tumours, segments are classified as LOH if one allele has a copy number state of 0 and t_CN_ ≤4.
- Non-WGD tumours do not have an OLOS category.
- NOC is defined as 1+1 in non-WGD tumours and 2+2 in WGD tumours.
- In non-WGD tumours, segments are classified as GAIN if 2< t_CN_ ≤5. In WGD tumours, segments are classified as GAIN if 4< t_CN_≤10.
- In non-WGD tumours, segments are classified as AMP if t_CN_ >5. In WGD tumours, segments are classified as AMP if t_CN_ >10.

|  |  | **HD** | | **LOH** | | **OLOS** | | **NOC** | | **GAIN** | | **AMP** | |
| --- | --- | --- | --- | --- | --- | --- | --- | --- | --- | --- | --- | --- | --- |
| **Non-WGD** | **Copy number states** | 0 | 0 | 1 | 0 |  | | 1 | 1 | 2 | ≥1 | 4 | ≥2 |
|  |  |  | | 2 | 0 |  | |  | | 3 | ≥0 | 5 | ≥1 |
|  |  |  | |  | |  | |  | |  | | ≥6 | ≥0 |
|  | **Overall conditions** | t_CN_ = 0 | | 0< t_CN_ ≤ 2  1 allele = 0 | |  | | t_CN_ = 2 | | 2< t_CN_ ≤5 | | 5< t_CN_ | |
| **WGD** | **Copy number states** | 0 | 0 | 1 | 0 | 1 | 1 | 2 | 2 | 3 | ≥2 | 6 | ≥5 |
|  |  |  | | 2 | 0 | 2 | 1 |  | | 4 | ≥1 | 7 | ≥4 |
|  |  |  | | 3 | 0 | 3 | 1 |  | | 5 | ≥0 | 8 | ≥3 |
|  |  |  | | 4 | 0 |  | |  | |  | | 9 | ≥2 |
|  |  |  | |  | |  | |  | |  | | 10 | ≥1 |
|  |  |  | |  | |  | |  | |  | | ≥11 | ≥0 |
|  | **Overall conditions** | t_CN_ = 0 | | 0< t_CN_ ≤4  1 allele = 0 | | 1< t_CN_ ≤4  1 allele == 1 | | t_CN_ = 4 | | 4< t_CN_ ≤10 | | 10< t_CN_ | |

#### 6.3. Positional enrichment of copy number alterations

##### 6.3.1. Preparing GISTIC input

Recurrent arm-level copy number events, as well as focal amplifications and deletions, were identified using GISTIC (v2.0.2.3) ^56^. For all samples with CNA profiles passing QC criteria, a copy number segmentation file suitable for GISTIC input was generated using Battenberg output. Chromosomal coordinates and major (nMaj) and minor (nMin) copy number states were obtained for each copy number segment identified by Battenberg. In the case of subclonal copy number segments, nMaj and nMin values corresponding to the large tumour cell fraction were considered.

Per-segment normalised copy numbers ($SegCN$) were calculated differently for tumours with WGD (where ploidy was assumed to be 4) and without WGD (where ploidy was assumed to be 2). $SegCN$ was thresholded to a minimum of -2 and maximum of 2.

For non-WGD tumours, per-segment normalised copy number was calculated as:

$$SegCN=(nMaj+nMin)-2$$

For non-WGD tumours from males, X chromosome per-segment normalised copy number was calculated as:

$$SegCN=(nMaj+nMin)-1$$

For WGD tumours, per-segment normalised copy number was calculated as:

$$SegCN=\frac{(nMaj+nMin)-4}{2}$$

For WGD tumours from males, X chromosome per-segment normalised copy number was calculated as:

$$SegCN=(nMaj+nMin)-2$$

##### 6.3.2. Running GISTIC

GISTIC was run using the following parameters: *-conf 0.99 -broad 1 -qvt 0.25 -genegistic 1 -gcm extreme -brlen 0.5 -rx 0 -twoside 1 -scent median -armpeel 1 -arb 1 -refgene hg38.UCSC.add_miR.160920.refgene.mat*

##### 6.3.3. Prioritising likely gene targets of focal amplifications and deletions

Candidate target genes at focal amplifications and deletions were annotated using the following criteria:

1. Overlap with genes at focal amplifications and deletions reported in a previous pan-cancer study that used GISTIC ^57^. Comparisons were made both with the overall pan-cancer GISTIC analysis, as well as GISTIC analysis restricted to the given tumour type. Special consideration was given to genes specifically highlighted by Zack, et al. ^57^ as being candidates.
2. Overlap with Cosmic Cancer Gene Census genes and whether their annotated role (oncogene [OG], tumour suppressor gene [TSG] or ambiguous) is consistent with the copy number change (OG with amplifications and TSG with deletions) ^24^.
3. Overlap with driver genes identified in this study and whether their likely role (OG, TSG or ambiguous) is consistent with the copy number change (OG with amplifications and TSG with deletions).

Based on the above criteria, consensus driver genes were manually assigned to peaks. Comparisons were made with all potential gene synonyms as available from the HUGO gene nomenclature name committee (<https://www.genenames.org/>).

##### 6.3.4. Defining copy number segments overlapping recurrent CNAs

Alterations from the broad analysis with *Q*<0.05 were taken to indicate recurrent arm-level events. Copy number segments comprising greater than half of the total chromosome arm size were taken to indicate arm-level events.

In the case of focal events identified by GISTIC, the “wide region” was used to compare potential extent of overlap with copy number segments. Segments were defined as overlapping focal events if either the segment interval comprised greater than half of the focal region, or vice versa, using pybedtools and bedtools (v2.3.0) ^58,59^.

Tumours were considered to have specific arm-level or focal deletions if an overlapping copy number segment was annotated as HD or LOH (as described above). Similarly, tumours were considered to have specific arm-level or focal amplifications if an overlapping copy number segment was annotated as GAIN or AMP. In the case of subclonal CNAs, nMaj and nMin values corresponding to the largest cell fractions were considered.

#### 6.4. Extrachromosomal DNA detection

Potential extrachromosomal DNA (ecDNA) molecules were detected from tumour bam files using AmpliconArchitect v1.2 (<https://github.com/virajbdeshpande/AmpliconArchitect>) ^60^. Briefly, per-tumour “seed” regions were prepared from Battenberg copy-number segmentation output if a segment was >100kb and the total copy number was > 5. AmpliconArchitect was then run using these “seed” regions to extract overlapping sequence reads from the tumour bamfile and construct candidate amplicons.

Candidate amplicons were classified using AmpliconClassifier v0.4.6 (<https://github.com/jluebeck/AmpliconClassifier>) into the following categories: 1) Cyclic (truly circularised ecDNA) 2) Complex non-cyclic 3) Linear amplification 4) No amp/invalid. Amplicons were highlighted if containing a known highly amplified oncogene (*MDM2*, *MYC*, *EGFR*, *CDK4, ERBB2, SOX2, TERT, CCND1, E2F3, CCNE1, CDK6, MDM4, NEDD9, MCL1, AKT3, BCL2L1, ZNF217, KRAS, PDGFRA, AKT1, MYCL, NKX2-1, IGF1R, PAX8*; as per ^61^).

#### 6.5. Estimation of telomere content

Telomere content was estimated from tumour and germline bam files by the methods Telomere Hunter v1.1.0 (<https://pypi.org/project/telomerehunter/>) ^62^ and Telomerecat v3.3.0 (<https://github.com/cancerit/telomerecat>) ^63^ using default parameters.

Telomere content was normalised by: $\log2\left( \frac{Tumour content}{Normal content} \right)$.

### 7. Patterns of somatic structural variation

#### 7.1. Classification of simple and complex structural variants

Rearrangements identified by the graph-based consensus approach were grouped into footprints and clusters based on their proximity within the genome, the overall number of events in the genome, and the size of these events, using ClusterSV ^20^. Rearrangement footprints represent sets of rearrangement breakpoints that are positionally associated, whilst rearrangement clusters represent sets of rearrangements that are mechanistically associated. Rearrangement footprints were described using the string approach proposed by Li, et al. ^20^. Rearrangement clusters were classified as being a simple event (deletion, tandem duplication, balanced inversion, balanced translocation, unbalanced translocation, or simple unclassified) or a complex event (chromoplexy, chromothripsis, or complex unclassified). Simple and complex events were defined as clusters comprising ≤2 or ≥3 individual rearrangements respectively.

Chromothripsis events were inferred using established criteria ^64,65^. A rearrangement cluster was defined as chromothripsis if it met all the following criteria:

- A contiguous series of four genome segments oscillating between two copy number states, or five genome segments oscillating between three copy number states.
- At least six interleaved intra-chromosomal rearrangements, as per Cortés-Ciriano, et al. ^64^.
- No evidence (false discovery rate > 0.2) that the distribution of intra-chromosomal fragment join orientations diverge from a multinomial distribution with equal probabilities for each of the four orientation categories (duplication-like, deletion-like, head-to-head inversion, and tail-to-tail inversion).

A rearrangement cluster was defined as chromoplexy if it met all the following criteria:

- Contains a chain of rearrangements spanning at least three chromosomes ^20^. SV chains were identified using a graph-based approach, in which nodes represent breakpoints, and are connected by an edge if they are not involved in the same rearrangement and fall within 1Mb of each other. Graph-based approach implemented using the igraph R-package ^66^.
- At least 50% of rearrangement footprints in the cluster represent balanced translocations, either with no observed copy number change, or a deletion bridge between the break ends.
- Consists of between 3 and 30 rearrangements.

#### 7.2. Identification of simple structural variation hotspots

Rates of somatic structural variation differ throughout the genome and are influenced by local genomic features ^20^. Genome regions enriched for simple SVs were therefore identified using a permutation-based approach considering genomic features associated with structural variation occurrence, as per Glodzik, et al. ^67^. Deletions, tandem duplications, balanced inversions, balanced inter-chromosomal translocations, and unclassified simple SVs were considered separately. Individual rearrangements that form parts of complex SVs were excluded from this analysis. Primary MSS and MSI tumours were also analysed separately, whilst primary POL tumours and metastases were not considered due to low sample numbers.

##### 7.2.1. Evaluating relationships between genomic features and SV rates

Negative binomial regression was used to test associations between genomic features and numbers of SVs of each simple class ^67^. The following features were included in the models: average total copy number across the bin in the CRC sample set, GC content, the presence of genes highly or lowly expressed in CRC, ALU repeats, other genomic repeats, segmental duplications, fragile sites, replication timing, and DNase, H3K36me3 and H3K9me3 peaks^67^. Highly and lowly expressed genes were defined as those with mean RSEM value in the top 25% and bottom 75% of protein-coding genes in TCGA samples with RNA-Seq ^31^. ALU and other genomic repeats were obtained from the UCSC Genome Browser ^68^. Segmental duplications were obtained for GRCh38 from the Segmental Duplication Database ^69^. Fragile sites were obtained from Bignell, et al. ^19^. Replication timing data from CRC epithelial cells (HCT116) were obtained from ReplicationDomain ^70^. DNAse-seq data (ENCFF443KCU) and ChIP-seq data for histones H3K36me3 (ENCFF553QXG) and H3K9me3 (ENCFF482DLD) were obtained for the large intestine from ENCODE ^71^.

##### 7.2.2. Permuting SVs

SVs were simulated to test whether the number of SVs observed in a region was greater than expected by chance given the local genomic features ^72^. SVs were simulated for each simple SV class, preserving the number and length (distance between intra-chromosomal SV break ends) of SVs observed in the CRC sample sets. To simulate SVs, the genome was divided into non-overlapping 1Mb bins and the genomic features (listed above) of each bin summarized. All genomic features were normalized to a mean of 0 and standard deviation of 1 to aid comparisons. The number of break ends expected in each bin was then estimated using the effect estimates from the previously generated negative binomial regression model. For each observed SV, an SV was simulated by sampling a bin under probabilities proportional to the expected numbers of break ends in each bin. For intra-chromosomal SVs, a partner break end was then simulated by selecting the position either upstream or downstream (with equal probability) equal in distance to the distance between the two break ends in the observed SV. For inter-chromosomal SVs, a partner break end was simulated by sampling a bin under probabilities proportional to the expected numbers of break ends in each bin, excluding bins on the same chromosome. SVs were re-simulated if either break end fell within an uncallable region (a telomere or centromere). SVs were simulated 1,000 times to generate a null distribution of expected SV numbers for the 1Mb bins.

##### 7.2.3. Identifying SV hotspots

Piece-wise constant fitting (PCF) was used to identify regions of the genome containing greater numbers of SV break ends than expected ^72^. SV break ends were first sorted by position and the distance between successive break ends calculated. PCF was then applied to the log_10_ of these inter-mutational distances (IMD). SV hotspots were identified by first computing the observed ($d_{i}^{obs}$) and expected ($d_{i}^{exp}$) number of breakends per base pair for each PCF segment (*i*):

$$d_{i}^{obs}=\frac{a_{i}}{s_{i}}$$

$$d_{i}^{exp}=\frac{\sum_{j=1}^{n} b_{j}}{ns^{bin}}$$

Where *a_i_* is the number of break ends in the segment, *s_i_* is the length of the segment in base pairs, *n* is the number of bins overlapping the segment, *b_j_* is the expected number of SVs in bin *j*, and *s^bin^* is the bin size (1Mb). A simple SV enrichment factor ($\beta_{i}^{simple}$) is then computed for each PCF segment as:

$$\beta_{i}^{simple}=\frac{d_{i}^{obs}}{d_{i}^{exp}}$$

The PCF algorithm requires parameters *γ* (that controls the smoothness of the segmentation) and *k_min_* (the minimum number of mutations in a segment). False discovery rates (FDRs) at each *β^simple^* value were estimated by applying PCF to both the observed and simulated SV sets and dividing the mean number of segments with a *β^simple^* value at least as great in the simulated SV sets by the number of segments with a *β^simple^* value at least as great in the observed SV set. A maximum FDR of one was set and FDR values equal to zero were changed to the lowest non-zero FDR value observed. Optimal *γ* and *k_min_* values were chosen by repeating this process for values of *γ* between 1 and 20, and values of *k_min_* between 2 and 20, and selecting values that maximized the number of hotspots identified, whilst minimizing the FDR. In the final analysis *γ*=10 was used throughout, whilst *k_min_*=2 was used for translocations in primary MSS samples, *k_min_*=4 was used for unclassified simple variants in primary MSI samples, and *k_min_*=10 was used otherwise. SV hotspots where no SVs were supported by CNAs were considered potential artefacts and removed. Overlapping SV hotspots identified in the same sample sets were collapsed.

##### 7.2.4. Classification of SV hotspots as fragile sites

SV hotspots were classified as fragile sites if they satisfied at least three of the following six criteria (this threshold was chosen by assessing the co-occurrence of these criteria):

- Were late replicating ^73^. Replication timing data from CRC epithelial cells (HCT116) were obtained from ReplicationDomain. Late replicating regions were defined as those with mean Repli-Seq values ≤0.
- Had low gene-density ^74^. A threshold of five genes per Mb was used.
- Overlapped a gene greater than 300kb in size. This threshold was chosen as fragile sites generally occur in chromosome regions containing genes at least 300kb in size ^75^.
- The overlapping gene of greatest size was the focus of the SV enrichment. This was assessed by computing the ratio between SV break point densities in the overlapping gene of greatest size and intergenic regions flanking 1Mb upstream and downstream. A threshold of five was used.
- Overlapped a fragile site reported by Bignell, et al. ^19^. These fragile sites were originally obtained from either the NCBI or literature curation and were mapped from NCRI36 to GRCh38 co-ordinates using LiftOver ^68^.
- Overlapped a fragile site from identified in a pan-cancer analysis of whole-genome-sequenced tumours ^76^ and mapped from GRCh37 to GRCh38 co-ordinates using LiftOver ^68^.

SV hotspots were not considered as potential fragile sites if they contained an identified CRC driver gene. SV hotspots at potential fragile sites likely occur for mechanistic rather than selective reasons and were therefore not considered further ^19^.

##### 7.2.5. Identification of candidate gene targets of recurrent SVs

Genes were reported as candidate targets of recurrent SVs if they had been identified as targets in previously analyses ^29-31,76^ or were known CRC driver genes overlapping an SV hotspot. Numbers of samples with a focal change at a candidate gene were computed considering SVs and CNAs ≤3Mb in size ^77^ at least partially overlapping the gene coding sequence.

#### 7.3. Enrichment of complex structural variation enrichment

Genome regions enriched for complex SVs were identified using a permutation-based approach, considering chromothripsis, chromoplexy and unclassified complex SVs separately. MSS and MSI tumours were also analysed separately, whilst POL tumours and were not considered due to low sample numbers. The genome was first split into non-overlapping 1Mb bins and the observed number of tumour samples with complex SV footprints ($g_{i}^{obs}$) overlapping each bin (*j)* counted. Complex SV footprint positions were next permuted 100,000 times by randomly sampling genome regions equal in size to the footprints. The expected number of tumour samples with complex SV footprints ($g_{i}^{exp}$) overlapping each 100kb bin was then estimated as the mean number of tumour samples with SV footprints overlapping the bin across all permutations. A complex SV enrichment factor ($\beta_{j}^{complex}$) was calculated for bin (*j*) as:

$$\beta_{j}^{complex}=\frac{g_{j}^{obs}}{g_{j}^{exp}}$$

FDRs at each *β^complex^* value were estimated by computing *β^complex^* for each bin in both the observed and permuted SV sets and dividing the mean number of bins with a *β^complex^* value at least as great in the permuted SV sets by the number of bins with a *β^complex^* value at least as great in the observed SV set. A maximum FDR of 1 was set and FDR values equal to zero were changed to the lowest non-zero FDR value observed.

### 8. Mutational processes

#### 8.1. Characterising single-base-substitution, doublet-base-substitution and indel signatures

Single-base-substitution (SBS), doublet-base-substitution (DBS) and insertion and deletion (indel; ID) signatures were extracted *de novo* and related to known COSMIC signatures (v3.2) using SigProfilerExtractor ^78^. SBS, DBS and ID signatures were extracted using random initialization, 500 NMF replicates, and between 10,000 and 1,000,000 NMF iterations. We assumed the presence of between 1 and 30 SBS signatures (*minimum_signatures* and *maximum_signatures* parameters respectively), 1 and 15 DBS signatures, and 1 and 10 ID signatures. Default settings were used for all other parameters. Investigations of the novel DBS-A signature (**Supplementary Table 3**) hinted towards the signature being a technical artefact of the high number of short indels at homopolymer regions occurring in MSI samples.

#### 8.2. Characterising structural variant signatures

SV signatures were extracted considering only simple SVs, specifically deletions, tandem duplications, balanced and unbalanced inversions, and balanced and unbalanced inter-chromosomal translocations. Deletion and tandem duplication size distributions are multimodal (**Supplementary Fig. 6**), and we therefore classified these variants as ≤10kb, 10kb to 1Mb, and >1Mb. Variant site replication timing is also multimodal and we therefore classified variants as late, mid, or early replicating considering mean Repli-Seq thresholds of ≤-2, -2 to 2, and >2 (using CRC epithelial cell data from ReplicationDomain). Mechanisms of fragile site instability differ from other SVs and deletions and tandem duplications at fragile sites were therefore considered separately ^20^. Signatures were extracted using a hierarchical Dirichlet process (HDP) implemented in the hdp R package (v0.1.5) ^20^. The HDP structure was initialized with one common grandparent node, a parent node for each of the MSS, MSI and POL tumour subtypes, and a child node for each of the 1,765 tumour samples in which SVs were called. Four separate Markov chain Monte Carlo posterior sampling chains were run with 5,000 burn-in iterations, extracting 12 SV signatures. Extraction stability was assessed by splitting the cohort into halves, maintaining proportions of MSS, MSI and POL tumours, and re-extracting signatures from each half. Nine signatures extracted from the cohort halves showed high similarity between halves (cosine similarity >0.9; **Supplementary Fig. 6**) and high similarity with signatures extracted from the full cohort. These nine signatures were named SV1-9 and considered in subsequent analyses.

To investigate DNA repair mechanism perturbation, we correlated driver gene mutation with SV signature activity. A gene was considered mutated if it harboured a likely pathogenic germline SNP or indel (variants annotated as “Pathogenic” or “Likely Pathogenic” in ClinVar) ^79^, a likely oncogenic somatic SNV or indel, or a homozygous deletion at a gene exon. Pairwise associations between gene mutation and SV signature activity were tested for using multiple linear regression, including gene mutation status, age at sampling, primary tumour site and tumour sample purity as independent variables. Genes were considered if they were mutated in at least 1% of the tumour set. *TP53* mutation is associated with increased chromosomal instability, and *TP53* mutation status was therefore included in all models. The Yeo-Johnson extension to the Box-Cox-transformation was applied to mutation numbers to reduce heteroscedacity and ensure distributions were approximately normal ^80^. Samples with missing independent variable values were excluded. Due to mutational burden heterogeneity, only primary MSS tumours were considered in this analysis. *P*-values were adjusted for multiple testing using Bonferroni correction and a threshold of *P*=0.05 considered significant.

#### 8.3. Characterising copy number signatures

SigProfilerExtractor was used to extract copy number (CN) signatures in the 1,765 tumours with profiled CNAs ^81^. Where Battenberg identified a subclonal CNA, the copy number states corresponding to the largest tumour cell fraction were used, as SigProfilerExtractor cannot consider subclonal copy number states. Each copy number segment was assigned to one of 48 categories using SigProfilerMatrixGenerator, considering heterozygous or homozygous state, total copy number and segment length ^54,78^. Combinations of 1-30 de novo signatures were extracted and recommended solution was accepted, balancing Cosine distance with average stability. De novo CN signatures were then deconvoluted into their matching component COSMIC CN signatures from COSMIC v 3.

Each copy number signature was assigned as being active or inactive in each sample. Associations with MSI status, ploidy and HRD status were calculated via Fisher’s exact test, comparing samples with and without the phenotype with those that had or did not have the active signature.

#### 8.4. Predicting homologous recombination deficiency

Evidence of homologous recombination deficiency (HRD) was assessed using HRDetect ^72^. HRDetect considers six genomic features predictive of HRD: (1) proportion of deletions with microhomology, (2) SBS3 contribution, (3) SBS8 contribution, (4) rearrangement signature RS3 contribution, (5) rearrangement signature RS5 contribution, and (6) HRD index. HRDetect requires CNA data and was therefore run only on the 1765/2023 tumours passing CNA calling. SBS3 and SBS8 contribution estimates were obtained from SigProfiler. Rearrangement signatures RS3 and RS5 were computed using HRDetect, using rearrangement signature characterized by Nik-Zainal, et al. ^82^. Whilst HRDetect was trained on breast cancers, it has demonstrated high efficacy when applied to other cancer types ^72^. It was not possible to retrain HRDetect using our CRCs, as few tumours exhibited a pathogenic germline *BRCA1* or *BRCA2* variant with somatic loss of heterozygosity of the wild-type allele.

### 9.  Pathway analysis

#### 9.1. Analysis of disrupted pathways

Altered pathways were identified by integrating coding and non-coding mutations using ActivePathways ^83^. MSS, MSI and POL cancers were considered separately. 6 mutation features were used: coding driver P-values from IntOGen ^36^ and 3’ UTR, 5’ UTR, core promoter, distal promoter and non-canonical splice site P-values from OncodriveFML ^39^. We tested Reactome pathways obtained from MSigDB ^84^. All protein-coding genes included in at least one Reactome pathway were considered as the background gene set.

#### 9.2. Driver mutation co-occurrence

We used the DISCOVER algorithm ^85^ to investigate whether driver genes mutation, amplification or deletion showed evidence for pairwise co-occurrence or mutual exclusivity in primary MSS tumours. Simple methods such as Fisher’s exact test assume that the probability of a gene alteration is independent and identically distributed across genes^85^. However, as genes and tumours have heterogenous mutation rates, co-occurrence or mutual exclusivity relationships could be observed due to this confounder. DISCOVER accounts for this by adjusting for mutational heterogeneity at both the gene and patient level.

A gene was classified as altered if it was affected by either: (1) at least one non-synonymous SNV or indel, (2) a homozygous deletion or (3) a copy number gain (as defined above, conditional on tumour ploidy). For driver genes we included only pathogenic SNVs and indels, classified using OncoKB and dNdScv as described previously. For non-driver genes, for which mutations were included to assess tumour mutational burden, we considered SNVs and indels annotated as “moderate” or “high” impact in consequence classification from Ensembl v101.

Driver gene pairs were tested for mutual exclusivity and co-occurrence, on the condition that more than 20 mutational events occurred for a specific gene. *P*-values were adjusted for multiple testing using the step-down procedure which better controls the false discovery rate given discrete test statistics. For each relationship, the range of co-occurrences expected by chance, conditional on not achieving a *P*-value <0.001, was stated for both co-occurrence and mutual exclusivity. To ensure that relationships were not driven by heterogeneous mutation rates across colorectal sites, tumours were stratified by site (proximal colon, distal colon, and rectum) and all relationships re-assessed.

### 10. Immune profiling

#### 10.1. Human leukocyte antigen haplotyping

Human leukocyte antigen (HLA) typing of blood-derived normal samples was conducted using HLATyper, which is part of the Illumina Whole Genome Sequencing Service Informatic pipeline. The highest-ranking allele pair prediction for each type-I HLA allele (A, B and C) was taken to define a six-allele HLA set for each case.

#### 10.2. Immune escape prediction

We predicted three separate mechanisms of immune escape: (i) HLA gene mutation; (ii) HLA gene LOH; and (ii) mutation and LOH of other antigen presentation genes (APGs).

Somatic mutations in the HLA locus were predicted using POLYSOLVER ^86^. First, alleles were converted to a POLYSOLVER-compatible format (lower case, digits separated by underscore) and outputted into a patient-specific *winners.hla.txt* file. Next, the POLYSOLVER mutation-detection script (*shell_call_hla_mutations_from_type*) was run on matched tumour-normal pairs to call tumour-specific alterations in HLA-aligned sequencing reads using MuTect ^87^. Strelka (v2.9.9) ^88^ was also run to detect short insertions and deletions in HLA-aligned reads, as it offers increased sensitivity over POLYSOLVER’s default caller. Finally, both SNVs and indels passing quality control were annotated with POLYSOLVER’s annotation script (*shell_annotate_hla_mutations*).

LOH at the HLA locus was predicted using LOHHLA ^89^. The same *winners.hla.txt* files were used as input, with POLYSOLVER’s comprehensive deduplicated fasta of HLA haplotype sequences as reference. A type-I allele of a patient was annotated as “allelic imbalance” (AI) if the *P*-value corresponding to the difference in evidence for the two alleles was <0.01. Alleles with AI were further labelled as LOH if the following criteria held: (i) the predicted copy number of the lost allele was <0.50 with confidence interval <0.70; (ii) the copy number of the kept allele was >0.75; and (iii) the number of mismatched sites between alleles was >10.

We also evaluated somatic mutations and copy number status of the following APGs ^90^: *B2M, CALR, CANX, CIITA, ERAP1, ERAP2, HSPBP1, IRF1, PDIA3, PSMA7, PSME1, PSME2, PSME3*, *TAP1* and *TAP2*. First, somatic mutations were annotated using ANNOVAR ^91^. An APG was deemed mutated if it contained any non-synonymous, frameshift, stop-loss, or stop-gain mutation in its exons. The copy number status of each gene was evaluated using Battenberg output.

A sample was defined as immune escaped if it showed at least one of: (1) HLA mutation; (2) HLA LOH; or (3) APG mutation. HLA AI was not considered to provide immune escape as AI can arise from multiple sources (including subclonal LOH and unequal focal gains of the locus) and therefore the effect of AI on antigen presentation is uncertain. Where HLA alterations could not be fully evaluated (see Sample subsetting below), but no HLA or APG alteration was detected, the immune escape status was considered “Unknown”, since we could not eliminate the possibility of immune escape.

#### 10.3. Neoantigen prediction

We predicted neoantigens using NeoPredPipe; a Python-based pipeline combining ANNOVAR and netMHCpan (v4.0) ^91-93^. Briefly, all somatic SNVs and indels were annotated using ANNOVAR and for all non-synonymous exonic mutations the mutated peptide sequence was predicted. We took any 9- and 10-mer spanning the mutated amino acid(s), resulting in either (1) a 19-aa window for SNVs or (2) a peptide until the next predicted stop codon for frameshift mutations. These peptides were evaluated according to their novelty and predicted binding strength to the patient’s six-allele HLA set comprised of the HLA-A, -B and –C genes. Peptides novel compared to the healthy human proteome with binding rank of two or below (amongst the best 2% of binders compared to a set of random peptides) were reported as neoantigens. All patient-specific HLA alleles were used for neoantigen prediction, regardless of mutation or LOH status of the HLA locus.

We considered a mutation neoantigenic if at least one of its downstream mutated peptides was a neoantigen with respect to any of the patient’s six HLA alleles. We defined neoantigen burden as the total number of neoantigenic mutations in the sample. We also evaluated the following alternative measures: (1) the number of peptide-HLA binding pairs; (2) the number of strong binder (best 0.5% of peptides) peptide-HLA binding pairs; (3) the number of neoantigenic mutations in genes expressed in CRC (expression ≥10 TPM in ≥10% of TCGA CRCs) ^31^. We found that all these measures were very highly correlated with our definition of neoantigen burden: (1) R=0.993, (2) R=0.989, (3) 0.983; P<10^-16^ for all.

#### 10.4. Sample subsetting and statistical analysis

Eighty-five samples were excluded from neoantigen calling because netMHCpan was unable to predict at least one of their HLA haplotypes. 217 samples had one or more haplotypes incompatible with POLYSOLVER, for which HLA mutation and LOH calling was restricted to the compatible haplotypes (one, two and three haplotypes were excluded in 171, 37 and 9 samples respectively). In addition, LOH was not considered for 15 patients because they were homozygous for all type-I HLA genes. In total, 1,744/2,023 samples had complete neoantigen and HLA alteration information available.

As CRC subtypes (MSS, MSI and POL) have substantially different mutation and immune properties, all analysis were completed separately for each subtype. Pairwise comparisons were conducted using non-parametric two-sided Wilcoxon tests. Multivariate regression between immune escape types and neoantigen burden was performed using the *lm* function against the logarithm of neoantigen burden and therefore defined the fold change in burden associated with each escape type. The number of POL samples was insufficient for statistical analysis and the regression analyses were therefore only conducted for MSS and MSI tumours.

#### 10.5. Patient Harmonic Best Rank analysis

We computed the immunogenicity of a given mutation in a given patient using Patient Harmonic Best Rank (PHBR) ^94^ that takes into account all novel peptides produced by that mutation and all HLA alleles present in the patient. Low PHBR values correspond to mutations likely to be presented on the cell surface and hence with a high immunogenic potential while high PHBR mutations are less immunogenic. The overall immunogenic potential of a mutation within a cohort is defined as the median of PHBR values within that cohort. Briefly, for each mutation and HLA haplotype pair considered, we generated all 8-11-mers overlapping the mutation and evaluated their binding affinity to the HLA allele using the ‘all-predictions’ mode of NeoPredPipe. The best (lowest) rank was recorded. For a given patient, PHBR were computed as the harmonic mean of six best rank values corresponding to the patient’s six HLA haplotypes (homozygous alleles were counted twice). We computed PHBR values for all single nucleotide mutations located in driver genes that were present in at least four cancers in the cohort. The 85 samples with incompatible HLA alleles were excluded.

To evaluate the effect of HLA alterations on PHBR values, we repeated the same analysis for affected patients with a reduced set (<6) of HLA alleles that were un-altered. To measure the level of patient- (HLA-) dependent selection on driver genes, we compared PHBR values for mutations in these genes between patients did not carry the mutation and patients that did. Negative values indicate that mutations of the gene are enriched in patients where they have lower immunogenic potential. PHBR values between non-mutated and mutated patients were compared using Wilcoxon rank-test and p-values were adjusted for multiple testing using Benjamini-Hochberg correction.

Median PHBR across our whole CRC cohort positively correlated with mutation frequency, with the most common mutation, *BRAF* V600E, having a median PHBR of 4.9. This is in line with our previous observation that while *BRAF* V600E was the most recurrent present SNV in the cohort, it was not the most recurrently immunogenic one. We then focused on genes that were identified as a driver in a single subtype alone (e.g. *ASXL2*, a POL-specific driver). We found that drivers specific to metastases showed lower immunogenic potential (higher PHBR), with no mutations falling below a median PHBR of 1 that would indicate strong immunogenicity. The reverse analysis, comparing shared (identified in all subtypes) drivers in cohorts formed of each subtype revealed a significantly lower PHBR (higher immunogenicity) of these drivers in primary MSI samples. However, the mean difference in PHBR was small (0.06).

Since each mutation has a different immunogenic potential in each patient, we can also use PHBR to evaluate whether mutations are typically present in patients where they have a lower immunogenic potential – synonymous to mutations being absent/deleted from cancers where they are immunogenic – by comparing PHBR values between patients with and without a mutation. A higher PHBR (lower immunogenicity) for patients with mutations is indicative of HLA-based restriction on the driver mutations within the cohort.

We examined all driver mutations across the full CRC cohort and showed an increase in PHBR for mutation-carrying cancers. Next, we computed PHBR values for cancers carrying HLA alterations (HLA mutation or LOH), using only HLA alleles not affected by an alteration and compared to PHBR derived by the full HLA allele repertoire. We observed an increase in PHBR, suggesting that HLA alterations (even without further means of immune escape) can decrease the immunogenicity of driver mutations.

Finally, we evaluated the evidence of HLA-restricted mutation accumulation separately for all driver genes, by comparing PHBR values between wild-type and mutation-carrier cancers for mutations within each gene. A negative value of this measure is indicative of an immunogenicity-related selection on which patients will carry (at an observable frequency) a specific driver mutation. *BRAF, TP53, KRAS, SMAD4* and *PIK3CA* showed evidence of significant (Q<0.1) HLA-restriction, with *BRAF* showing the highest PHBR difference between mutated and non-mutated cancers.

### 11. Mitochondrial genome characterization

#### 11.1. Calling mitochondrial somatic single nucleotide variants and indels

Somatic mitochondrial SNVs and indels were called using Mutect2 v4.1.4.1 ^95^, with the light strand as reference based on the human mtDNA revised Cambridge reference sequence (rCRS). Somatic mitochondrial variants were excluded if they had:

- Low mapping quality score (<20).
- Low base quality score (<20).
- An alternative allele frequency <1%.
- Missing alternative reads in any stand direction.
- Location within hypermutated regions (302-316, 514-525 or 3106-3109).

Mutational distributions of SNVs, categorised by the six possible pyrimidine substitution classes, were constructed to analyse mutational processes. Distributions of substitutions on the D-loop, including and excluding variants between the two origins of replication ($\text{O}_{H}$ and Ori-b, between sites 16,197 and 191) were also analysed by substitution class ^96^. Pathogenic variants were identified using ClinVar ^79^, considering annotations where at least one submitter provided an “interpretation with assertion criteria and evidence”.

#### 11.2. Mitochondrial copy number estimation

Autosomal and mitochondrial genome coverage was computed using fastMitoCalc ^97^. Using estimated sample purity ($\rho$), tumour ploidy ($\varphi$) and mean coverage depth, tumour sample mitochondrial DNA copy number was estimated as per Gerstung, et al. ^55^:

$$Tumour sample mtDNA copy number=\frac{mtDNA mean coverage}{Autosomal DNA mean coverage}(\rho\varphi+2(1-\rho))$$

Mitochondrial copy number was estimated for only the 1,765/2,023 tumours that passed CNA calling and therefore have had purity and tumour ploidy estimates.

Linear regression was used to correlate mtDNA copy number with age at sampling, tumour stage, site of primary tumour, sex, and tumour purity. The Yeo-Johnson extension of the Box-Cox transformation was applied to mtDNA copy number. Linear regression was applied considering all tumours, and segregating MSS and MSI tumours. Regression results were adjusted for multiple testing using the Benjamini-Hochberg procedure.

#### 11.3. Selection of mitochondrial mutation and *POLG* Correlation

For the 13 mitochondrial protein-coding genes, selective pressure was quantified by calculating the respective dN/dS values using the dNdScv R-package (with non-mtDNA chromosomes removed from the reference genome) with default parameters ^38^. A global mitochondrial dN/dS value was also estimated, excluding *MT-ND6* due to a suspected replication bias. Results were adjusted for multiple testing using the Benjamini-Hochberg procedure. In addition, it was investigated whether *POLG* mutations resulted in altered mitochondria mutational burden compared to other tumours. Only the primary MSI cohort was analysed for this trait, as other sub-cohorts had too few tumours with non-synonymous *POLG* mutations.

### 12. Genomic impact of prior treatments

Whether individuals had received systemic treatment or colorectum-targeting radiotherapy prior to sampling was based on data from NHSD and PHE-NCRAS. For NHSD, records related to systematic treatment were obtained from the Admitted Patient Care and Outpatients tables using associated Office of Population Censuses and Surveys (OPCS)-4 codes. For PHE-NCRAS, records related to systemic treatment were obtained from the AV_TREATMENT table using the event description codes, and from the Systemic Anti-Cancer Therapy (SACT) table. For PHE-NCRAS, records related to radiotherapy were obtained from the AV_TREATMENT table using the event description codes, and from the National Radiotherapy Dataset (RTDS) table considering records associated with a CRC diagnosis.

In total, 315 participants received systemic treatment or radiotherapy prior to tumour sampling. 278 participants received systemic therapy prior to CRC sampling for sequencing, and information on the drugs administered was available for 182 of these participants. For 253 participants the systemic treatment was used to treat CRC, whilst for 25 participants it was used previously to treat another cancer. 94 participants received capecitabine, 23 received cetuximab, 93 received fluorouracil, 39 received irinotecan, 109 received oxaliplatin, 46 received steroids, and 28 received other drugs. 118 participants received colorectum-targeted radiotherapy prior to tumour sampling.

Associations between systemic treatment and colorectum-targeting radiotherapy prior to sampling with mutational signature activity were tested using multiple logistic regression. Prior treatment with radiotherapy, capecitabine, cetuximab, fluorouracil, irinotecan, oxaliplatin and steroids was included in the models as binary independent variants. Other treatments administered prior to sampling occurred in fewer than five individuals and were therefore not included in the models. One model was created for each of the 63 identified SBS, ID, DBS, and SV signatures, with signature presence encoded as a binary dependent variable based on whether any evidence of the signature was identified in each sample. 96 samples that received treatment prior to sampling, but for which the specific administered drugs were unknown, were not included. Both primary tumours and metastases were considered in these analyses. Treatment coefficient *P*-values were adjusted for multiple testing using Bonferroni correction and a threshold of *P*=0.05, considered significant. Treatment duration was measured as the time between the first and last treatment administration.

### 13. METASTASIS-SPECIFIC ANALYSES

Tumours were split between primary (n=1,354) and metastatic (n=105) MSS samples. Only MSS samples were included as there was just one MSI metastasis and no POL metastasis. Five primary tumours were matched to metastasis samples in this cohort, but for the purposes of the analysis all samples were treated as unmatched. To determine mutational burden, VCF files were filtered for PASS variants and the number of SNVs and indels summed. These were then divided by the total genome length (3088.27Mb). For the binned copy number analysis, the genome was first partitioned into 2,766 1Mb windows. For each sample, the absolute allele-specific copy number within each bin was recorded. If two copy number segments overlapped a bin, the copy number of the segment with the larger overlap was recorded. Copy numbers were then classified according to section 6.2. For each aberration type (Gain/LOH) the proportion of primary tumours with that aberration was compared to the proportion of metastatic samples with two-tailed Fisher’s exact tests. The difference between the proportions was then plotted as a trace along the genome with stars indicating significantly different bins. P-values were corrected for multiple testing (FDR<0.05). Absolute copy number calls were divided by mean integer ploidy to account for differences in ploidy between the two groups. The adjusted copy numbers for each bin were then compared between primaries and metastases via Wilcoxon signed-rank tests while correcting for multiple testing (FDR<0.05). The difference in the mean (ploidy-adjusted) copy number was then plotted as a trace along the genome, with stars indicating significant bins. The definition of immune escape was as described in section 11.2 and number of neoantigens determined according to the main metric in section 11.3.

### 14. Microbiome

#### 14.1. Microbial identification

Microbial sequences were identified using GATK PathSeq ^98^ aligned against the default PathSeq microbial genome bundles. A minimum clipped read length of 60bp was used with all other parameters set to their defaults. Unambiguously assigned reads were used for the decontamination steps. Thereafter the adjusted “score” output was used, which shares ambiguous reads between species. Score output for each sample was converted to microbial cells per human cell for each taxon by adjusting for microbial and human average genome size (average human genome calculated from copy number and tumour cell percentage data).

$$Microbial cells per human cell= \frac{{Microbial reads}/{Average microbial genome size}}{{Human reads}/{Average human genome size}}$$

This analysis showed that metastases had extremely low microbial content and therefore subsequent steps included only primary tumours unless otherwise stated. Reads passing PathSeq filters were realigned against the *E.coli* colibactin gene cluster ^99^ using bwa ^74^, and matching reads counted.

#### 14.2. Contaminants

Potential contaminant species were identified using methods developed by The Cancer Microbiome Atlas ^100^. Briefly, the prevalence of species found in primary tumours and matched blood was compared (**Supplementary Fig. 29**). Samples were called as positive for a species if two or more unambiguously aligned reads from the species was found. Species were deemed as likely tumour sample origin if a Fisher one-sided exact test found them to be more prevalent in the tumour sample than the blood sample (FDR<0.05) and blood sample prevalence was <20% of samples. Genus level scores were recalculated from species scores by only including the species scores that survived this decontamination step. To mitigate the effects of species with mixed biological and contaminant components ^101^ downstream steps were adjusted for NHS hospital Trust where possible (see below) as processing laboratory was a plausible source of contamination.

#### 14.3. Identifying taxa associated with colorectal cancer

CRC-associated taxa were identified by pooling all species level read numbers from eight published stool metagenomic studies ^102-107^. Application of LEfSe to these data identified 73 species and 37 genera associated with CRC ^108^. Bacterial species were classified as oral microbes if they were identified as “oral taxon” or “oral species” by PathSeq or if they were present in the expanded Human Oral Microbe Database (eHOMD) ^109^.

#### 14.4. Comparing microbiome to clinicopathological data

Microbial relative abundances were compared to clinicopathological data using decontaminated PathSeq output (**Supplementary Table 25**). Only tumours with complete data for the relevant categories were included in each comparison. Genus and species level alpha diversity was measured using the Shannon index and beta diversity using Bray-Curtis dissimilarity of relative abundance. Differences in beta diversity were measured by PERMANOVA using the adonis function ^110^ in Vegan using default settings, with permutations confined to within NHS Trusts using the “strata” setting to minimise cross-site contamination differences. Taxa differing between clinicopathological categories were measured using MaAsLin2 ^111^, with minimum abundance of 0, minimum prevalence of 0.1, and NHS Trust added as a random effect to minimise cross-site contamination differences.

### 15. Clinicopathological correlates

#### 15.1. Correlating variables

Multiple linear regression was used to investigate the relationship between clinicopathological features and numbers of SNVs, indels and SVs, and numbers of mutations attributed to SBS, ID, DBS, and SV signatures. Multiple logistic regression was similarly used to investigate the relationship between clinicopathological features and driver gene mutation, recurrent arm-level CNAs, recurrent focal CNAs, WGD, and evidence of CN signatures. Unlike SBS, ID, DBS and SV signatures, the activities of CN signatures do not represent numbers of mutations attributed to the signature^112^. We therefore considered only the presence and absence of CN signatures, not their activities.

Primary MSS and primary MSI tumours were considered separately. Signatures were tested if they were identified in at least 1% of the tumour set, driver genes were considered if they were mutated in at least 5% of the tumour set, and arm-level and focal copy number alterations were considered if identified as recurrent by GISTIC. *TP53* mutation is associated with increased chromosomal instability, and *TP53* somatic mutation status was therefore included in mutation number models. Considering multiple variables together in a single model is essential given that many of these variables are correlated, including age, primary tumour site, and stage ^24^. The Yeo-Johnson extension to the Box-Cox-transformation was applied to mutation numbers to reduce heteroscedacity and ensure distributions were approximately normal^80^. Samples with missing independent variable values were excluded. Primary tumour site and tumour stage were considered as ordinal variables. For each independent variable, *P*-values were adjusted for multiple testing using Bonferroni correction and a threshold of *P*=0.05 considered significant.

#### 15.2. Survival analysis

Correlation between clinicopathological and genomic variables with all-cause mortality was assessed using Cox proportional hazards models. Follow up time was measured from the date that the tumour was sampled to the corresponding patients most recent time of contact. The median follow-up time was 1075 days. Only individuals for which the primary tumour was sequenced were included. Due to proportional hazard assumption violation, individuals with MSS and MSI tumours were considered separately. Individuals were excluded if tumour sampling occurred prior to 1 January 2015 or the time between CRC diagnosis and tumour sampling was greater than one year. Hazard ratios were adjusted for sex, patent age at sampling (per 10 years), primary tumour site and Dukes stage. Due to small numbers of deaths, stage A and B categories were combined into a single category. Distal and rectum colon primary tumour site categories were also combined into a single category.

After excluding individuals with missing covariate data, the MSS and MSI cohorts comprised 836 (144 deaths) and 272 (48 deaths) individuals. The following variables were analysed:

- Total mutational burden (SNVs + Indels).
- SBS, DBS and ID mutational signature activity as binary indicators. Signatures were analysed if they were identified in <50% of tumours in the respective cohort.
- Immune escape status.

Where analyses required CNA profiles, smaller MSS and MSI cohorts comprising 810 (141 deaths) and 222 (40 deaths) individuals were used. The following variables were analysed using these smaller cohorts:

- Driver gene mutation status. Driver genes were considered mutated in a tumour if: (1) they contained an oncogenic mutation as defined by OncoKB and dNdScv annotation, (2) were homozygously deleted, or (3) were affected by a large copy number gain (total copy number state >5 for non-WGD tumours and total copy number state >10 copies for WGD tumours).
- WGD status.
- Chromosome-arm-level gains and deletions.
- Total SV number.

For each cohort, variables were only tested if at least 5% of deaths were present in each category. A variable was considered correlated with survival if it improved model fit via ANOVA and the z-test provided association evidence. The Benjamini-Hochberg procedure was used to adjust for multiple testing. Proportional hazards assumption violations were analysed for each test.

#### 15.3. Normal colorectal epithelial cell signatures

Numbers and proportions of SNVs associated with each SBS signature were obtained from Lee-Six, et al. ^113^. Where multiple crypts from the same colon region were sampled in a single individual, the median number and proportion of SNVs associated with each SBS signature was computed across these samples.

### 16. Code availability

**Supplementary Table 1** lists software versions used in this study and the URLs from which they were obtained.

### 17. Data availability

The 100,000 Genomes Project data can be accessed by joining the Colorectal Cancer Genomics England Clinical Interpretation Partnership (GeCIP) Domain (<https://www.genomicsengland.co.uk/portfolio/colorectal-cancer-gecip-domain>) using the following steps: (1) signing of the GeCIP Participant Agreement by your institution; (2) completion and submission of your personal GeCIP domain application (https://www.genomicsengland.co.uk/join-a-gecip-domain); (3) review of your application by the GeCIP Domain lead; (4) verification of your identify by your institution via direct communication with Genomics England; (5) signing of the GeCIP rules (https://www.genomicsengland. co.uk/wp-content/uploads/2019/07/GeCIP-Rules_29-08-2018.pdf) and completion of the Information Governance training. The Genomics England data access agreement can be obtained from https://figshare.com/articles/GenomicEnglandProtocol_pdf/4530893/5. All analysis of Genomics England data must take place within the Genomics England Research Environment (https://www.genomicsengland.co.uk/understanding-genomics/data). The 100,000 Genomes Project publication policies can be obtained from https://www. genomicsengland.co.uk/about-gecip/publications.
